# Neofunctionalized carbonic anhydrases in the biosynthesis of neuroactive plant alkaloids

**DOI:** 10.1101/2022.11.01.514683

**Authors:** Ryan S. Nett, Yaereen Dho, Chun Tsai, Daria Wonderlick, Yun-Yee Low, Elizabeth S. Sattely

## Abstract

Plants synthesize numerous alkaloids that mimic animal neurotransmitters. The diversity of alkaloid structures is achieved through the generation and tailoring of unique carbon scaffolds. However, many neuroactive alkaloids belong to a scaffold class for which no biosynthetic route or enzyme catalyst is known. By studying highly coordinated, tissue-specific gene expression in plants that produce neuroactive Lycopodium alkaloids, we identified a new enzyme class for alkaloid biosynthesis: neofunctionalized α-carbonic anhydrases (CAHs). We show that three CAH-like (CAL) enzymes are involved in a cryptic biosynthetic route to a key bicyclic precursor of the Lycopodium alkaloids, and additionally, we describe a series of oxidative tailoring steps that generate the optimized acetylcholinesterase inhibition activity of huperzine A. Our findings suggest a broader involvement of CAL enzymes in specialized metabolism and provide an example for how successive scaffold tailoring steps can drive potency against a natural protein target of interest.

## MAIN TEXT

The plant kingdom produces a large variety of compounds that affect cognition in animals.(*1*) These molecules likely act to protect against herbivory, and also make plants a rich source of therapeutics for treating neurological disease.(*2, 3*) Many of these neuroactive compounds are alkaloids, nitrogen-containing compounds derived predominantly from amino acids, which act by mimicking animal neurotransmitters.(*2*) Neuroactive alkaloids modulate the function of many different proteins involved in neuronal signaling, thereby causing alterations in behavior and cognition. These bioactivities have long been recognized, as alkaloid-rich plants have served as important botanical medicines for thousands of years, and many neuroactive alkaloids, such as the FDA-approved drugs morphine (analgesic), galantamine (dementia treatment), and atropine (muscarinic acetylcholine receptor antagonist), are still used in the clinic.(*4*)

The diversity in neuroactive alkaloid structures in plants is generated through complex biosynthetic mechanisms that convert primary building blocks (e.g. amino acids) into a variety of scaffolds that can be tailored to produce specific, bioactive end-products. Alkaloid scaffolds are typically generated by an enzymatic transformation that condenses two substrates to yield a novel, polycyclic structure.(*5*) However, unlike other major classes of plant natural products (e.g. terpenoids and polyketides), there is no single chemical theme or enzyme class that is implicated in alkaloid scaffold generation. For example, while multiple alkaloid families are generated via Pictet-Spengler condensations, the enzymes that catalyze these reactions belong to unrelated protein families that have convergently evolved.(*6*) Furthermore, many classes of alkaloids are derived through unique chemical transformations that have not previously been observed in plants. This is exemplified within the lysine-derived quinolizidine and Lycopodium alkaloids, which serve as the precursors for hundreds of potent, neuroactive compounds,(*7*) and whose scaffolds are thought to be constructed by unique, undefined chemical transformations for which no enzyme catalyst has been identified in nature.(*8, 9*) This inability to predict enzymes that build alkaloid scaffolds confounds the rapid elucidation of biosynthetic pathways in plants, and suggests that there are novel enzyme classes within plant metabolism yet to be identified.

Our interest in alkaloid scaffold biogenesis led us to focus on the Lycopodium alkaloids. These molecules are produced by plants within the Lycopodiaceae family (club mosses)(*9*) and consist of over 400 structurally-diverse, polycyclic alkaloids that have been studied as toxins and potential medicines.(*10, 11*) Perhaps the most well-known member of this alkaloid class is huperzine A (HupA, **17**),(*12*) an acetylcholine mimic that reversibly inhibits acetylcholinesterase (AChE) and which has been explored as a potential dementia treatment.(*13*) More broadly, the complexity and diversity of structures within the Lycopodium alkaloids has intrigued chemists for over a century,(*14*) and these compounds continue to be major targets for novel chemical synthesis strategies and isolation of novel structures.(*15*) However, while significant progress has been made in their total syntheses,(*16, 17*) the mechanisms that plants use to synthesize the many, diverse Lycopodium alkaloid scaffolds have remained largely unknown, and suggest the involvement of novel enzymes.

### Discovery of novel scaffold-generating enzymes

Prior isotope tracer studies (see **Fig S1** for summary) have demonstrated that the Lycopodium alkaloid scaffolds originate from two units each of a lysine-derived heterocycle (1-piperideine, **1**) and a polyketide substrate derived from malonyl-CoA (3-oxoglutaric acid, **2**, or its thioester analog).(*9*) These experiments enabled the recent identification of a biosynthetic route to 4-(2-piperidyl)acetoacetic acid (4PAA, **3**) and pelletierine (**4)**, the likely building blocks for all Lycopodium alkaloids (**Fig 1a**).(*18–21*) Specifically, we demonstrated that three enzymes from the HupA-producing club moss *Phlegmariurus tetrastichus* (lysine decarboxylase, *Pt*LDC; copper amine oxidase, *Pt*CAO; and piperidyl ketide synthase, *Pt*PIKS) are sufficient to convert the primary metabolites L-lysine and malonyl-CoA into **3**, which can spontaneously decarboxylate to yield **4** (**Fig 1a**).(*21*) While radio-isotope labelling studies with **4** have demonstrated this compound to be incorporated into downstream alkaloids, it was determined that this 8-carbon precursor is only incorporated into one “half” of 16-carbon Lycopodium alkaloid scaffolds (**Fig 1b & Fig S1**).(*22, 23*) In contrast, L-lysine, cadaverine, **1**, and **2**, which are presumed precursors to **4**, were all shown to be incorporated into both halves of this scaffold (**Fig S1**).(*22, 24–28*) These data suggest that a phlegmarine-type scaffold (**Fig 1b**) is formed through the pseudo-dimerization of a **4**-like molecule and a compound from which it is irreversibly derived, which has been proposed to be **3**, or an oxidized derivative.(*22, 23*)

**Figure 1.**
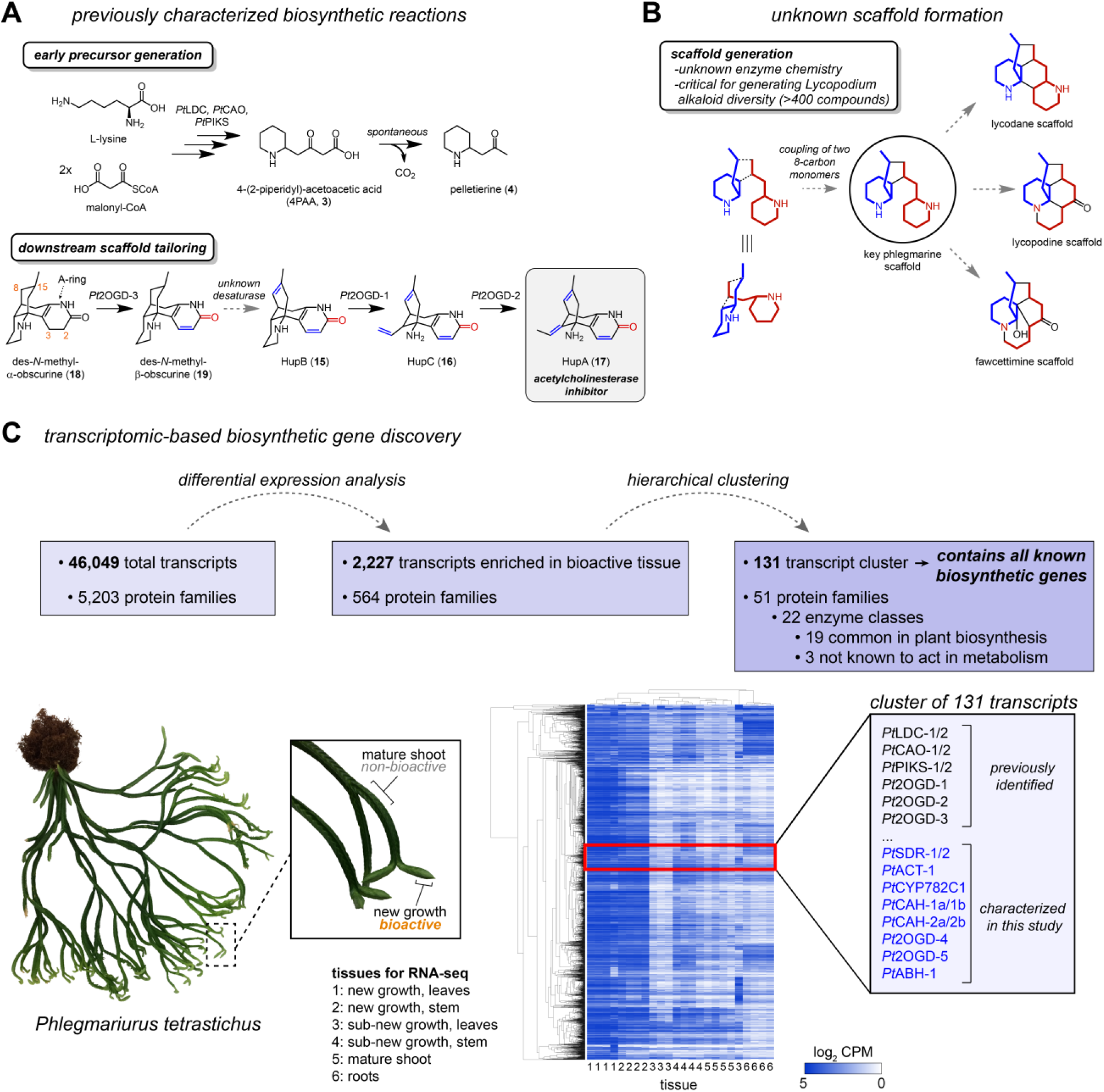
**A)** General biosynthetic scheme for HupA (**17**), along with previously identified enzymes. **B)** Summary of proposed scaffold formations for the Lycopodium alkaloids. **C)** Overview of experimental workflow for identifying new biosynthetic enzyme candidates.

While a condensation between **3** and **4** is plausible, it was unclear what type of enzyme could catalyze this type of reaction. Moreover, it was not clear if **3** and/or **4** needed to be further derivatized/processed prior to coupling. Because of this, we chose to rely heavily on the high level of biosynthetic gene co-expression that we had previously observed within our transcriptome of *P. tetrastichus*.(*21*) To leverage our previous results, we performed hierarchical clustering with these data to generate co-expressed clusters of transcripts (**Fig 1c**). This analysis revealed a single co-expressed cluster of 131 transcripts (cluster131) that contained all of the previously identified biosynthetic genes (*Pt*LDC-1, *Pt*LDC-2, *Pt*CAO-1, *Pt*CAO-2, *Pt*PIKS-1, *Pt*PIKS-2, *Pt*2OGD-1, *Pt*2OGD-2, and *Pt*2OGD-3).(*21*) This cluster was highly enriched with transcripts encoding for metabolic enzymes from multiple protein families commonly involved in natural product biosynthesis - e.g. cytochromes P450 (CYPs), Fe(II)/2-oxoglutarate dependent dioxygenases (2OGDs), methyltransferases, acyltransferases, and dehydrogenase/reductase enzymes - suggesting that it may contain the requisite biosynthetic machinery for Lycopodium alkaloid scaffold biosynthesis.

It had previously been proposed that a **4**-derived diene (**8**) could potentially serve as one of the co-substrates for scaffold formation.(*22*) We considered that the formation of this compound would require two key events: an oxidation of **4** to form the imine, and the reduction and elimination of the ketone oxygen. A related sequence of transformations had been reported in the context of morphine biosynthesis,(*29*) suggesting its plausibility in a plant system. To test candidate enzymes for this proposed route, we used *Agrobacterium* mediated DNA delivery in *Nicotiana benthamiana* as a transient gene expression platform. This allowed for production of **3** and **4** as substrates and the combinatorial testing of selected gene candidates (see **Methods** for details). We were unable to identify an oxidase that could act directly on **4**, so we instead prioritized dehydrogenase/reductase family enzymes within cluster131 that could potentially catalyze the ketone reduction. Only one short-chain dehydrogenase/reductase (SDR) family gene was found within this cluster (*Pt*SDR-1), and this had a close homolog (*Pt*SDR-2; 88.6% aa identity) that could be found in a slightly expanded co-expression cluster (273 transcripts; cluster273). When added to the transient expression experiments in *N. benthamiana* (containing *Pt*LDC, *Pt*CAO, and *Pt*PIKS), both SDR homologs led to modest consumption of **4**, and the detection of two new mass ions via LC-MS ([M+H]^+^ = *m/z* 144.1383) that correspond to reduction of the ketone to the alcohol (**Fig 2, Fig S2**). Comparison to a standard of 1-(piperidin-2-yl)propan-2-ol (i.e. reduced pelletierine) stereoisomers (**5**) confirmed these two new compound peaks to be diastereomers of **5**. Interestingly, *Pt*SDR-1 and *Pt*SDR-2 seemed to form different ratios of **5** stereoisomers. It had previously been noted that *Pt*PIKS produces racemic **4**,(*20*) which suggested that these SDR enzymes may utilize different enantiomers of **4** as substrate. Chiral LC-MS analysis confirmed the production of racemic **4** by *Pt*PIKS, and further demonstrated that *Pt*SDR-1 mainly consumed (*S*)-**4** to produce (*S, S*)-**5** (a.k.a (+)-sedridine), but also apparently acted on (*R*)-**4** to produce a small amount of (*R,S*)-**5** (a.k.a (+)-allosedridine) (**Fig S2**). *Pt*SDR-2 consumed both enantiomers of **4** to produce an equimolar amount of (*S, S*)-**5** and (*R, S*)-**5** (**Fig S2**), and appeared to be more active within our system.

**Figure 2.**
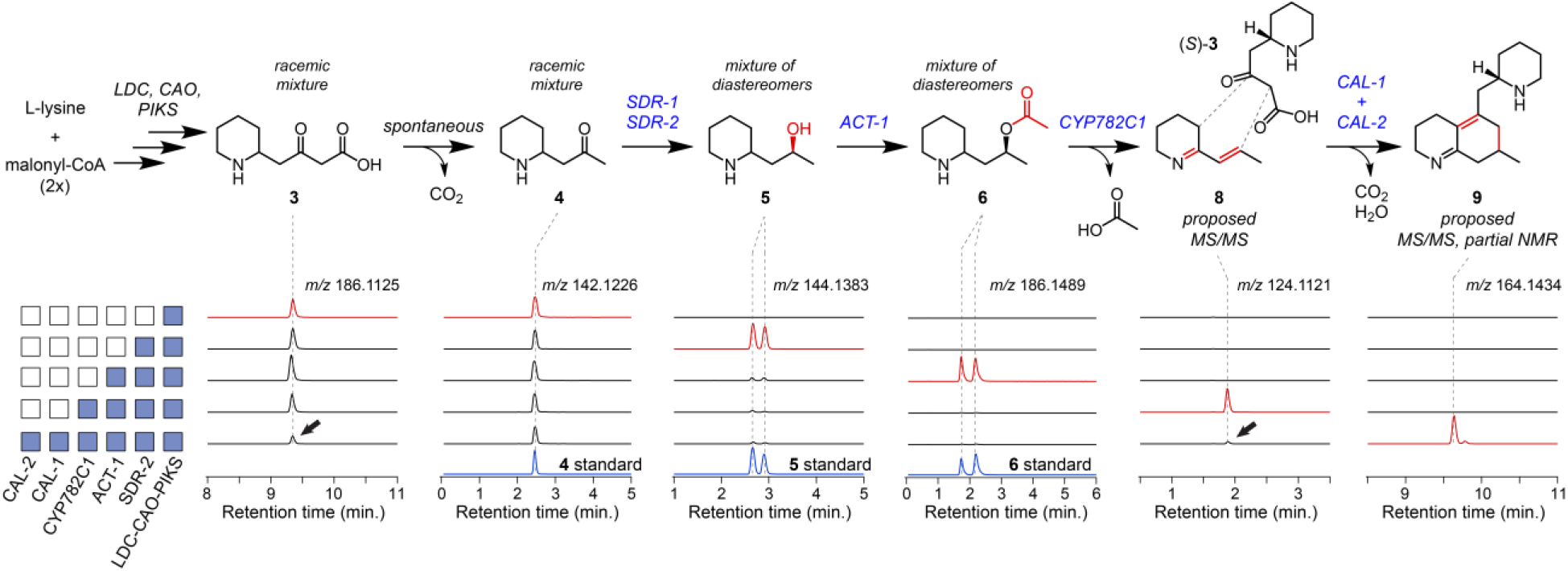
Stepwise discovery of early biosynthetic enzymes contributing to scaffold formation. Shown are the extract ion chromatograms (EICs) pertaining to the relevant *m/z* value for each defined or proposed intermediate. Note that y-axes for each set of chromatograms are on different scales, but scales are constant within an EIC plot. Black arrows indicate presumed consumption of substrates upon addition of CAL-1 and CAL-2.

Given the precedent for *O*-acylations in generating leaving groups for elimination in natural product biosynthesis,(*29, 30*) we next screened BAHD acyltransferase family enzymes for activity, as six unique gene sequences from this family could be found in cluster131. Addition of one acyltransferase (*Pt*ACT-1) to the transient co-expression system led to consumption of both (*S, S*)-**5** and (*R, S*)-**5**, and production of a new compound ([M+H]^+^ = *m/z* 186.1489) consistent with the addition of an *O*-acetyl group (**Fig 2, Fig S3**). Comparison to a synthesized standard and chiral LC-MS analysis confirmed this to be a mixture of two *O*-acetylated diastereomers, (*S, S*)-**6** and (*R, S*)-**6**, which shows this enzyme can *O*-acetylate regardless of the stereochemistry of the piperidine-alkyl bond (**Fig S3**).

The production of (*S, S*)-**6** and (*R, S*)-**6** was consistent with our hypothesis for elimination-mediated formation of the proposed diene (**8**). Because formation of **8** would require oxidation of the *O*-acetylated substrate(s), we next screened CYP and 2OGD family enzymes found within cluster131. One CYP enzyme (*Pt*CYP782C1) was found to consume both (*S, S*)-**6** and (*R, S*)-**6** within our transient expression system (**Fig 2, Fig S4**). This revealed the presence of two new compounds: one that corresponded to a single oxidation (desaturation) of the *O*-acetylated substrate (**7**, [M+H]^+^ = *m/z* 184.1332), as well as another that shared the same exact mass as **8** ([M+H]^+^ = *m/z* 124.1121). Both compounds were almost entirely lost if samples were incubated at room temperature for 1 hour (**Fig S4**), which is consistent with previous descriptions of **8**.(*31*) While this instability limited our ability to access authentic standards, MS/MS analysis supported the structures of **7** and **8**. This suggests a series of transformations in which the diastereomers of **6** are oxidized to produce **7**, which then undergoes an allylic elimination to yield **8** (**Fig S4**). Reactions of **6** (mixture of stereoisomers) with *Pt*CYP782C1 microsomes produced in yeast confirmed this activity (**Fig S4**), and importantly, allowed us to access **8** as an *in vitro*-generated substrate for any downstream enzymatic studies.

Production of **8** within our transient expression system suggested that we had potentially accessed the relevant substrates for initial scaffold formation (i.e. a phlegmarine-type scaffold, **Fig 1b**). However, it was difficult to select candidate enzymes given the lack of precedence for enzymes that could promote this type of chemistry; furthermore, it was uncertain exactly what “dimer” substrate combination should be tested. As a more untargeted approach, we opted to test candidates from cluster131 in batch combinations by enzyme family regardless of their prior association with specialized metabolism. In this process, we observed that a batch of four α-carbonic anhydrase (CAH) family proteins produced a new mass signature ([M+H]^+^ = *m/z* 164.1434) when transiently expressed in *N. benthamiana* leaves with the rest of the established pathway (**Fig 2, Fig S5**). The calculated molecular formula of this new feature (C_11_H_18_N) was unanticipated given the expected 8-carbon substrates, and this metabolite was only found using untargeted metabolomic analysis of our data with XCMS.(*32*) However, additional analysis of in-source MS adducts and fragments identified two co-eluting mass signatures that corresponded to the mass of a 16-carbon molecule ([M+H]^+^ = *m/z* 247.2169 and [M+2H]^2+^ = *m/z* 124.1121), suggesting that we had potentially accessed a scaffold from the condensation of two 8-carbon, nitrogen-containing substrates (**Fig S5**). With this in mind, the *m/z* 164 ion seemed to correspond to an ionization-induced loss of piperidine during mass spectrometry analysis (**Fig S5**). Subsequent MS/MS fragmentation of both the parent ion (*m/z* 247) and the in-source fragment (*m/z* 164) suggested that this new compound (mz247) possesses a phlegmarine-type scaffold (**Fig S5**). Moreover, mz247 could be detected within extracts from the biosynthetically active tissue of *P. tetrastichus* (**Fig S5**), which gave us confidence that this compound was relevant to Lycopodium alkaloid metabolism. We subsequently found that two of the batch-tested CAH-like (CAL) enzymes (named as *Pt*CAL-1a and *Pt*CAL-2a) were necessary and sufficient for this scaffold formation to occur (**Fig 2, Fig S5**), and that no apparent activity could be detected with either of the enzymes on their own. Additionally, we found that cluster131 contained homologs of *Pt*CAL-1a (*Pt*CAL-1b, 89.6% aa identity) and *Pt*CAL-2a (*Pt*CAL-2b, 70.1% aa identity) that exhibited the same activity (**Fig S5**).

We considered that this new compound (mz247) could be formed from a dimerization of **8**. However, when **8** was provided as a substrate to *Pt*CAL-1a and *Pt*CAL-2a independently of the full reconstituted pathway in *N. benthamiana* (**8** was generated by co-infiltrating **6** as a substrate for transiently-expressed *Pt*CYP782C1), formation of mz247 was not observed (**Fig S5**). Given this result, we considered that the scaffold may be a pseudo-dimer that requires the use of both **8** and another upstream pathway intermediate as a co-substrate. In support of this hypothesis, we could reconstitute production of mz247 in *N. benthamiana* through the combination of *Pt*CAL-1/*Pt*CAL-2a with a module for producing **8** (*Pt*CYP782C1 and synthetic **6** as substrate) and a module for producing **3** and **4** (*Pt*LDC, *Pt*CAO, and *Pt*PIKS) (**Fig S5**). Additionally, we observed consumption of both **8** and **3** that was concurrent with production of mz247, which supports these compounds as the coupling partners as the “dimers” that are condensed to form the scaffold molecule (**Fig 2, Fig S5**). Through chiral LC-MS, we further determined that only (*S*)-**3** was noticeably consumed in this reaction (**Fig S5**), which supports the specific condensation of (*S*)-**3** with **8** to form mz247. This is consistent with previously proposed mechanisms that implicate **3** as the nucleophile to initiate scaffold formation with an electrophilic co-substrate.(*22, 23*)

We scaled up production of mz247 in *N. benthamiana* for purification and structural confirmation of this compound, but this proved to be difficult, as it appeared to degrade over the course of purification. However, we were able to purify a putative oxidized product of this scaffold ([M+H]^+^ = *m/z* 263.2118) that accumulated during purification, and structural analysis of this molecule confirmed that it contained the predicted phlegmarine scaffold (see **Supplementary Materials** for structural characterization). Between the structure of this oxidized by-product, MS/MS fragmentation of mz247, and the chemical logic of a condensation between (*S*)-**3** and **8**, we predict that mz247 possesses the core phlegmarine scaffold with a conjugated α/β-unsaturated imine (compound **9**). Interestingly, a similar α/β-unsaturated imine moiety has been used within the chemical synthesis of Lycopodium alkaloid scaffolds,(*15*) where it was noted to be oxygen sensitive, and the NMR structure of our oxidized by-product (**9’**) is consistent with an oxidation of **9**. Thus, we propose that *Pt*CAL-1 and *Pt*CAL-2 homologs are jointly responsible for formation of **9** via the condensation of (*S*)-**3** and **8**, and that this serves as the key phlegmarine scaffold-forming reaction in Lycopodium alkaloid biosynthesis.

To our knowledge, CAH family proteins have not previously been shown to act directly in the biosynthesis of specialized metabolites. Therefore, this appears to represent a striking neofunctionalization within this enzyme class. Additionally, we were surprised by the apparent co-functionality of these two enzymes, since proteins from the CAH family usually function as monomers.(*33, 34*) To better understand the novel functionality of *Pt*CAL-1a/*Pt*CAL-2a, we next worked to establish an *in vitro* reaction assay. Both enzymes possess a predicted *N*-terminal signaling peptide that indicates trafficking through the secretory pathway, which suggested they may be localized to the apoplast (the extracellular compartment in plant leaves). To assess this, we expressed His-tagged versions of these proteins in *N. benthamiana*, and used Western blotting of different protein fractions (apoplast and cellular) to evaluate *Pt*CAL-1a and *Pt*CAL-2a localization. This confirmed that both enzymes are mainly present within the apoplast (**Fig 3a, Fig S6**), indicating that enzyme activity could be assessed in this protein fraction. Interestingly, we also observed that the presence of *Pt*CAL-2a was strongly affected by the co-expression of *Pt*CAL-1a. In particular, very little *Pt*CAL-2a protein could be detected when it was expressed alone, but upon co-expression of *Pt*CAL-1a, *Pt*CAL-2a was readily detected, and appeared to exhibit post-translational modifications of an unknown nature (**Fig 3a; Fig S6**). Beyond localization, this information was critical for enzyme assay development since the pH of the apoplast is typically relatively low (pH 4-5), which suggested that these proteins may only be functional at a lower pH.

**Figure 3.**
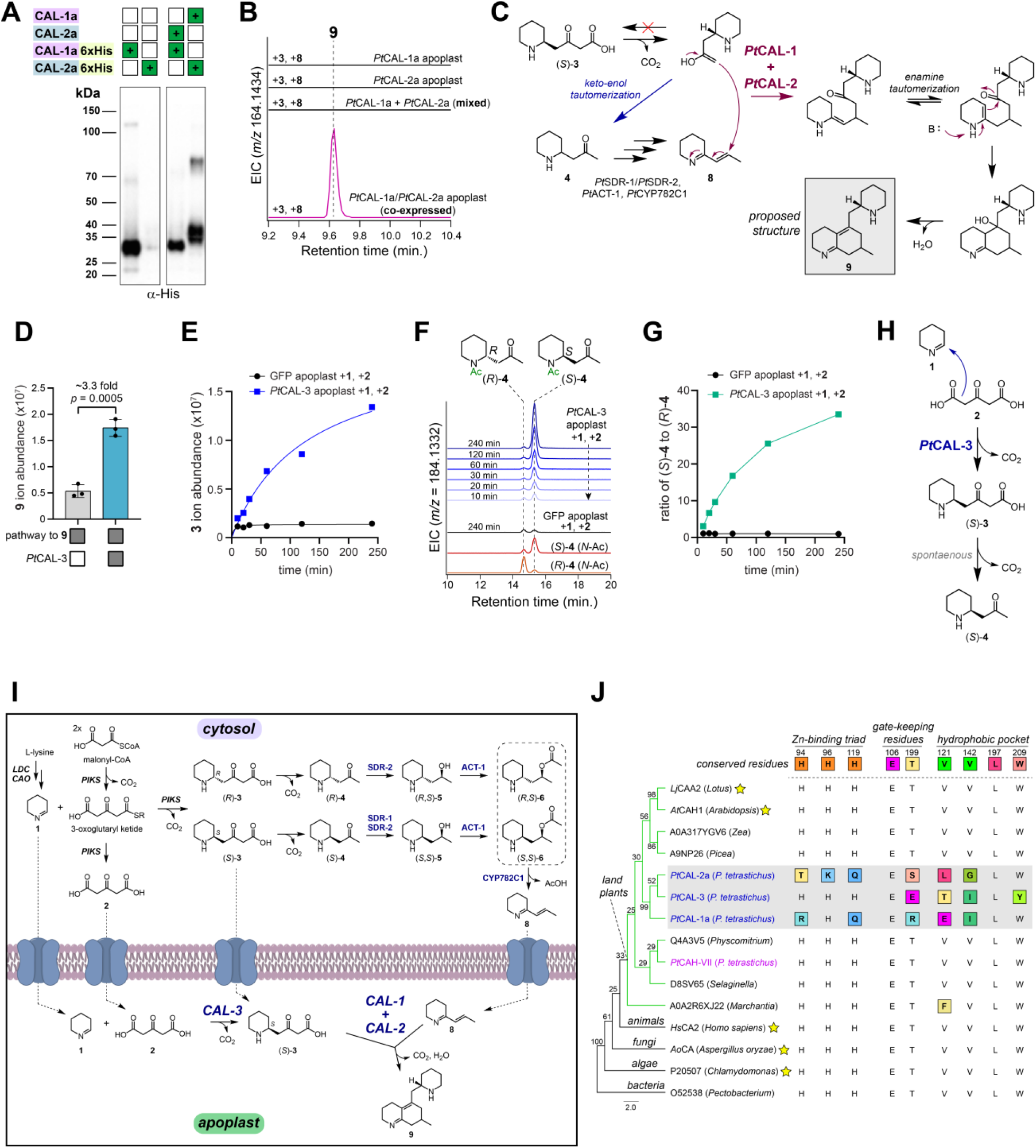
Characterization of multiple carbonic anhydrase-like (CAL) enzymes in Lycopodium alkaloid biosynthesis. **A)** Western blot of C-terminal His-tagged *Pt*CAL-1a and *Pt*CAL-2a. Enzymes were expressed alone, or in co-expression, within *N. benthamiana*, and shown here are blots of apoplast protein extracts. Note that these images are representative of a full blot shown in SI Figure S6. **B)** EICs for the production of a putative phlegmarine scaffold (**9**) by *N. benthamiana* apoplast extracts containing *Pt*CAL-1a and/or *Pt*CAL-2a when **8** and **3** are generated *in situ* as substrates. **C)** Proposed mechanism for the formation of a phlegmarine scaffold (**9**) by *Pt*CAL-1a and *Pt*CAL-2a. **D)** Effect of co-expressing *Pt*CAL-3 with the rest of the scaffold-generating pathway genes (*Pt*LDC-2, *Pt*CAO-1, *Pt*PIKS-1, *Pt*SDR-2, *Pt*ACT-1, *Pt*CYP782C1, *Pt*CAL-1a, and *Pt*CAL-2a). Shown are the ion counts (*m/z* 164.1434) for **9**. Statistical comparison was performed with Welch’s t-test. n = 3 for each condition. **E)** Formation of 4PAA (**3**) over time by apoplast extracts containing *Pt*CAL-3 when **1** and **2** are included as substrates, as measured by ion abundance of **3** (*m/z* 186.1125). **F)** Chiral LC-MS analysis of **4** enantiomers (after *N*-acetylation) that are generated during the apoplastic *Pt*CAL-3 reaction with **1** and **2** as substrates. Shown are EICs for the mass ion pertaining to *N*-acetylated **4** (*m/z* 184.1332). **G)** Ratio of (*S*)-**4** to (*R*)-**4** enantiomers (measured via chiral LC-MS analysis of *N*-acetylated derivatives) over time in the apoplastic *Pt*CAL-3 reaction. **H)** Proposed condensation of **1** and **2** catalyzed by *Pt*CAL-3 to produce (*S*)-**3. I)** Biosynthetic proposal for the early chemical transformations in Lycopodium alkaloid biosynthesis. **J)** Phylogenetic tree (MUSCLE alignment, Neighbor-joining tree) of CAH family proteins from multiple kingdoms of life. Bootstrap values (100 replicates) are located at nodes. Also shown are the major active site residues for each aligned protein. Changes to the canonical/conserved sequence are highlighted in colored boxes. Stars indicate proteins that have verified canonical carbonic anhydrase activity. An expanded alignment/phylogenetic tree can be found in Figure S8.

Using these apoplast extracts *in vitro*, we demonstrated activity (**Fig S6**) for *Pt*CAL-1a/*Pt*CAL-2a when **3** and **8** were supplied as substrates enzymatically (via the action of purified *Pt*PIKS +**1** and malonyl-CoA, and *Pt*CYP782C1 microsomes +**6** and NADPH, respectively). Notably, the enzymatic activity of *Pt*PIKS yields both **3** and **4** (through decarboxylation of **3**), and no production of **9** was observed for *Pt*CAL-1a/*Pt*CAL-2a when synthesized **4** was added as the cosubstrate with **8** (**Fig S6**). We further confirmed that the free acid of **3** (as opposed to a thioester conjugate) acts as the co-substrate, as **3** generated *in situ* (through spontaneous condensation of **1** and **2** substrates) could replace the *Pt*PIKS enzyme reaction within this *in vitro* system (**Fig 3b, Fig S6**). As observed with *in planta* experiments, only the (*S*) enantiomer of **3** was consumed in the presence of *Pt*CAL-1a/*Pt*CAL-2a, thereby supporting enantiospecific scaffold generation with this specific co-substrate (**Fig S6**). Consistent with *N. benthamiana* experiments and Western blot analysis, we observed maximal activity in apoplast extracts from leaves in which *Pt*CAL-1a and *Pt*CAL-2a were co-expressed; individual expression of each enzyme and subsequent mixing did not yield significant product formation (**Fig 3b**), suggesting that the presence of each protein within the cell is critical for their proper production and function. Although the exact mechanism by which these CAL enzymes function is not yet known, the condensation of (*S*)-**3** and **8** seems to resolve one of the outstanding questions related to the aforementioned isotope labeling studies in Lycopodium alkaloid metabolism, whereby condensation occurs between a substrate (**8**) that is irreversibly derived from its co-substrate (**3**) to yield a key scaffold (**Fig 3c**).

Encouraged by the identification of these neofunctionalized CAH enzymes, we considered that other transcriptionally co-regulated CAL genes might also have a role in this biosynthetic pathway. Upon testing another four CAL candidates with our established biosynthetic pathway via transient expression in *N. benthamiana*, we found a distinct CAL gene (*Pt*CAL-3) that led to a ∼3-fold increase in the abundance of **9** (**Fig 3d**). Analysis of all pathway intermediates that accumulated in this experiment revealed a shift in the abundance of **5** diastereomers to an enrichment of (*S,S*)-**5**, suggesting that *Pt*CAL-3 could be acting to influence the stereochemistry of precursor substrates (**Fig S7**). Including *Pt*CAL-3 with different combinations of pathway genes demonstrated that this enzyme is acting upstream of **4** formation, as we could observe a shift from racemic **4** to an enrichment of (*S*)-**4** (**Fig S7**). As with *Pt*CAL-1a and *Pt*PtCAL-2a, *Pt*CAL-3 contains an *N*-terminal signaling peptide and was found to be localized to the apoplast (**Fig S6**), and we were able to establish functional *in vitro* assays using apoplast extract from *N. benthamiana* leaves expressing this gene. In these assays, we demonstrated that including *Pt*CAL-3 apoplast with the PIKS reaction (using **1** and malonyl-CoA as substrates) led to an enrichment of (*S*)-**4** to (*R*)-**4** over time (**Fig S7**), and we speculated *Pt*CAL-3 protein may be accelerating the rate of **3** and/or **4** formation in a stereoselective manner. To decouple the activity of *Pt*CAL-3 from *Pt*PIKS, we tested **1** and **2** as substrates, which can spontaneously condense at a slow rate to form **3**. We observed a drastically accelerated increase in the formation of **3** compared to a control apoplast extract (**Fig 3e & Fig S7**), and determined that only the (*S*) enantiomer is enriched over time (**Fig 3f, 3g & Fig S7**). This demonstrates that *Pt*CAL-3 is acting to catalyze the condensation of **1** and **2** for the enantioselective biosynthesis of (*S*)-**3** (**Fig 3h**). In support of this, PIKS orthologs produce **2** as a major product,(*20, 21*) and thus the requisite substrates for *Pt*CAL-3 (**1** and **2**) are present from the activity of earlier enzymes in the pathway (*Pt*CAO and *Pt*PIKS).

Because *Pt*CAL-1a/*Pt*CAL-2a condense (*S*)-**3** with **8** to generate the core Lycopodium alkaloid scaffold, the specific production of (*S*)-**3** by *Pt*CAL-3 helps to explain the observed increase in **9** when *Pt*CAL-3 is present (**Fig 3d & Fig S6**). Furthermore, the presumed co-localization of these CAL enzymes within the apoplast provides a mechanism by which (*S*)-**3** can be directly utilized within scaffold formation without being fully consumed by the enzymatic steps that synthesize **8**, which are localized to the cytosol (**Fig 3i**). Our results also provide a rationale for the racemic production of **3**/**4** by the PIKS enzyme.(*20*) Since natural products are predominantly synthesized by enzymes in an optically-pure form,(*35*) it was initially quite surprising that PIKS would generate a racemic mixture of products. However, we have shown that during the course of biosynthesis, the stereocenter of **4**, as well those present in **5** and **6**, are lost upon formation of **8**. Thus, the corresponding stereoselectivity of metabolic enzymes (e.g. *Pt*SDR-2, *Pt*ACT-1, and *Pt*CYP782C1), may not have been strongly selected upon in the evolution of **8** biosynthesis, and the activity of *Pt*CAL-3 provides a bypass of these events to generate the stereospecific product needed for the scaffold generation.

To our knowledge, the identification of these three CAL enzymes represents the first known example of proteins from the CAH family participating directly within a metabolic pathway. Canonically, CAH enzymes catalyze the interconversion of CO_2_ and bicarbonate, and are critical for numerous cellular processes, including pH control, CO_2_ concentrating, and lipid metabolism.(*36*) The CAH family typically utilizes an extremely highly-conserved histidine triad to coordinate a Zn^2+^ cofactor, which generates the reactive hydroxide ion used to hydrate CO_2_. It is notable that in homologs of both *Pt*CAL-1 and *Pt*CAL-2, this histidine triad has been mutated (**Fig 3j, Fig S8**). In the case of *Pt*CAL-1, two of the three histidines are mutated, whereas all three are mutated in *Pt*CAL-2. Previous analysis of analogous mutations in CAHs have determined that perturbation of this triad leads to a loss in Zn^2+^-binding and carbonic anhydrase activity,(*37, 38*) and thus the mutations observed in *Pt*CAL-1 and *Pt*CAL-2 would seem to indicate a novel catalytic mechanism in these enzymes. While *Pt*CAL-3 retains this histidine triad, several other highly conserved active site residues involved in substrate binding (hydrophobic pocket) and proton shuttling have been altered (**Fig 3j, Fig S8**), presumably to accommodate the increase in substrate size relative to CO_2_/bicarbonate. More detailed work is needed to understand how these mutations may lead to CAH neofunctionalization, but these alterations in highly conserved CAH active site residues will provide a prominent basis for future mechanistic studies.

### Enzymatic tailoring for the production of neuroactive HupA

In our previous study of HupA (**17**) biosynthesis, we had identified three 2OGDs (*Pt*2OGD-1, *Pt*2OGD-2, and *Pt*2OGD-3) that function in the downstream tailoring reactions required to produce **17** from proposed precursors (**Fig 1**).(*21*) However, we were initially unable to identify an enzyme that could act on these substrates to form the 8,15-double bond (see **Fig 1a** for numbering) that is present in **17** and many other Lycopodium alkaloids, suggesting that we had not been testing the correct substrate(s). The simplest Lycopodium alkaloid with the same “lycodane” scaffold (**Fig 1b**) as **17** is flabellidine (**10**),(*39*) which contains an *N*-acetyl group on the A-ring nitrogen. Milligram quantities of this molecule had previously been purified,(*40*) which allowed us to test this as a substrate in *N. benthamiana* leaves expressing our oxidase gene candidates from cluster131 (CYPs and 2OGDs). Through this approach, we identified a pair of 2OGD enzymes that acted sequentially to convert **10** into downstream, oxidized products (see **Supplementary Materials** for additional details). The first of these enzymes (*Pt*2OGD-4) oxidized **10** to a molecule with an exact mass that is consistent with the installation of a carbonyl (proposed structure **11**, [M+H]^+^ = *m/z* 303.2067) (**Fig S9**), while the second enzyme (*Pt*2OGD-5) consumed **11** and produced a new, desaturated compound (proposed structure **13**, [M+H]^+^ = *m/z* 301.1911) (**Fig S10**). Although authentic standards were not available for these compounds, we suspected that *Pt*2OGD-4 was catalyzing formation the A-ring carbonyl, while *Pt*2OGD-5 was installing the 8,15-double bond.

If these were the oxidations, then the only remaining oxidation would be A-ring desaturation, which we have shown to be catalyzed by *Pt*2OGD-3.(*21*) However, *Pt*2OGD-3 did not consume **13**, and thus we hypothesized that *N*-deacetylation must precede this desaturation. Accordingly, we found an alpha/beta hydrolase family enzyme (*Pt*ABH-1) within cluster131 that consumed **13** to produce the *N*-deacetylated compound lycophlegmarinine D (**14**),(*41*) which verified the positioning of the carbonyl and double bond installed by *Pt*2OGD-4 and *Pt*2OGD-5, respectively (**Fig S11**). Addition of *Pt*2OGD-3 to the transiently co-expressed combination of *Pt*2OGD-4, *Pt*2OGD-5, and *Pt*ABH-1 led to the consumption of **14** and the formation of huperzine B (**15**) (**Fig S12**), and the subsequent addition of *Pt*2OGD-1 and *Pt*2OGD-2 allowed for the complete stepwise conversion to huperzine C (**16**), and ultimately, **17** (**Fig 4a, Fig S13**). While **17** has generated the most interest as a potential pharmaceutical,(*13*) hundreds of Lycopodium alkaloids have been isolated and structurally characterized,(*11*) including many congeners of **17** pathway intermediates that differ in their degree of unsaturation. Indeed, by mixing and matching enzymes from this downstream biosynthetic module, we were able to reconstitute the biosynthesis of 15 different Lycopodium alkaloids from **10** as an initial substrate, including many previously isolated and characterized compounds (**Fig 4a-d, Fig S13**; see **Supplementary Materials** for additional details). This demonstrates that the enzymes we identified contribute to a metabolic network of Lycopodium alkaloids within the endogenous plants, thereby explaining some of the notable structural diversity found among this class of alkaloid.

**Figure 4.**
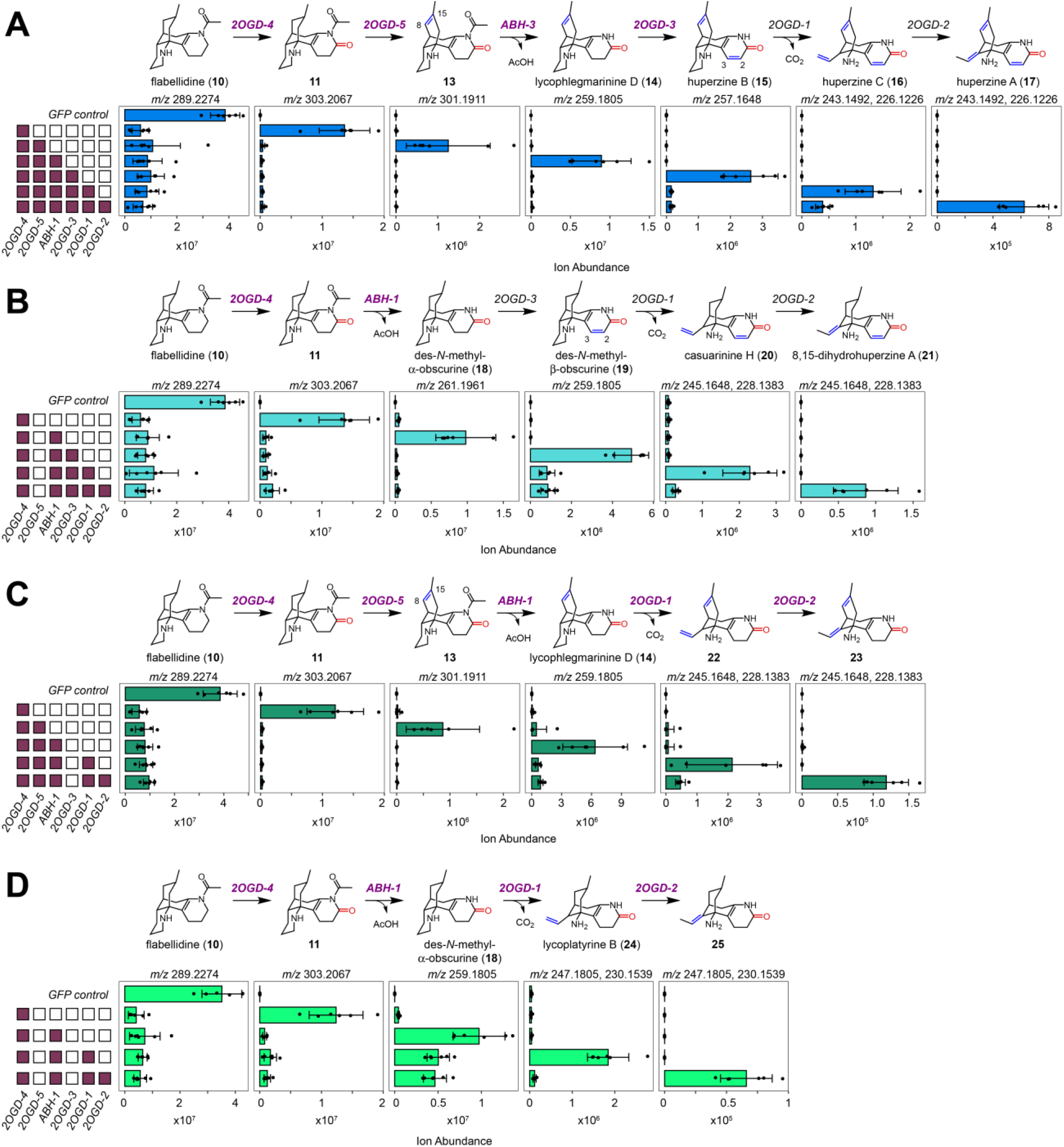
Step-by-step biosynthesis of downstream Lycopodium alkaloids. **A)** Generation of HupA (**17**). **B)** Generation of 8,15-dihydro congeners. **C)** Generation of 2,3-dihydro congeners. **D)** Generation of 2,3,8,15-tetrahydro congeners. For all panels, filled in boxes to the left indicate presence of biosynthetic genes within our *N. benthamiana* transient expression system. Shown below each compound is the ion abundance for the indicate mass ions (*m/z*) for each compound. n = 6 for experimental condition. New enzymes, or new reactions for previously described enzymes, are colored purple. Any Lycopodium alkaloids with common names have been verified with authentic standards.

The exact biological functions of Lycopodium alkaloids for the native plants have not been determined, but the ability of these compounds to inhibit AChE, a critical enzyme in animal neuronal signaling, suggests that they may act to deter herbivory via this mechanism. Indeed, AChE is a common target of insecticides,(*3*) and the high accumulation of Lycopodium alkaloids within the plant supports their role as toxic, defense molecules. It is interesting to note that **17** exhibits the most potent AChE inhibitory activity of any Lycopodium alkaloid measured thus far, and that this inhibition activity decreases with each prior intermediate in the pathway (**Fig 5**). This appears to represent a metabolic structure-activity relationship among the Lycopodium alkaloids, wherein each of the enzymatic transformations en route to produce **17** enhance AChE inhibitory activity. While we cannot be sure that the biological function of **17** is to inhibit animal AChE enzymes, the relationship between Lycopodium alkaloid biosynthesis and AChE inhibitory activity suggests that this metabolic pathway has evolved successive biosynthetic steps that increase the potency of these alkaloids step-by-step to achieve the production of an “optimized” AChE inhibitor.

**Figure 5.**
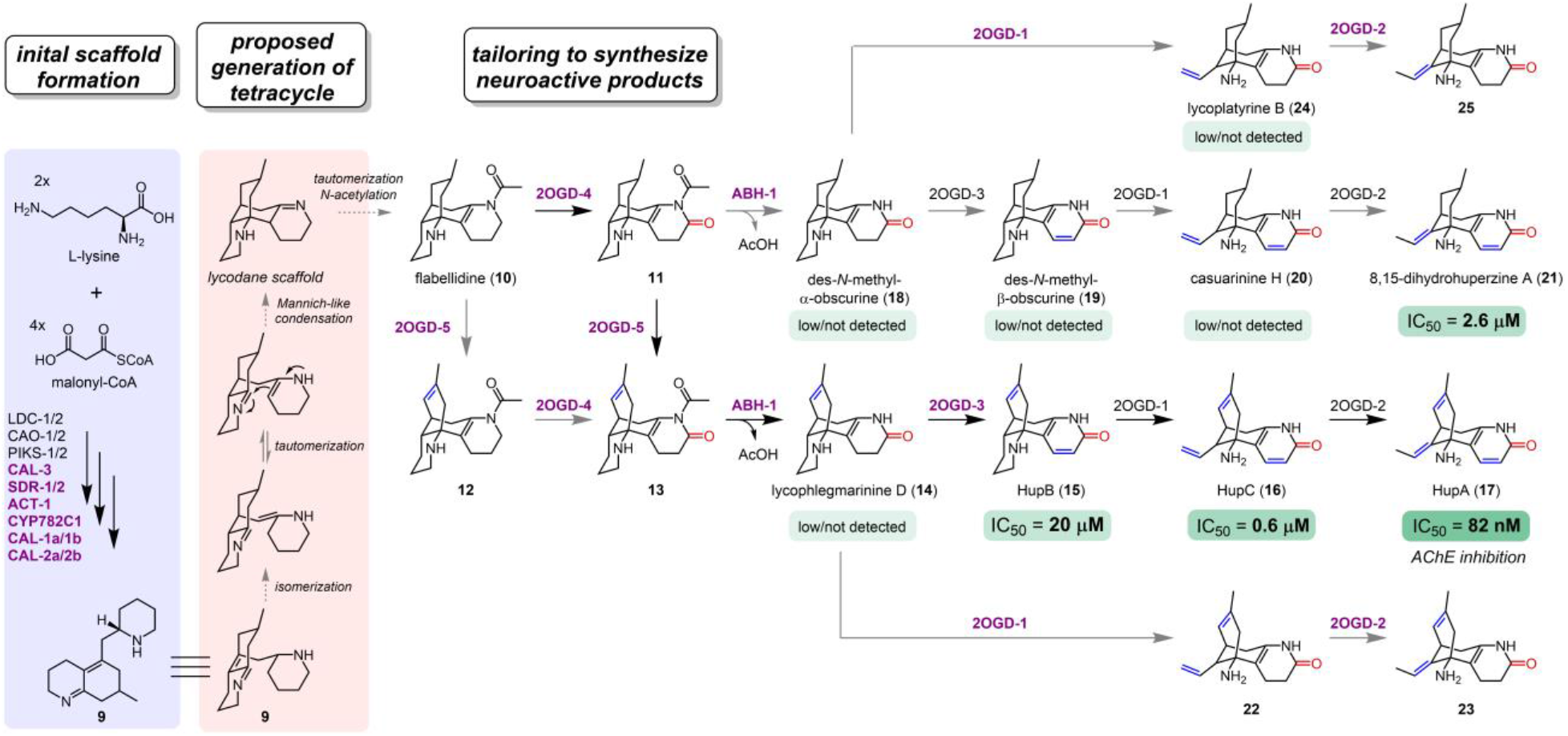
A metabolic network for the generation of an optimized AChE inhibitor, HupA (17). New enzymes, or new reactions for previously described enzymes, are colored purple. Any Lycopodium alkaloids with common names have been verified with authentic standards. Shown below Lycopodium alkaloids are their IC_50_ values for the inhibition of AChE, if previously tested. Citations for these values can be found in the Methods.

In support of our proposed biosynthetic pathway, all major biosynthetic intermediates from **4** to **9** and **10** to **17** could be detected within in extracts from the active site of **17** biosynthesis in *P. tetrastichus* (**Fig S14, Fig S15**). Transformation of the phlegmarine scaffold of **9** into the tetracyclic lycodane scaffold found in downstream alkaloids would putatively only require a double bond isomerization and enamine-imine condensation (**Fig 5**). Final *N*-acetylation of this scaffold on the A-ring would then yield **10**, thereby connecting upstream biosynthesis to the downstream transformations required to produce **17**. The identification of **10** as a precursor to **17** sheds critical light on the tentative chemical logical of this final tetracyclic scaffold formation. In particular, the addition of the *N*-acetyl group to the A-ring likely serves as a protecting group that “locks” the tetracyclic lycodane scaffold in place, which would otherwise be in equilibrium with the enamine/imine (**Fig 5**). In agreement with this premise, the *N*-acetyl group is only lost following formation of the A-ring lactam by *Pt*2OGD-4, which would also serve to deactivate the basicity of the nitrogen and protect the stability of this tetracyclic ring structure. While we do not know the nature of the enzyme(s) required to convert **9** into the theoretical enamine/imine intermediate, we can be confident that an acetyltransferase family enzyme is required for the final step to yield **10**.

Our efforts in identifying new enzymes in **17** biosynthesis (**Fig 5**) provide fundamental insight into the previously cryptic reactions used to build and tailor the scaffold structures of neuroactive Lycopodium alkaloids, and greatly expand our broader understanding of the enzymatic capabilities present within the plant kingdom. Most notably, our identification of multiple, neofunctionalized CAH family enzymes suggests that proteins from this family may have widespread roles throughout plant metabolism. Ultimately, our results place CAL proteins among a relatively short list of enzymes known in plants for the biosynthesis of alkaloid scaffolds.

## METHODS

### Chemicals and reagents

All common chemicals and reagents were obtained from commercial vendors. A mixture of 1- (piperidin-2-yl)propan-2-ol stereoisomers (**5**) was obtained commercially (MilliporeSigma). Authentic standards of (*S,S*)-**5** (a.k.a. (+)-sedridine) and (*S,R*)-**5** (a.k.a. (-)-allosedridine) were provided by Professor Paul Evans (University College Dublin). An authentic standard of lycophlegmarinine D (**14**) isolated from *Phlegmariurus phlegmaria*(*41*) was provided by Professor Ke Pan (China Pharmaceutical University) and 8,15-dihydrohuperzine (**21**) was provided by Professor Richmond Sarpong (University of California, Berkeley).(*15*) The following Lycopodium alkaloids were previously isolated from *Lycopodium platyrhizoma* (*40*): flabellidine (**10**), des-*N*-methyl-alpha-obscurine (**18**), des-*N*-methyl-beta-obscurine (**19**), casuarinine H (**20**), and lycoplatyrine B (**24**). Confirmatory NMR spectra for **10** and **20** can be found in the **Supplementary Materials**; those of **18** and **19** were previously reported.(*21*) The following Lycopodium alkaloids were purchased from commercial vendors: huperzine B (**15**, MilliporeSigma), huperzine C (**16**; two independent sources: Shanghai Tauto Biotech Co., Ltd and Toronto Research Chemicals Inc.), huperzine A (**17**, ApexBio Technology LLC).

### Transcriptomic and co-expression analysis

Transcriptomic data of *Phlegmariurus tetrastichus* was previously generated using PacBio IsoSeq for establishing a high-quality reference transcriptome of full-length sequences, and Illumina HiSeq 4000 for quantification of gene expression across many tissue types and biological samples.(*21*) Protein sequences encoded by each transcript were annotated with the best-hit Pfam term(*42*) using HMMER (http://hmmer.org/). Differential expression analysis was performed between samples from new growth leaves (biosynthetically active for HupA production) and mature shoot tissue (inactive for HupA production) using edgeR.(*43*) This analysis yielded 2227 unique transcripts that had significantly higher expression within the new growth leaves. These transcripts were then included in hierarchical clustering analysis using Cluster 3.0.(*44*) For this, expression counts (*TMM*-normalized, counts per million) for each transcript were normalized to the median expression value for that transcript, and these values were then log_2_-transformed. Transcripts were then hierarchically clustered using the Pearson correlation (centered) metric with average linkage, and visualized in TreeView software (https://jtreeview.sourceforge.net/). Relevant clusters were identified based upon the presence of previously characterized genes from Lycopodium alkaloid biosynthesis (*Pt*LDC-1, *Pt*LDC-2, *Pt*CAO-1, *Pt*CAO-2, *Pt*PIKS-1, *Pt*PIKS-2, *Pt*2OGD-1, *Pt*2OGD-2, and *Pt*2OGD-3). This allowed for the identification of a minimally-sized cluster of 131 transcripts that contained all previously characterized transcripts. Specific clusters of transcripts (cluster131 & cluster273) referenced herein are compiled in a separate **Supplementary Data** file.

### Agrobacterium-mediated transient expression

Candidate genes were cloned using cDNA from *P. tetrastichus* new growth leaves, much as previously described.(*21*) Following PCR amplification with primers containing appropriate overhangs, PCR products were gel purified and inserted into previously digested (AgeI/XhoI) pEAQ-HT plasmid (Kan^R^) using isothermal DNA assembly. Assembled plasmid reactions were transformed into *E. coli* NEB 10-beta cells (New England Biolabs) and plated on selective LB agar plates (50 µg/mL kanamycin) for overnight growth at 37 °C. Colonies were screened using PCR, and the sequences of PCR products were confirmed using Sanger sequencing. Positive transformants were then used to inoculate 4 mL liquid LB cultures, which were then shaken overnight at 37 °C. Plasmids were subsequently purified via mini-prep and inserts were again sequenced-verified using Sanger sequencing. Plasmids containing genes of interest were transformed into *Agrobacterium tumefaciens* GV3101 (Gent^R^) using the freeze-thaw method, plated onto selective LB agar plates (50 µg/mL kanamycin, 30 µg/mL gentamycin), and grown for 2 days at 30 °C. Positive transformants were verified via colony PCR, and these were then inoculated into 2 mL liquid LB cultures, which were shaken for two days at 30 °C. Colony PCR was again used to verify the presence of the plasmid construct in the liquid cultures, after which 25% glycerol stocks were prepared and stored at -80 °C for future use.

Screening of candidate genes via *Agrobacterium*-mediated transformation in *Nicotiana benthamiana* was performed much as previously described.(*21, 45*) *Agrobacterium* strains harboring plasmid constructs of interest were first thickly streaked from glycerol stocks onto LB agar plates (50 µg/mL kanamycin, 30 µg/mL gentamycin) and grown for 2 days at 30 °C. This lawn of cell growth was then removed using a sterile pipette tip, resuspended in 0.5 mL LB, and then pelleted via centrifugation at 8000 x *g* for 5 min. Cells were then resuspended in 0.5 mL of *Agrobacterium* induction media (10 mM MES, 10 mM MgCL_2_, 150 µM acetosyringone, pH 5.6) and allowed to incubate at room temperature for at least 1 hour. The concentrations of cell resuspensions were measured by taking their OD600, and combinations of strains of interest were then combined at a final OD600 of 0.2-0.3 for each strain. A needleless syringe was then used to infiltrate these strain mixtures into the abaxial side of *N. benthamiana* leaves from 4-5 week old plants, which were germinated and grown exactly as previously described.(*21, 45*) For a typical experiment, 3 leaves from 3 different plants were used for each strain mixture in order to minimize any batch effects or biological variation among plants. Following infiltration, plants were grown as usual for 3-5 days, after which leaves were excised for subsequent metabolite extraction. For substrate co-infiltration experiments, plants were grown for 3 days after *Agrobacterium* infiltration, after which 100 µL of substrate (25 µM in water) was infiltrated into the infected portion of the leaf using a needleless syringe. The area infiltrated with substrate was marked, and after one additional day of plant growth, this area was excised for subsequent metabolite analysis.

### Metabolite extraction

Following transient gene expression, *Agrobacterium-*infected leaf tissue was excised, placed in a pre-weighed 2 mL Safe-Lock tube (Eppendorf), and immediately snap frozen in liquid nitrogen. Typically, only one-quarter of a leaf was excised for analysis. When substrate was co-infiltrated, the entire marked area of infiltration was excised and snap frozen. Snap-frozen samples were either stored at -80 °C or immediately lyophilized to dryness. Following lyophilizing, samples were kept on ice or at 4 °C during all stages of processing. After removal from the lyophilizer, samples were weighed to collect dry masses. A 5 mm diameter steel bead was then added to each sample tube, and plant tissue was homogenized to a powder by shaking at 25 Hz for 2 min on a ball mill homogenizer (Retsch MM 400). Steel beads were removed with tweezers and homogenized tissue was extracted with an appropriate volume of solvent. For routine extraction 80% methanol (MeOH) in water was added at an amount of 20 µL solvent per milligram of dry leaf weight, and after mixing, samples were incubated on ice for at least 20 min. During the course of our experiments, we noted that certain intermediates (e.g. **3, 7**, and **8**) would be depleted over time, either due to decomposition or reactivity with other metabolites. We found that extracting samples with ice-cold water +0.1% (v/v) formic acid would improve the stability of these compounds without any major losses in alkaloid yield. As such, most of the LC-MS chromatograms that are shown for early pathway intermediates were derived from experiments in which water +0.1% formic acid was used as the extraction solvent.

After incubation, samples were briefly vortexed, and cell debris was pelleted via centrifugation at 10,000 x *g* and 4 °C for 5 min. After centrifugation, samples were prepared differently based upon the type of chromatographic analysis that was to be used (e.g. C18 vs HILIC). Samples related to the analysis of the early biosynthetic pathway (i.e. any of the products generated by *Pt*LDC-1/2, *Pt*CAO-1/2, *Pt*PIKS-1/2, *Pt*SDR-1/2, *Pt*CYP782C1, *Pt*CAL-1/*Pt*CAL-2, and *Pt*CAL-3) were diluted 10-fold in ice-cold acetonitrile (ACN) to better match the starting solvent conditions for HILIC analysis. Samples related to the analysis of downstream intermediates (i.e. any intermediates downstream of **10**) were diluted 1:1 with water +0.1% formic acid. All samples were then filtered through Multiscreen Solvinert filter plates (MilliporeSigma, Hydrophilic PTFE, 0.45 µm pore size) and subsequently transferred into LC-MS vials, which were stored at -20 °C or -80 °C until analysis.

### Preparation of metabolites for chiral analysis

Many of the early intermediates could only be observed by HILIC analysis, which made it difficult to resolve enantiomers with standard chiral chromatography. Protection of the secondary amines of **4** and its pathway derivatives via *N*-acetylation allowed us to readily separate enantiomers (**Fig S2**). The *N*-acetylation of standards, plant extracts, and enzyme reactions was performed as follows. A 10 µL aliquot of sample was diluted into 90 µL ACN (for standards, 10 µL of a 10 mM stock solution in MeOH was used), and 200 µL of acetic anhydride was then added. Samples were then heated at 60 °C for 30 min, though we noted that heating was not strictly necessary for *N*-acetylation to readily occur. After this incubation, samples were moved onto ice for at least 5 min, after which 300 µL of MeOH was added to quench the reaction. Quenched samples were then filtered and transferred into LC-MS vials, as described above. Standards were subsequently diluted to a concentration of 10-20 µM in 80% MeOH prior to analysis.

### LC-MS analysis

Samples were routinely analyzed on two different LC-MS instrument setups: **1)** an Agilent 1260 high-performance liquid chromatography (HPLC) instrument paired with an Agilent 6520 Accurate-Mass quadrupole time-of-flight (Q-TOF) mass spectrometer (6520 LC-MS), or **2)** an Agilent 1290 Infinity II UHPLC paired with a coupled Agilent 6546 Q-TOF mass spectrometer (6546 LC-MS). For both instruments, all samples were analyzed using electrospray ionization (ESI) in positive ionization mode. Reversed-phase (C18) analysis was predominantly performed on the 6546 LC-MS using a ZORBAX RRHD Eclipse Plus C18 column (Agilent, 1.8 μm, 2.1 × 50 mm) with water +0.1% formic acid and CAN +0.1% formic acid as mobile phases. HILIC analysis was predominantly performed on the 6520 LC-MS using a Poroshell 120 HILIC-Z column (Agilent, 2.7 μm, 2.1 × 100 mm) with water and 9:1 ACN:water, each with 0.1% formic acid and 10 mM ammonium formate, as mobile phases. Chiral chromatography was performed on the 6520 LC-MS using a CHIRALPAK® IC-3 column (Daicel, 3 μm, 4.6 × 100 mm) with water +0.1% formic acid and ACN +0.1% formic acid as mobile phases. Specific LC-MS method parameters can be found in the **Supplementary Materials**. When applicable, mass ions pertaining to individual metabolites were fragmented using targeted MS/MS. This was normally performed with multiple collision energies (10V, 20V, and 40V), but most of the presented data was collected with a collision energy of 20V.

LC-MS data was routinely visualized and analyzed using MassHunter Qualitative analysis. Quantification of relative ion abundance was performed using MassHunter Quantitative analysis. For untargeted analysis, data files were converted into mzML format, and XCMS software(*32*) was used to identify any differentially-produced mass ions between different gene expression conditions/reactions. This output was typically filtered to remove low abundance ions (less than 1×10^5^ ion abundance) and any ions that were not clearly differential between treatments (*p*-value > 0.2). XCMS analysis was typcially followed with CAMERA software(*46*) analysis to identify potential in-source ion adducts of detected metabolites.

### Apoplast protein isolation

The three CAL enzymes identified in this study have predicted N-terminal signal peptides, which were identified using the TargetP-2.0 server (https://services.healthtech.dtu.dk/service.php?TargetP-2.0).(*47*) Preliminary confocal microscopy of GFP-tagged proteins did not support their major localization to be the ER or Golgi, and initial analysis of images suggested that they may be localized to the apoplast. To assess this possibility, CAL genes with or without C-terminal 6x His tags were transiently expressed in *N. benthamiana*, as described above. Each CAL gene was expressed individually; additionally, *Pt*CAL-1a and *Pt*CAL-2a were transiently co-expressed in the same leaf since we had found them to co-function with Lycopodium alkaloid biosynthesis. At 4 days post *Agrobacterium* infiltration, apoplast protein extracts were isolated using the infiltration-centrifugation method, much as previously described.^24^ Two leaves per reaction were excised from the plant and submerged in ice-cold apoplast extraction buffer (100 mM MES, 300 mM NaCl, pH 5.5) within an open-capped 50 mL Falcon tube, and these tubes were placed in a plastic vacuum chamber attached to a Welch Model 2025 vacuum pump. The chamber was brought down to full vacuum, and after two minutes at this pressure, the vacuum was slowly released to allow for buffer to infiltrate the leaf apoplastic space. Buffer-infiltrated leaves were carefully removed from the Falcon tubes, blotted dry with paper towels, and were then rolled into Parafilm and placed in a plunger-less 5 mL plastic syringe. The syringe was placed in a 15 mL Falcon tube, and this was then centrifuged at 1000 x *g* and 4 °C for 10 min to collect apoplast extract. The resulting extract was then centrifuged at 10,000 x *g* and 4 °C for 15 min to pellet any larger cellular debris, and the supernatant was concentrated using an Amicon® Ultra-4 Centrifugal Filter Unit (10 kDA MWCO, MilliporeSigma UFC501024). Protein concentrations were measured using the BIO-RAD Protein Assay or Bradford assay (Abcam 119216) and adjusted with apoplast extraction buffer to a final concentration between 0.5-1.5 mg/mL. Aliquots of the extracts were then snap frozen in liquid nitrogen and stored at -80 °C.

### Western blot analysis of plant extracts

To determine localization of CAL proteins within our *N. benthamiana* transient expression system, we performed Western blot analysis of epitope tagged versions of each protein. Each CAL gene was PCR amplified from previously generated plasmid constructs using primers with overhangs for subsequent isothermal assembly into pEAQ-HT plasmid digested at the AgeI/XmaI restriction sites, which creates constructs with a C-terminal 6xHis tag. The reverse primer in this cloning strategy omitted the native stop codon of the CAL coding sequences to ensure that the final coding sequence included the C-terminal tag. These constructs were sequenced verified, transformed into *Agrobacterium tumefaciens* GV3101, and these strains were then used to transiently express these genes in *N. benthamiana*, as described above.

For the analysis of different protein fractions, apoplast extracts were prepared exactly as described above. Once apoplast extracts were obtained, the remaining leaf tissue was flash-frozen in liquid nitrogen and lyophilized to dryness. Lyophilized leaf tissue was pulverized to a powder with 5 mm stainless steel beads in a ball mill homogenizer (Retsch MM400) at 25 Hz for 2 min. Protein from homogenized samples was then extracted with ice-cold phosphate-buffered saline (PBS) supplemented with Halt protease and phosphatase inhibitor cocktail (Thermo Scientific PI78443) using 20 µL buffer per mg dry leaf mass. This was incubated on ice for 20 min with periodic, gentle inversion, after which samples were centrifuged at 18,210 x *g* for 10 min at 4 °C to remove insoluble plant material. The remaining supernatant was kept, and represented the “internal” cell fraction, which would presumably contain cytosolic and microsomal proteins. Protein concentration was determined by Bradford assay (Abcam 119216), and extracts were stored at -80 °C until future use.

Samples for immunoblots were prepared by adding 4X NuPAGE LDS sample buffer (Fisher Scientific AAJ61894AC) to a final concentration of 1X sample buffer with 2.5% β-mercaptoethanol, and samples were then heated for 20 min at 70 °C. Total protein for apoplast (2.5 µg) and PBS extracts (5 μg) was separated on NuPAGE gels, and then transferred onto a PVDF membrane (BIO-RAD 1704272) using a Trans-Blot semi-dry transfer system (BIO-RAD). Blots were blocked in EveryBlot blocking buffer (BIO-RAD 12010020) for >5 min at room temperature and incubated with mouse anti-His antibody (Genscript A00186) at 0.1 µg/ml in EveryBlot buffer for 1 hr at room temperature or overnight at 4 °C. After washing three times with PBST (PBS +0.1% Tween), blots were incubated with horse anti-mouse IgG, HRP-linked antibody (Cell signaling Technology 7076) at 1:3000 dilution. Blots were then washed five times with PBST and imaged with an iBright FL1500 Imaging System (Thermo Fisher Scientific).

### *Heterologous expression of* CYP782C1 *in yeast, microsomal protein preparation, and in vitro enzyme assays*

Expression of *Pt*CYP782C1 in *Saccharomyces cerevisiae* (yeast) was performed as previously described.(*45, 48*) Briefly, the coding sequence of *Pt*CYP782C1 was PCR amplified and annealed into the pYeDP60 plasmid. This plasmid construct was transformed into *S. cerevisiae* WAT11 (*ade2*) and positive transformants were selected on synthetic drop-out medium plates lacking adenine (6.7 g/l yeast nitrogen base without amino acids, 20 g/L glucose, 2 g/L drop-out mix minus adenine, 20 g/L agar) via growth at 30 °C for two days. Presence of the plasmid constructs were confirmed via colony PCR. A single, positive colony was used to inoculate a starter 4 mL culture of liquid drop-out medium, which was grown at 28 °C and 250 RPM. Following two days of growth, 2 mL of the starter culture was used to inoculate 500 mL of YPGE medium (10 g/L Bacto yeast extract, 10 g/L Bacto peptone, 5 g/L glucose and 3% (v/v) ethanol). This culture was grown at 28 °C and 250 RPM until reaching a cell density of 5×10^7^ cells/mL, which was estimated via OD600 measurements. After reaching this density, expression was induced by adding 50 mL of a sterile galactose solution (200 g/L) to achieve a concentration of approximately 10% (v/v). The culture was then grown at 28 °C and 250 RPM for another 16 hours to achieve a cell density of approximately 5×10^8^ cells/mL, after which this culture was immediately used for microsomal protein isolation, which was performed exactly as previously described.(*48*) Microsomal protein was stored in TEG buffer (50 mM Tris-HCl, 1 mM EDTA, 20% [v/v] glycerol, pH 7.4), aliquoted into 1.5 mL microfuge tubes, snap frozen in liquid nitrogen, and stored at -80 °C.

Enzyme reactions with *Pt*CYP782C1-enriched microsomal protein were performed in potassium phosphate buffer (50 mM potassium phosphate, 100 mM sodium chloride, pH 7.8) and typically contained 4 µg of microsomal protein (final concentration of 0.02 µg/µL), 500 µM NADPH, and 50 µM of **5** substrate in a total reaction volume of 200 µL. Control reactions omitted NADPH. Following addition of all components, reactions were incubated at room temperature for a minimum of 10 min. At specific time points, 20 µL aliquots of the reaction were added to 180 µL of ACN +0.1% formic acid to quench the reaction. Quenched reactions were then filtered and transferred into LC-MS vials, as previously described. Products of *Pt*CYP782C1 activity on **5** were assessed on LC-MS using HILIC analysis.

### In vitro enzyme reactions with apoplastic CAL protein

Reactions with CAL-enriched apoplast were routinely performed in low pH potassium phosphate buffer (50 mM potassium phosphate, 100 mM NaCl, pH 5.9) at a volume of 20 µL. For *Pt*CAL-1a/*Pt*CAL-2a, these reactions contained approximately 1 µg of apoplast protein for each CAL (final concentration of 0.02 µg/µL). Control reactions used apoplast from leaves expressing only one CAL, or with apoplast generated from GFP-expression *N. benthamiana* leaves. The requisite substrates for this reaction were generated through the activities of *in vitro Pt*PIKS-1 and *Pt*CYP782C1 enzyme reactions. We found that the *Pt*CYP782C1 microsomal reaction did not work well at the lower pH (pH 5-6) at which the CAL enzymes seemed to be most active (**Fig S4**). Therefore, prior to the CAL reactions, we ran a separate *Pt*CYP782C1 microsomal protein assay in high pH buffer (50 mM potassium phosphate, 100 mM sodium chloride, pH 7.8), much as described above, for a minimum of 2 hr to generate sufficient **8** as a substrate. To maximize the amount of **8** produced, substrate-generating *Pt*CYP782C1 reactions (100 µL) contained 1.5 mM of substrate (**6**), 10 µg of *Pt*CYP782C1 microsomes (final concentration of 0.1 µg/µL), and 4 mM NADPH. After these incubations, a 2 µL aliquot of the *Pt*CYP782C1 reaction (now containing **8**) was added to the *Pt*CAL-1a/*Pt*CAL-2a apoplast enzyme assay setup (20 µL total reaction volume). To generate **3** and **4** as potential substrates, 1 µg of previously purified *Pt*PIKS-1 enzyme(*21*) and 150 µM **1** and 300 µM malonyl-CoA were added directly to the CAL reaction mixtures. After thorough mixing, reactions were incubated at room temperature. An alternative route for producing **3** and **4** independently of thioester intermediates was achieved by mixing stocks of **1** (10 mM in water) and **2** (10 mM in water; always prepared fresh to minimize compound decomposition) in equal proportion, followed by incubation at room temp for 1-2 hr, as these two substrates can non-enzymatically condense to yield **3** (which can spontaneously decarboxylate to produce **4**). A 2 µL aliquot of this mixture was then added as a component of the *Pt*CAL-1a/*Pt*CAL-2a enzyme reaction (20 µL total reaction volume) in addition to the *Pt*CYP782C1 microsomal reaction mixture. After pre-designated incubation times, reactions were quenched by diluting 10-fold into ACN.

For *Pt*CAL-3 activity assays, 5 µg of *Pt*CAL-3 apoplast (final concentration of 0.1 µg/µL apoplast protein), was diluted into in low pH potassium phosphate buffer (50 mM potassium phosphate, 100 mM NaCl, pH 5.75) at a volume of 50 µL just as with *Pt*CAL-1a/*Pt*CAL-2a. To generate **3** and **4** as potential substrates *in vitro*, 1 µg of previously purified *Pt*PIKS-1 enzyme(*21*) was added to this reaction (final concentration 0.02 µg/µL) with 150 µM **1** and 300 µM malonyl-CoA added as substrates. In follow up experiments, the PIKS reaction was omitted, and **1** and **2** were added as direct substrates to a final concentration of 500 µM each. When **4** was tested as a substrate, it was added at a concentration of 150 µM. All reactions were incubated at room temperature for pre-designated amounts of time, after which aliquots were quenched via 5-fold dilution in ice-cold ACN. For all CAL apoplast enzyme reactions, product formation was predominantly assessed via LC-MS using HILIC analysis. To assess the formation of specific enantiomers, or consumption of specific enantiomeric substrates, quenched reactions were *N*-acetylated and analyzed by chiral LC-MS, as described above.

### *Synthesis of* **6** *stereoisomers*

To synthesize **6** stereoisomers, 150 mg of previously synthesized pelletierine (**4**, oil, 1 mmol)(*21*) was added to 1 mL MeOH in a glass vial with a magnetic stir bar. This mixture was stirred on ice, and 0.095 g (2.5 eq) of NaBH4 was then added slowly. This reaction was allowed to incubate on ice for 2 hr. The reaction was quenched via the addition of 2 mL DI water followed by 2 mL of 2 M HCl. The pH of the reaction was increased to pH 10 with 6 M NaOH (∼0.3mL), and this was then extracted with diethyl ether (5 × 5 mL). The organic fractions were pooled, dried with anhydrous sodium sulfate, clarified using filter, and evaporated to dryness using a rotary evaporator system. A portion of this residue, which would be mainly composed of **5** stereoisomers, was then *O*-acetylated following an established protocol.(*49*) To accomplish this, 50 mg (0.35 mmol) of the synthesized **5** stereoisomers was dissolved in 100 µL of 6 N HCl in a glass vial. Next, 100 µL of acetic acid was added, and this mixture was cooled to ∼0 °C in an ice bath. Once this mixture was chilled, 1 mL of acetyl chloride was slowly added dropwise. This reaction was then incubated in the ice bath for 1 hr, with periodic, gentle mixing. After this incubation, a 1 µL aliquot of this reaction was diluted in 1 mL water +0.1% (v/v) formic acid, and this was analyzed via C18 LC-MS to confirm the formation of the same acetylated compounds that were produced by *Pt*ACT-1. The full reaction was diluted in 25 mL of ice-cold DI water, then clarified through filter paper.

The putative **6** stereoisomers were then purified by using a Sep-Pak C18 12 cc, 2g Vac Cartridge (Waters). To do so, this cartridge was pre-equilibrated with 3 CVs of ACN +0.1% (v/v) formic acid, followed by equilibration with 4 CVs of water +0.1% (v/v) formic acid. The reaction mixture was then loaded onto the cartridge and the solvent was allowed to flow through. The loaded cartridge was then washed with 3 CV of water +0.1% (v/v) formic acid, and the products (visibly yellow on the cartridge) were eluted with 30% ACN in water (with 0.1% v/v formic acid). Small (∼0.5 mL) fractions of the eluent were collected, and 1 µL of each were diluted in water +0.1% (v/v) formic acid and analyzed via C18 LC-MS to confirm the presence of putative **6** diastereomers. Relatively pure fractions were combined, diluted into 20 mL water +0.1% (v/v) formic acid and re-purified over the same type of cartridge, much as described above. For this second round of purification, ACN in water (+0.1% v/v formic acid) was added as an eluent at incrementally increasing concentrations (1 CV each of 2, 4, 6, 8, 10, 20, and 40% ACN). Collected fractions were screened via LC-MS, and pure fractions were combined, frozen, and lyophilized to dryness. The resulting purified compound (∼20 mg) consisted of a yellowish powder. For structural confirmation, this was dissolved in CDCl_3_ and ^1^H and ^13^C NMR analysis was performed using a Varian Inova 500 MHz NMR spectrometer.

### Synthesis of enantioenriched (R)- and (S)-pelletierine (**4**)

Enantiomers of **4** were synthesized by following a previously established protocol.(*50*) To a 25 mL round bottom flask with a magnetic stir bar were added 1-piperideine (**1**, 81mg, 0.97 mmol, 1 eq), acetone (3.26 mL, 44.46 mmol, 46 eq), DMSO (3.26 mL), water (0.41 mL), and either D- or L-proline (21.2 mg, 0.19 mmol, 0.2 eq). L-proline was used to achieve enantioenriched (*S*)-**4**, while D-proline was used to produce enantioenriched (*R*)-**4**.(*50*) The reaction mixtures were stirred for 1 hr at room temperature, after which 10 mL of saturated sodium bicarbonate in water was added. This was then extracted twice with 50 mL of dichloromethane. These organic fractions were combined and then extracted with 50 mL of brine. Residual water was removed from the remaining organic extract via the addition of anhydrous magnesium sulfate, after which this extract was clarified through filter paper and dried on a rotary evaporator system. The remaining yellow/brown oil represented the **4** product. Successful reactions were confirmed by *N*-acetylating a fraction of the product and analyzing via chiral LC-MS, as described above. This method resulted in approximately 70% enantiomeric excess for each specified enantiomer.

### Scaled-up production of CAL-1a/CAL-2a enzymatic product

To achieve milligram quantities of the observed product of *Pt*CAL-1a/*Pt*CAL-2a (*m/z* 164), the leaves of 109 *N. benthamiana* plants (410 g fresh weight) were vacuum infiltrated(*51*) with a combination of *Agrobacterium* strains necessary for engineering the production of this compound (*Pt*LDC-2, *Pt*CAO-1, *Pt*PIKS-1, *Pt*SDR-2, *Pt*ACT-1, *Pt*CYP782C1, *Pt*CAL-1a, *Pt*CAL-2a, and *Pt*CAL-3). To prepare sufficient quantities of *Agrobacterium* for this scale, *Agrobacterium* strains harboring the necessary gene constructs were first streaked on selective LB agar plates (50 µg/mL kanamycin, 30 µg/mL gentamycin) and grown for 2 days at 30 °C to achieve colonies. Single colonies were then used to inoculate 1 L liquid LB cultures (50 µg/mL kanamycin, 30 µg/mL gentamycin), which were shaken overnight at 30 °C and 250 RPM. Bacteria were then pelleted via centrifugated at 5000xg for 10 min, after which they were resuspended in a minimal volume of *Agrobacterium* induction buffer. Bacterial densities were measured via OD600, and strains were mixed together into a 3 L volume of induction buffer such that each strain had a final calculated density of OD600 0.2. This solution was transferred into a plastic beaker, and this was placed into a plastic, vacuum desiccator. Each *N. benthamiana* plant was placed upside-down into the *Agrobacterium* mixture, the desiccator chamber was brought down to vacuum for 2 min using Welch Model 2025 vacuum pump, which removed air from the leaves, and pressure was then slowly released, which results in *Agrobacterium* solution infiltrating the previous air space of the leaves. This process was repeated for all 109 *N. benthamiana* plants. Infiltrated plants were then grown as usual for 6 days, after which they were harvested and stored at -80 °C until compound purification.

To extract metabolites, frozen plant samples were homogenized in a blender along with 1.5 L of 100% ethanol (EtOH). This extract was allowed to incubated overnight at room temperature in a 4 L flask, after which plant material was removed via clarification over filter paper (this was repeated twice to remove particulates). This EtOH extract was then dried on a rotary evaporator with gentle heating from a water bath (∼30 °C), after which ∼50 mL of water still remained. This was resuspended in 400 mL of 3% tartaric acid in water (w/v), and then extracted with 3 × 200 mL ethyl acetate (EtOAc) to remove hydrophobic compounds. The pH of the aqueous extract was then increased to pH 8-9 using sodium bicarbonate, and this was then extracted with 3 × 200 mL EtOAc. LC-MS screening of extracts demonstrated that almost none of the Lycopodium alkaloid-related intermediates were extracted from the aqueous phase at pH 8-9; instead, this fraction largely contained nicotine-related alkaloids that are native to *N. benthamiana* metabolism. The aqueous phase was then basified to pH 10-11 using 6 M NaOH, and this was extracted with 3 × 400 mL EtOAc. LC-MS screening confirmed that nearly all of the Lycopodium alkaloid intermediates, including our desired compound (proposed **9**), could be found in this organic extract. These EtOAc fractions were combined, dried with anhydrous magnesium sulfate, clarified via filter paper, and then evaporated to dryness using a rotary evaporator. The remaining residue (∼170 mg) consisted mainly of a yellow/brown oil. This was re-dissolved in 20 mL of EtOAc, and this was filtered to remove any insoluble components and then evaporated to dryness. This residue was then resuspended in a minimal volume of 50:50 hexanes/EtOAc (∼5 mL), and was purified using a Biotage® Selekt Flash Purification System with a Biotage® Sfär KP-Amino D column (50 µm particle size, 5 g volume). Purification conditions consisted of an initial isocratic elution of 100% hexanes/0% EtOAc for 3 column volumes (CVs), followed by a gradient from 100% hexanes/0% EtOAc to 0% hexanes/100% ethyl acetate over 10 CVs, with a final 5 CVs at 0% hexanes/100% EtOAc. All fractions were collected in 10 mL increments. Each fraction was then screened for **9** via LC-MS with HILIC conditions. This purification strategy allowed for partial purification of our compound. Fractions containing **9** were combined, evaporated to dryness, and subjected to the same purification workflow several times (with smaller fraction sizes) in order to achieve pure **9**. All other fractions, which contained other Lycopodium alkaloid-related compounds, were dried and saved at 4 °C for future use.

Unfortunately, we found that our isolated compound (predicted **9**) was relatively unstable; resuspension of this compound in either deuterated chloroform (CDCl_3_) or deuterated methanol (CD_3_OD) and analysis via ^1^H NMR demonstrated loss of indicative chemical shifts over time, although this did allow us to obtain a crude ^1^H NMR (CDCl_3_, 500 MHz) for this compound (**Figs S18 & S19**). Additionally, upon drying of our sample from CDCl_3_, we noted a color change from yellow/brown to red. Loss of our compound was confirmed via LC-MS analysis. However, we observed that during the course of purification, a compound pertaining to an oxidation of **9** (*m/z* 263.2118; equal to **9** + oxygen) accumulated to high levels. This compound (**9’**) exhibited a similar LC-MS retention time to **9**, and had an MS/MS fragmentation pattern that appeared to indicate a phlegmarine-like scaffold structure, which suggested that it may be an oxidized by-product of **9** (**Fig S20**). As such, we purified this compound using the same strategy outlined above (yield of ∼3 mg), and determined a putative structure (proposed **9’**) via NMR analysis. For **9’**, deuterated acetonitrile (CD_3_CN) was used as a solvent, and spectra were collected on a Varian Inova 600 MHz spectrometer at room temperature (see **Figs S21-S27**).

### Sequence analysis of CAL genes proteins

All analyses of CAL genes and proteins were performed in Geneious (version 2019.2). To generate protein alignments of CAH family proteins, an assortment of protein sequences containing the α-carbonic anhydrase (CAH) domain were downloaded from UniProt (https://www.uniprot.org/). The majority of these downloaded proteins were selected from plant species (these were selected pseudo-randomly in order to capture a breadth of phylogenetic diversity), and we included all CAHs from that plants that, to our knowledge, have been biochemically verified to have canonical CAH activity. We also included sequences from animals, fungi, algae, and bacteria, including multiple proteins that have been biochemically verified to have canonical activity. The human CAII protein (a.k.a. “CA2”, “hCA II”; UniProt ID: P00918) was used as a reference for amino acid numbering in alignments, as this is likely the most rigorously-studied CAH protein.(*36*) The downloaded CAH proteins and the CAH family proteins identified in our transcriptomic data set (for a set of 80 proteins total) were aligned using the MUSCLE algorithm, and phylogenetic trees were constructed using the Neighbor-joining method (100 bootstraps) with the Jukes-Cantor genetic distance model. The trees shown in **Fig 3** and **Fig S8** have been transformed to align all sequences. Shown adjacent to each tree in these figures are the amino acid sequences that align to well-defined active site residues in human CAII. Any changes to these residues are indicated within the figure, and are color-coded by amino acid.

### IC_50_ values for AChE inhibition by lycodane-type Lycopodium alkaloids

Previous work has determined the ability of various Lycopodium alkaloids to inhibit AChE. A selection of these results are compiled and listed in Fig 5. References for each of the IC_50_ values for each of the compounds are cited as follows: lycophlegmarinine D (**14**) (*41*), huperzine B (**15**) (*52*), huperzine C (**16**)(*52*), huperzine A (**17**) (*53*), des-*N*-methyl-a-obscurine (**18**) (*54*), des-N-methyl-b-obscurine (**19**) (*55*), casuarinine H (**20**) (*56*), 8,15-dihydrohuperzine A (**21**) (*57*), lycoplatyrine B (**24**) (*40*). Compounds annotated as “low/not detected” were not found to have AChE inhibition within the detectable range of each experiment in question (typically, IC_50_ values in these experiments were not measurable, or were greater than 30 µM).

## ACKNOWLEDGEMENTS

We thank Prof. Frank Schroeder (Cornell University) and Jack Liu (Stanford University) for assistance and useful discussion related to NMR analysis. We also thank Prof. David Nelson (University of Tennessee) for providing cytochrome P450 nomenclature. Thank you to Prof. Ke Pan (China Pharmaceutical University), Prof. Richmond Sarpong, and Prof. Paul Evans (University College Dublin) for providing us with authentic standards. We acknowledge Prof. George Lomonossoff for providing us with the pEAQ-HT plasmid. The research in this manuscript was supported by NIH R01 GM121527. R.S.N. was supported as a Howard Hughes Medical Institute Fellow of the Life Sciences Research Foundation.

## Notes

### Competing Interest Statement

The authors have declared no competing interest.

