## Supplementary Information for "Neofunctionalized carbonic anhydrases in the biosynthesis of neuroactive plant alkaloids"

[1]Department of Chemical Engineering, Stanford University, Stanford, CA 94305
[2]HHMI, Stanford University, Stanford, CA 94305
[3]Department of Chemistry, Stanford University, Stanford, CA 94305
[4]Biophysics Program, Stanford University, Stanford, CA 94305
[5]Department of Chemistry, Faculty of Science, Universiti Malaya, 50603 Kuala Lumpur, Malaysia
[6]current affiliation: Department of Molecular and Cellular Biology, Harvard University, Cambridge, MA 02138
*co-corresponding authors

| | page(s) |
|---|---|
| **Supplementary Methods** | 2-3 |
| **Supplementary Results** | 4 |
| **Supplementary Figures** | 5-44 |
| Figure S1. Background on the chemistry and biogenesis of Lycopodium alkaloids | 5-6 |
| Figure S2. Functional characterization of *Pt*SDR-1 and *Pt*SDR-2 | 7-8 |
| Figure S3. Functional characterization of *Pt*ACT-1. | 9-10 |
| Figure S4. Functional characterization of *Pt*CYP782C1. | 10-11 |
| Figure S5. Functional characterization of *Pt*CAL-1 and *Pt*CAL-2. | 12-13 |
| Figure S6. *In vitro* characterization of *Pt*CAL-1 and *Pt*CAL-2. | 14-15 |
| Figure S7. Functional characterization of *Pt*CAL-3. | 16-17 |
| Figure S8. Phylogenetic analysis of CAH family proteins across multiple kingdoms of life. | 18 |
| Figure S9. Functional characterization of *Pt*2OGD-4. | 19-20 |
| Figure S10. Functional characterization of *Pt*2OGD-5. | 21-22 |
| Figure S11. Functional characterization of *Pt*ABH-1. | 23-24 |
| Figure S12. Verification and additional characterization of *Pt*2OGD-3 function. | 25-26 |
| Figure S13. Verification and additional characterization of function for *Pt*2OGD-1 and *Pt*2OGD-2. | 27-28 |
| Figure S14. Detection of early biosynthetic intermediates in extracts of *P. tetrastichus*. | 28 |
| Figure S15. Detection of downstream biosynthetic intermediates in extracts of *P. tetrastichus.* | 29-30 |
| Figure S16. $^{1}$H NMR spectrum from the synthesis of 6 stereoisomers. | 30 |
| Figure S17. $^{13}$C NMR spectrum from the synthesis of 6 stereoisomers. | 31 |
| Figure S18. $^{1}$H NMR (crude) of the purified product (putative 9, *m/z* 247.2169) of *Pt*CAL-1/*Pt*CAL-2. | 32 |
| Figure S19. $^{1}$H NMR (crude) of putative 9 (zoomed in). | 33 |
| Figure S20. LC-MS identification of a 9-related oxidized by-product (putative 9’). | 34 |
| Figure S21. NMR assignment of the oxidized scaffold by-product 9’ (*m/z* 263.2118). | 35 |
| Figure S22. $^{1}$H spectrum of the oxidized scaffold by-product 9’ (*m/z* 263.2118). | 36 |
| Figure S23. $^{13}$C spectrum of the oxidized scaffold by-product 9’ (*m/z* 263.2118). | 37 |
| Figure S24. COSY spectrum of the oxidized scaffold by-product 9’ (*m/z* 263.2118). | 38 |
| Figure S25. HMBC spectrum of the oxidized scaffold by-product 9’ (*m/z* 263.2118). | 39 |
| Figure S26. HSQC spectrum of the oxidized scaffold by-product 9’ (*m/z* 263.2118). | 40 |
| Figure S27. TOCSY spectrum of the oxidized scaffold by-product 9’ (*m/z* 263.2118). | 41 |
| Figure S28. $^{1}$H NMR spectrum ($CDCl_3$, 400 MHz) of flabellidine (10) isolated from *Lycopodium platyrhizoma*.(*1*) | 42 |
| Figure S29. $^{13}$C NMR spectrum ($CDCl_3$, 100 MHz) of flabellidine (10) isolated from *Lycopodium platyrhizoma*.(*1*) | 42 |
| Figure S30. $^{1}$H NMR spectrum ($CDCl_3$, 400 MHz) of casuarinine H (20) isolated from *Lycopodium platyrhizoma*. | 43 |
| Figure S31. $^{13}$C NMR spectrum ($CDCl_3$, 100 MHz) of casuarinine H (20) isolated from *Lycopodium platyrhizoma*. | 44 |
| **Supplementary References** | 44 |

**SUPPLEMENTARY METHODS**

### HPLC method parameters

| | |
|---|---|
| **Instrument:** 6520 LC-MS | |
| **Method:** HILIC 17-min gradient | |
| **Column:** Poroshell 120 HILIC-Z column (Agilent, 2.7 µm, 2.1 × 100 mm) | |
| **Solvent A:** water with 0.1% (v/v) formic acid, 10 mM ammonium formate | |
| **Solvent B:** 9:1 ACN:water with 0.1% (v/v) formic acid, 10 mM ammonium formate | |
| **Flow rate:** 0.25 mL/min | |
| **Injection volume:** 2 uL | |
| **Notes:** Used for analyzing early biosynthetic intermediates (**1** thru **9**) | |
| **% Solvent B** | **time (min)** |
| 100 | 0 |
| 100 | 3 |
| 60 | 8 |
| 100 | 9 |
| 100 | 17 |

| | |
|---|---|
| **Instrument:** 6546 LC-MS | |
| **Method:** C18 9-min gradient | |
| **Column:** ZORBAX RRHD Eclipse Plus C18 column (Agilent, 1.8 µm, 2.1 x 50 mm) | |
| **Solvent A:** water with 0.1% (v/v) formic acid | |
| **Solvent B:** ACN with 0.1% (v/v) formic acid | |
| **Flow rate:** 0.6 mL/min | |
| **Injection volume:** 1 uL | |
| **Notes:** Used for analyzing downstream biosynthetic intermediates (**10** thru **24**) | |
| **% Solvent B** | **time (min)** |
| 3 | 0 |
| 3 | 0.50 |
| 50 | 6.50 |
| 95 | 6.51 |
| 95 | 8.00 |
| 3 | 8.01 |
| 3 | 9.00 |

| **Instrument:** 6520 LC-MS | |
|---|---|
| **Method:** Chiral 33-min gradient | |
| **Column:** CHIRALPAK® IC-3 column (Daicel, 3 µm, 4.6 x 100 mm) | |
| **Solvent A:** water with 0.1% (v/v) formic acid | |
| **Solvent B:** ACN with 0.1% (v/v) formic acid | |
| **Flow rate:** 0.4 mL/min | |
| **Injection volume:** 2 uL | |
| **Notes:** Used for analyzing *N*-acetylated stereoisomers of **4**, **5**, and **6**. | |
| **% Solvent B** | **time (min)** |
| 3 | 0 |
| 3 | 1 |
| 70 | 21 |
| 97 | 22 |
| 97 | 27 |
| 3 | 28 |
| 3 | 33 |

### Mass spectrometer parameters

| **Instrument:** 6520 LC-MS | |
|---|---|
| **Mode** | ESI positive, MS |
| **Drying gas temp** | 300 ºC |
| **Drying gas flow rate** | 11 L/min |
| **Nebulizer** | 35 psig |
| **Fragmentor** | 150 V |
| **Skimmer** | 65 V |
| **OCT 1 Rf Vpp** | 750 V |
| **Vcap** | 3500 V |

| **Instrument:** 6546 LC-MS | |
|---|---|
| **Mode** | ESI positive, MS |
| **Drying gas temp** | 325 ºC |
| **Drying gas flow rate** | 10 L/min |
| **Nebulizer** | 35 psig |
| **Fragmentor** | 135 V |
| **Skimmer** | 45 V |
| **OCT 1 Rf Vpp** | 750 V |
| **Vcap** | 4000 V |

### SUPPLEMENTARY RESULTS

#### *Identification of downstream oxidases*

The most highly reduced Lycopodium alkaloid with the same core "lycodane" scaffold as **17** (see **Fig 1b**) is flabellidine (**10**),(*2*) which contains an *N*-acetyl group on the A ring nitrogen. This molecule had previously been isolated from plants that also produce more highly oxidized lycodane-type alkaloids,(*1*) suggesting it to be a logical precursor, and we used this for substrate co-infiltration into *N. benthamiana* leaves expressing our oxidase gene candidates. Through this approach, we identified a pair of 2OGD enzymes that act to consume **10** and produce oxidized products. The first of these enzymes (*Pt*2OGD-4) appeared to oxidize **10** to a molecule with an exact mass that is consistent with the installation of a carbonyl (proposed structure **11,** $[M+H]^+$ = *m/z* 303.2067) (**Fig S9**). Additionally, we observed minor products that corresponded to the addition of a hydroxyl group ($[M+H]^+$ = *m/z* 305.2224). We speculate that this may represent a hemiaminal intermediate en route to the carbonyl, that could isomerize to form different scaffold types such as that of the lycopodine-related alkaloids (**Fig 1b, Fig S9**). The second enzyme (*Pt*2OGD-5) consumed **10** and produced a new compound with an exact mass indicative of a desaturation (proposed structure **12**, $[M+H]^+$ = *m/z* 287.2118) (**Fig S10**). In agreement with these observed activities, transient co-expression of *Pt*2OGD-4 and *Pt*2OGD-5 with **10** as substrate led to the formation of a compound with both the carbonyl and the desaturation (proposed structure **13**, $[M+H]^+$ = *m/z* 301.1911) (**Fig S10**). Without authentic standards for these substrates, it was not initially possible to confirm the location of these oxidations, but we suspected that *Pt*2OGD-4 was catalyzing formation of the carbonyl to yield the ring A lactam, while *Pt*2OGD-5 was installing the 8,15-double bond. Eventually, we determined that **13** was converted into lycophlegmarinine D (**14**) via the *N*-deacetylation activity of *Pt*ABH-1 (**Fig S11**). Because we could confirm the structure of **14**, this verified the proposed activities of *Pt*2OGD-4 and *Pt*2OGD-5 for the installation of the A-ring carbonyl and the 8,15-double bond, respectively.

#### *Characterization of a Lycopodium alkaloid metabolic network*

Once we had established a biosynthetic pathway from **10** to **17**, we next wanted to determine whether we could access other previously reported Lycopodium alkaloids that appear to differ from **17** pathway intermediates only in their degree of unsaturation. To assess the potential production of these molecules, we reconstituted biosynthesis from **10** with specific 2OGD desaturases (*Pt*2OGD-3 and/or *Pt*2OGD-5) omitted from the co-expressed set of enzymes. We found that omission of *Pt*2OGD-5 led to the consecutive production of the following 8,15-dihydro compounds: **18**, **19**, casuarinine H (**20**), and 8,15-dihydrohuperzine A (**21**), which were all confirmed via authentic standards (**Fig 4, Fig S13**). In a similar fashion, the omission of *Pt*2OGD-3 led to the production of putative 2,3-dihydro congeners: 2,3-dihydrohuperzine C (proposed structure **22**, $[M+H]^+$ = *m/z* 245.1658, $[M-NH_2]^+$ = *m/z* 228.1383) and 2,3-dihydrohuperzine A (proposed structure **23**, $[M+H]^+$ = *m/z* 245.1658, $[M-NH_2]^+$ = *m/z* 228.1383), which to our knowledge have not previously been described (**Fig 4, Fig S13**). Finally, omission of both *Pt*2OGD-5 and *Pt*2OGD-3 led to the formation of lycoplatyrine B (**24**) and a previously undescribed Lycopodium alkaloid that we predict to be 2,3,8,15-tetrahydrohuperzine A (proposed structure **25**, $[M+H]^+$ = *m/z* 247.1805, $[M-NH_2]^+$ = *m/z* 230.1539) (**Fig 4, Fig S13**). These results confirmed the relative promiscuity of these downstream enzymes, thereby allowing for the production of a network of Lycopodium alkaloids.

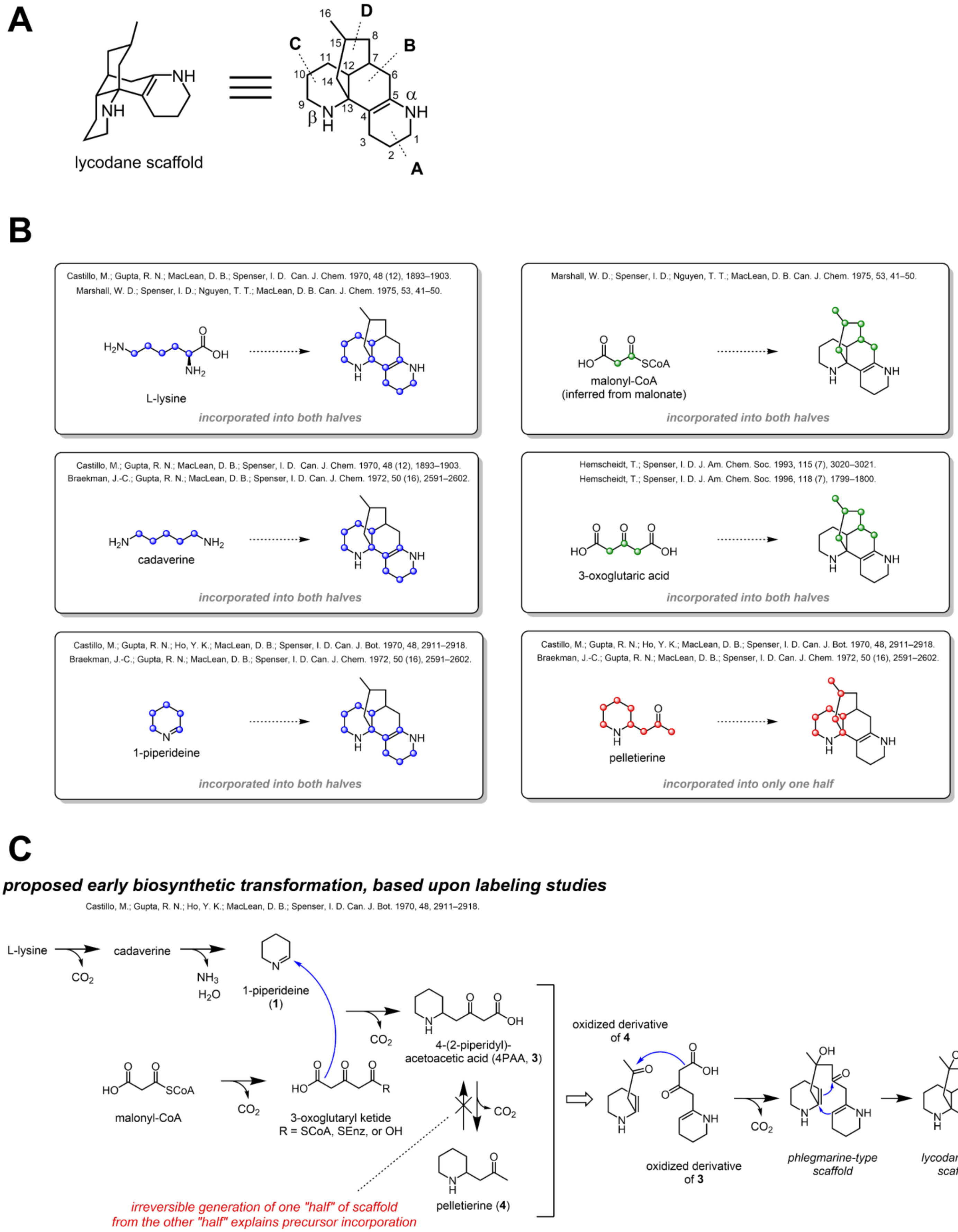

**Figure S1. Background on the chemistry and biogenesis of Lycopodium alkaloids. A)** Chemical schematic of the lycodane scaffold, a major structural class of Lycopodium alkaloid, in two different orientations. Also shown is atom numbering and ring annotation for this scaffold. Note that this is not an observed alkaloid, but a representation of the simplest form of this scaffold. **B)** Summary of previous isotope labeling studies that have defined Lycopodium alkaloid precursors. Colored spheres are used to track location of carbons. Note that these do not represent the specific results of the prior labeling studies, but rather summarize the proposed incorporation of substrate based upon the collective data. Also, isotope labeling studies were performed to evaluate incorporation of substrates into lycopodine, which bears a different scaffold. The incorporation into the lycodane scaffold is inferred, and as previously been postulated.(*3*) Relevant manuscripts for each isotope labeling study are shown within the figure. **C)** Biosynthetic proposal(*3*) accounting for all of the isotope labeling studies. Note that molecules downstream of **3** and **4** have not been evaluated and are hypothetical.

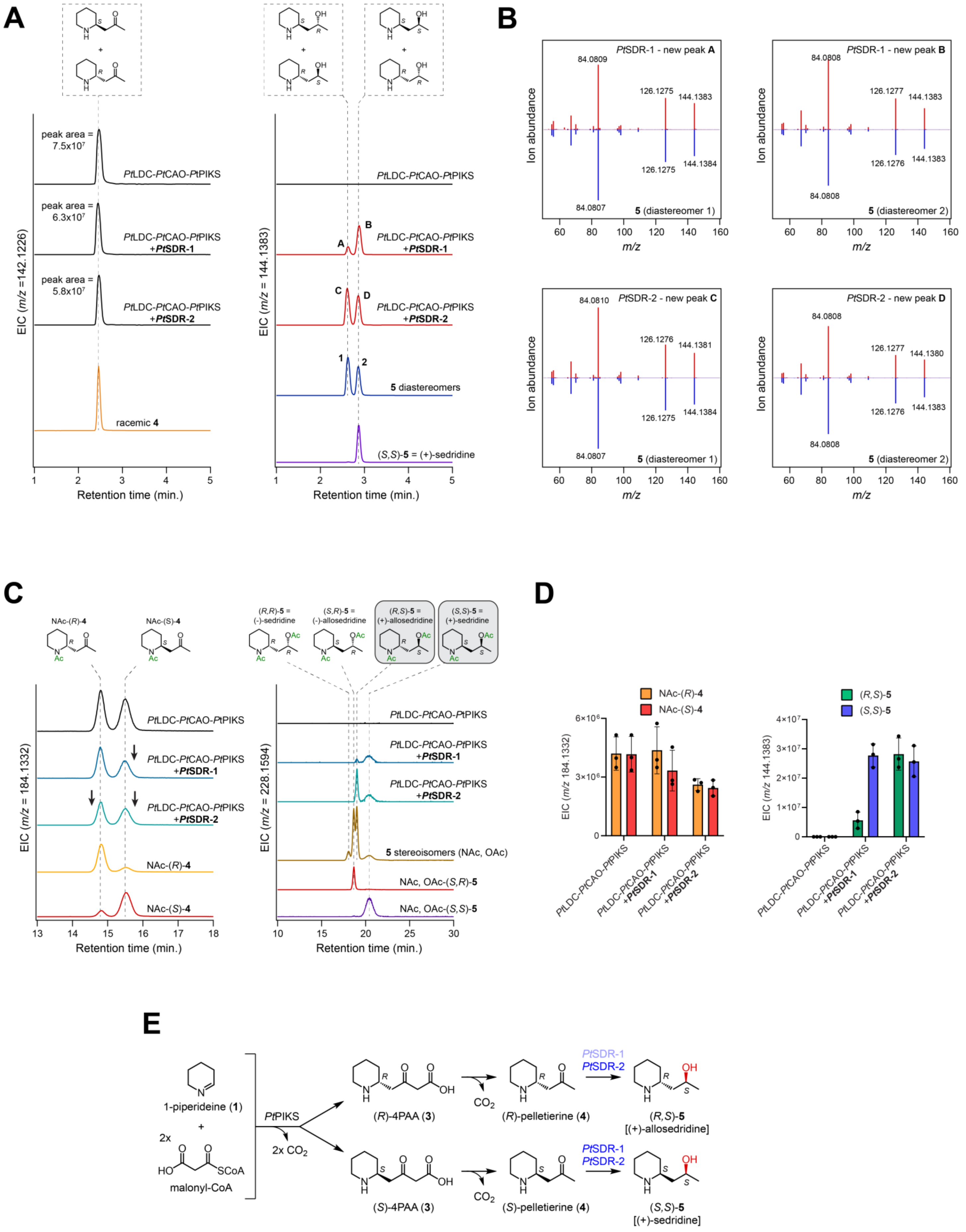
A
peak area = 7.5x10^7
PtLDC-PtCAO-PtPIKS
peak area = 6.3x10^7
PtLDC-PtCAO-PtPIKS +PtSDR-1
peak area = 5.8x10^7
PtLDC-PtCAO-PtPIKS +PtSDR-2
racemic 4
EIC (m/z =142.1226)
Retention time (min.)
EIC (m/z = 144.1383)
A
B
C
D
1
2
5 diastereomers
(S,S)-5 = (+)-sedridine
B
PtSDR-1 - new peak A
PtSDR-1 - new peak B
PtSDR-2 - new peak C
PtSDR-2 - new peak D
5 (diastereomer 1)
5 (diastereomer 2)
Ion abundance
m/z
C
NAc-(R)-4
NAc-(S)-4
(R,R)-5 = (-)-sedridine
(S,R)-5 = (-)-allosedridine
(R,S)-5 = (+)-allosedridine
(S,S)-5 = (+)-sedridine
EIC (m/z = 184.1332)
EIC (m/z = 228.1594)
5 stereoisomers (NAc, OAc)
NAc, OAc-(S,R)-5
NAc, OAc-(S,S)-5
D
NAc-(R)-4
NAc-(S)-4
(R,S)-5
(S,S)-5
EIC (m/z 184.1332)
EIC (m/z 144.1383)
E
1-piperideine (1)
2x malonyl-CoA
PtPIKS
2x CO2
(R)-4PAA (3)
(S)-4PAA (3)
CO2
(R)-pelletierine (4)
(S)-pelletierine (4)
PtSDR-1
PtSDR-2
(R,S)-5 [(+)-allosedridine]
(S,S)-5 [(+)-sedridine]

**Figure S2. Functional characterization of *Pt*SDR-1 and *Pt*SDR-2. A)** Transient expression of *Pt*SDR-1 and *Pt*SDR-2 together with a biosynthetic module for the production of **4** (*Pt*LDC, *Pt*CAO, an *Pt*PIKS) in *N. benthamiana*. Shown are LC-MS extracted ion chromatograms (EICs) for **4** ($[M+H]^+$ = *m/z* 142.1226) and products of *Pt*SDR-1 and *Pt*SDR-2 (A, B, C, and D) that each pertain to a single reduction ($[M+H]^+$ = *m/z* 144.1383), which are shown to represent stereoisomers of **5** via comparison to authentic standards. **B)** MS/MS spectra (*m/z* 144.1383, 20V) for the new compounds produced by *Pt*SDR-1 (A & B) and *Pt*SDR-2 (C & D) in comparison to the co-eluting stereoisomers of **5** standard. **C)** Chiral LC-MS analysis of the biosynthetic products produced in *N. benthamiana*. Samples were *N*-acetylated to allow for retention and separation on a chiral column. Note that hydroxy groups were also acetylated under our derivatization conditions. The left panel shows biosynthetic *N*-acetyl (NAc)-**4** enantiomers ($[M+H]^+$ = *m/z* 184.1332) in comparison to synthesized standards, while the right panel shows biosynthetic NAc, *O*-acetyl (OAc)-**5** stereoisomers ($[M+H]^+$ = *m/z* 228.1594) in comparison to authentic standards. **D)** Quantification of **4** enantiomers (as *N*-acetylated derivatives) that are consumed by *Pt*SDR-1 and *Pt*SDR-2 and **5** diastereomers that are produced within the *N. benthamiana* transient expression system. **E)** Biosynthetic proposal for the production of **5** diastereomers from racemic **4** via the activity of *Pt*SDR-1 or *Pt*SDR-2.

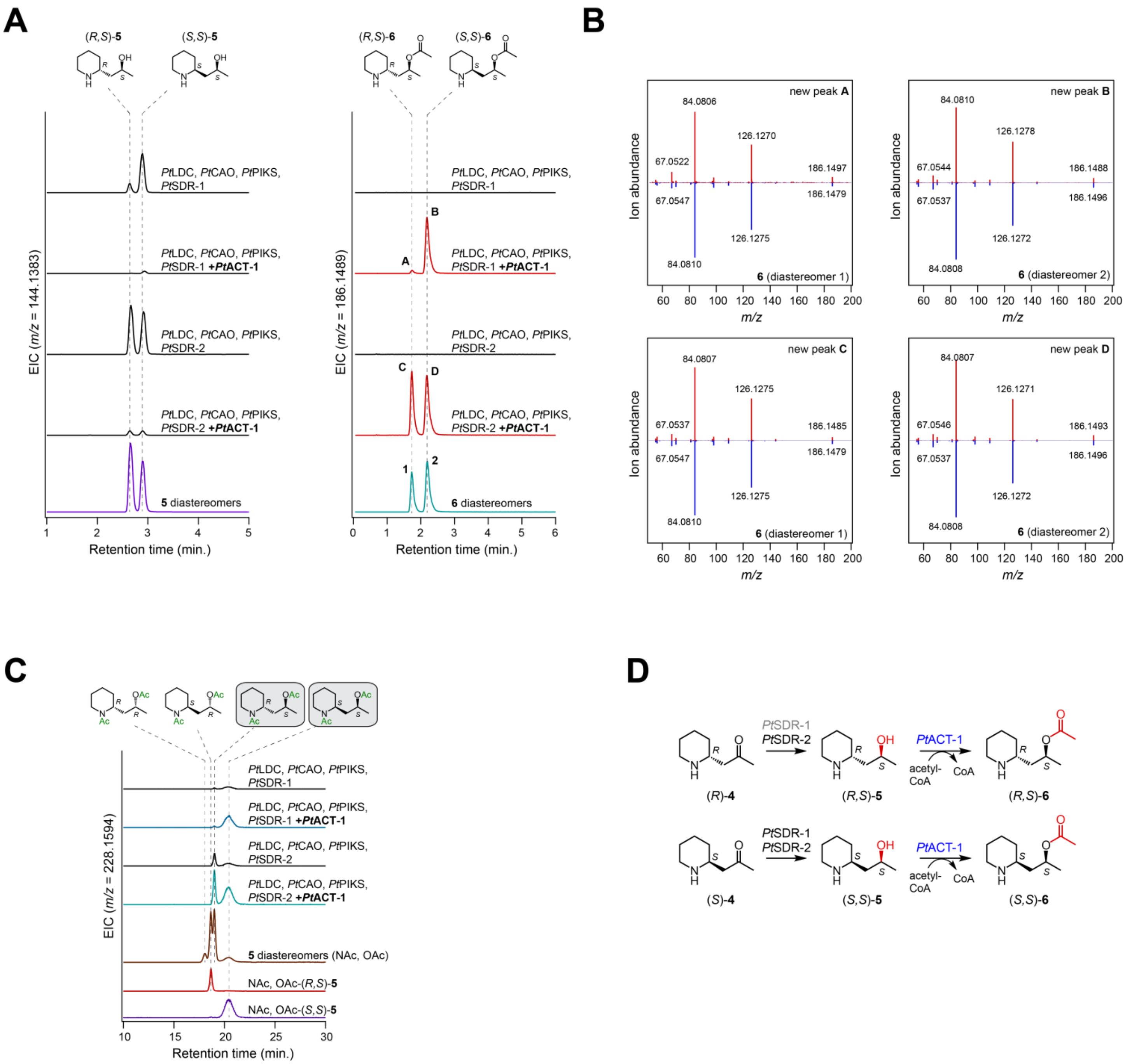

**Figure S3. Functional characterization of *Pt*ACT-1. A)** Transient expression of *Pt*ACT-1 with a biosynthetic module for the production of **5** diastereomers (*Pt*LDC, *Pt*CAO, *Pt*PIKS, and *Pt*SDR-1 or *Pt*SDR-2) in *N. benthamiana*. Shown are LC-MS extracted ion chromatograms (EICs) for **5** diastereomers ($[M+H]^+$ = *m/z* 144.1383) and products of *Pt*ACT-1 (A, B, C, and D) that each pertain to the addition of an acetyl group ($[M+H]^+$ = *m/z* 186.1489), which are shown to represent diastereomers of **6** via comparison to a synthesized standards. **B)** MS/MS spectra (*m/z* 186.1489, 20V) for the new compounds produced by *Pt*ACT-1 (A & B for experiments with *Pt*SDR-1; C & D for experiments with *Pt*SDR-2) in comparison to the co-eluting stereoisomers of **6** standard. **C)** Chiral LC-MS analysis of the biosynthetic products produced by *Pt*ACT-1 in *N. benthamiana*. Samples were *N*-acetylated to allow for retention and separation on a chiral column. Note that hydroxy groups were also acetylated under our derivatization conditions, so products of *Pt*SDR-1/*Pt*SDR-2 would also gain an *O*-acetyl moiety. Shown are biosynthetic *N*-acetyl (NAc)-**6** diastereomers ($[M+H]^+$ = *m/z* 228.1594) in comparison to authentic standards. **D)** Biosynthetic proposal for the activity of *Pt*ACT-1 to convert **5** diastereomers into **6** diastereomers.

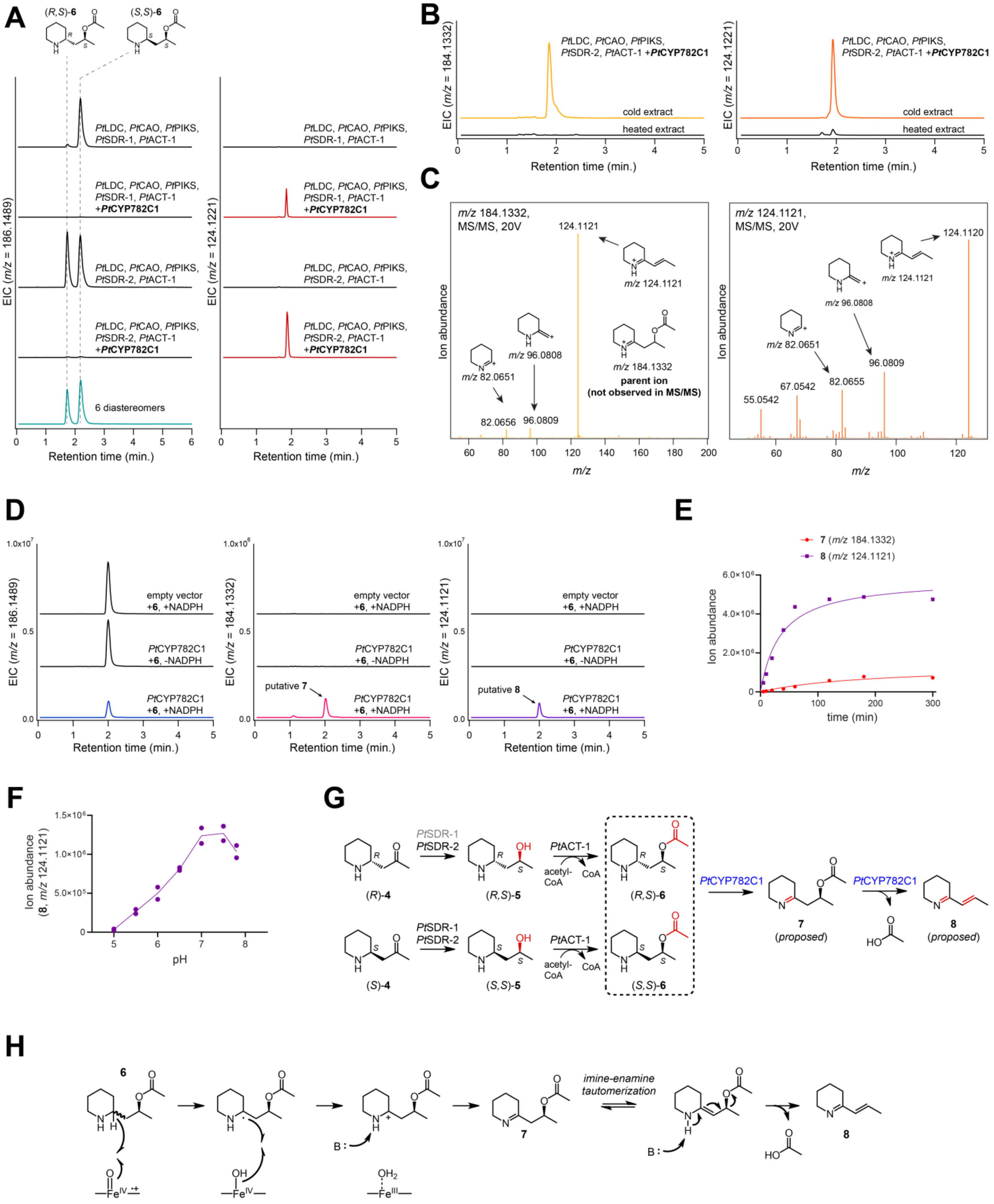

**Figure S4. Functional characterization of *Pt*CYP782C1.** Transient expression of *Pt*CYP782C1 with a biosynthetic module for the production of **6** diastereomers (*Pt*LDC, *Pt*CAO, *Pt*PIKS, *Pt*SDR-1 or *Pt*SDR-2, and *Pt*ACT-1) in *N. benthamiana*. **A)** LC-MS extracted ion chromatograms (EICs) for **6** diastereomers ($[M+H]^+$ = *m/z* 186.1489) and a product of *Pt*CYP782C1 that each represents both an oxidation and elimination of the *O*-acetyl group ($[M+H]^+$ = *m/z* 124.1121). **B)** The two new mass features (left panel, $[M+H]^+$ = *m/z* 184.1332; right panel, $[M+H]^+$ = *m/z* 124.1121) generated by *Pt*CYP782C1 activity were detected in extract that was prepared under cold conditions, but were mostly lost upon incubation at room temperature. **C)** MS/MS spectra of the two new compounds produced by *Pt*CYP782C1 (*m/z* 184.1332, 20V & *m/z* 124.1121, 20V), along with predicted ion fragment structures. **D)** *In vitro* assays with yeast microsomes containing *Pt*CYP782C1 protein. Shown are LC-MS chromatograms representing **6**, as well as the two mass features (**7**, *m/z* 184.1332 & **8**, *m/z* 124.1121) previously identified as putative products of *Pt*CYP782C1. **E)** Formation of **7** and **8** over the course of *in vitro* reaction with *Pt*CYP782C1-enriched yeast microsomes. Product abundance is calculated as the integration of the peak generated in the EIC for each mass ion **F)** Relative activity of *Pt*CYP782C1 microsomes at varying pH **G)** Biosynthetic proposal for the activity of *Pt*CYP782C1 on **6** diastereomers to product **7** and **8**. **H)** Possible catalytic mechanism for the conversion of **6** into **8**.

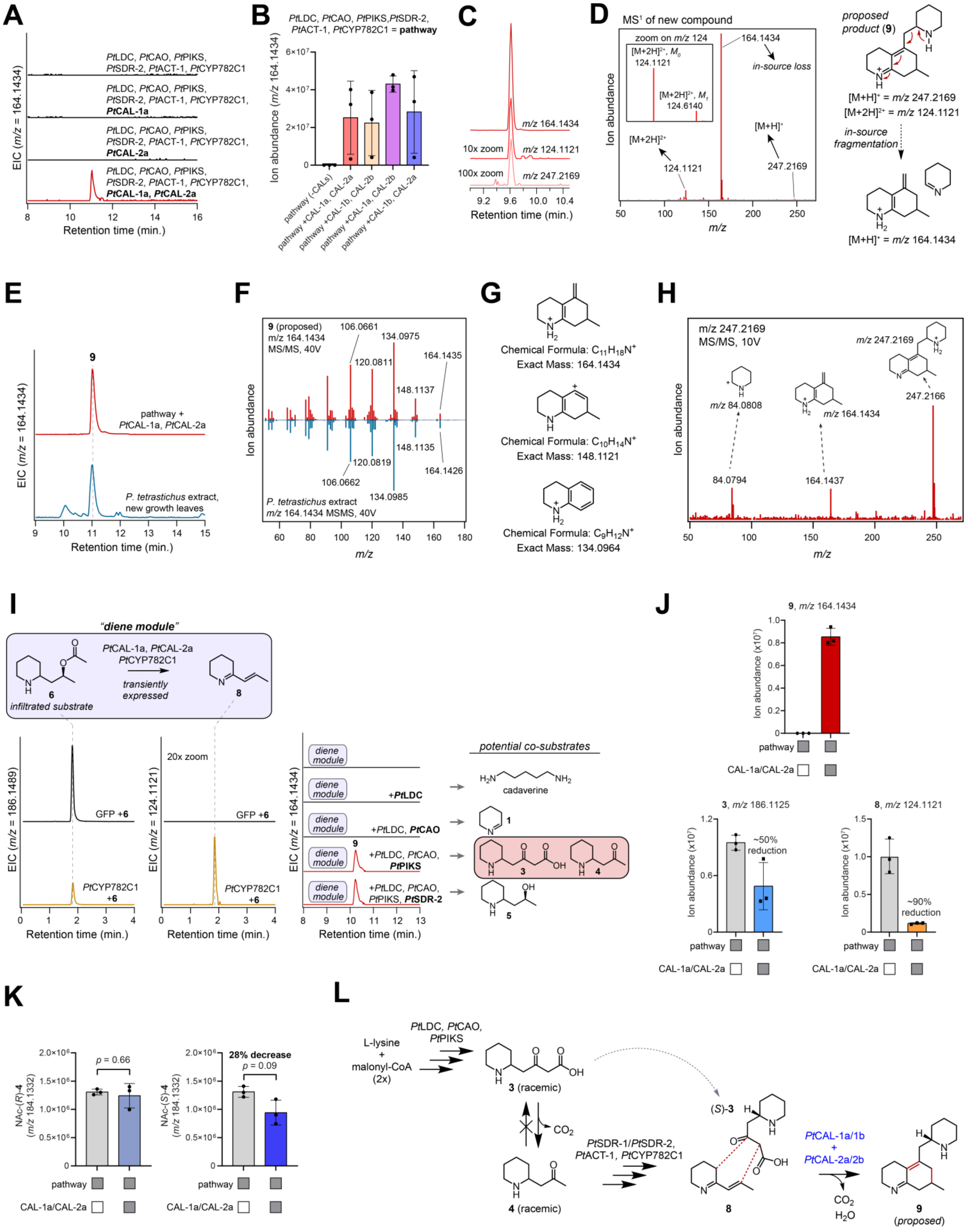
A
B
PtLDC, PtCAO, PtPIKS, PtSDR-2, PtACT-1, PtCYP782C1 = pathway
C
D
MS1 of new compound
proposed product (9)
E
F
G
H
I
"diene module"
PtCAL-1a, PtCAL-2a
PtCYP782C1
transiently expressed
infiltrated substrate
potential co-substrates
cadaverine
J
K
L
L-lysine
malonyl-CoA (2x)
3 (racemic)
4 (racemic)
9 (proposed)

**Figure S5. Functional characterization of *Pt*CAL-1 and *Pt*CAL-2. A)** Transient expression of *Pt*CAL-1a and *Pt*CAL-2a with a biosynthetic module for producing **8** (*Pt*LDC, *Pt*CAO, *Pt*PIKS, *Pt*SDR-2, *Pt*ACT-1, and *Pt*CYP782C1). Shown is an LC-MS extract ion chromatogram (EIC) for the major ion ($[M+H]^+$ = *m/z* 164.1434) associated with the activity of *Pt*CAL-1a/*Pt*CAL-2a when they are both co-expressed in the transient expression system. **B)** Quantification of new product (*m/z* 164.1434) abundance through the activity of *Pt*CAL-1 and *Pt*CAL-2 homologs in different combinations. **C)** Multiple mass ions were found to co-elute with *m/z* 164.1434, suggesting that this ion could be an artefact of in-source fragmentation. **D)** $MS^1$ profile of the new compound generated by *Pt*CAL-1/*Pt*CAL-2. Note the presence of presumed parent mass ions ($[M+H]^+$ = *m/z* 247.2169, $[M+2H]^{2+}$ = *m/z* 124.1121), which suggest that *m/z* 164.1434 results from an in-source loss of 1-piperideine from the proposed product **9** during ionization in the mass spectrometer. **E)** LC-MS EICs (*m/z* 164.1434) comparing the biosynthetic product of *Pt*CAL-1/*Pt*CAL-2 (**9**, proposed) to a co-eluting compound within the new growth leaf tissue of *Phlegmariurus tetrastichus*. **F)** MS/MS spectra (*m/z* 164.1434, 40V) comparing the biosynthetic product to the compound identified within *P. tetrastichus* extract. **G)** Proposed structures for major ion fragments shown in panel F. **H)** MS/MS spectrum (*m/z* 247.2169, 40V) of the parent ion for the new compound with predicted structures of fragments. **I)** Deconvolution of the substrates required for *Pt*CAL-1/*Pt*CAL-2 activity. For this, *Pt*CYP782C1, *Pt*CAL-1, and *Pt*CAL-2 were transiently expressed in *N. benthamiana* and **6** was co-infiltrated as substrate. This demonstrated consumption of **6** (*m/z* 186.1489, left panel) and production of **8** (*m/z* 124.1121, middle panel), but no production of **9**. With this established, different combination of upstream genes were included in the transient co-expression system to provide putative substrates necessary for the formation of **9** (*m/z* 164.1434, right panel). **J)** Production of **9** (*m/z* 164.1434) coincides with the depletion of **3** (*m/z* 186.1125) and **8** (*m/z* 124.1121). Relative product abundance was quantified via integration of peaks generation in EICs. **K)** Chiral chromatography of *N*-acetylated precursors was performed to assess which enantiomer of **3** (measured here via consumption of **4**) serves as the substrate for production of **9**. **L)** Biosynthetic proposal for the condensation of (*S*)-**3** and **8** by *Pt*CAL-1 and *Pt*CAL-2 to product the proposed phlegmarine scaffold of **9**.

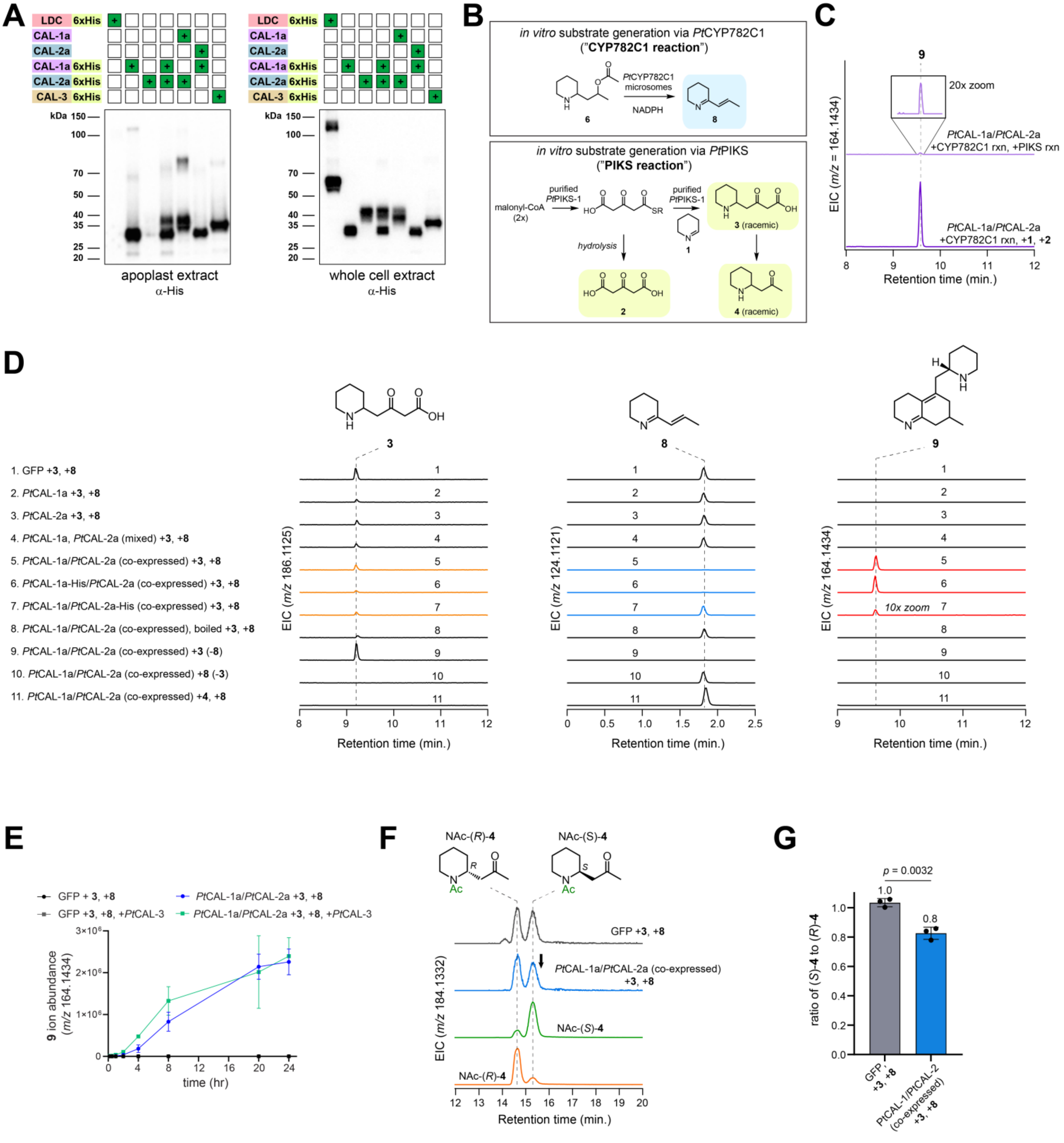

**Figure S6. Characterization of apoplast-enriched *Pt*CAL-1 and *Pt*CAL-2. A)** Western blots of His-tagged CAL protein constructs. The left panel represents protein isolated within the apoplast extract, while the right panel represents protein isolated from the whole cell lysate, which represents "internal" (i.e. non-apoplast) protein (e.g. cytosolic and microsomal protein). The *Lycopodium clavatum* LDC, an ortholog of *Pt*LDC that is proposed to be cytosolic,(*4*) was used as a control for internal protein. **B)** *In vitro* enzyme reactions used to generate putative substrates for *Pt*CAL-1a and *Pt*CAL-2a apoplast extracts. **C)** Confirmation of *Pt*CAL-1a/*Pt*CAL-2a (co-expressed) apoplast activity when the PIKS reaction or **1** & **2** are provided as substrates along with the CYP782C1 reaction. Shown is a LC-MS extracted ion chromatogram (EIC) for **9** (*m/z* 164.1434). Note that production of **9** in this system was dramatically higher when **1** and **2** were used as substrates with the

CYP782C1 reaction. For all other panels in this figure, indication of +**3** indicates that **1** & **2** were used to produce this substrate spontaneously *in vitro*. **D)** *In vitro* apoplast extract reactions with different combinations of apoplast extracts and substrates. The different conditions are listed and number to the left of this panel. Shown are the EICs for the substrates (**3** and **8**) as well as the product (**9**). Note that in these experiments, **3** is generated by the spontaneous condensation of **1** and **2**. **E)** Time course of **9** production, as measured via ion abundance (*m/z* 164.1434) Shown are GFP apoplast extracts (control) or *Pt*CAL-1a/*Pt*CAL-2a (co-expressed) extracts with **3** and **8** generated as *in vitro* substrates. Additionally, the presence of *Pt*CAL-3 within this reaction was assessed here. **F)** Chiral LC-MS EICs analyzing the abundance of *N*-acetylated **4** enantiomers in the *Pt*CAL-1a/*Pt*CAL-2a +**3**, +**8** reaction. Note the decrease in the abundance of NAc-(*S*)-**4** in the presence of *Pt*CAL-1a/*Pt*CAL-2a (indicated with arrow). **G)** Quantification of the ratio between NAc-(*S*)-**4** and NAc-(*R*)-**4** ion abundances in either GFP control reactions, or *Pt*CAL-1a/*Pt*CAL-2a reactions. Average ratios are listed above each bar. n = 3 for each condition. The statistical comparison was made using Welch's test.

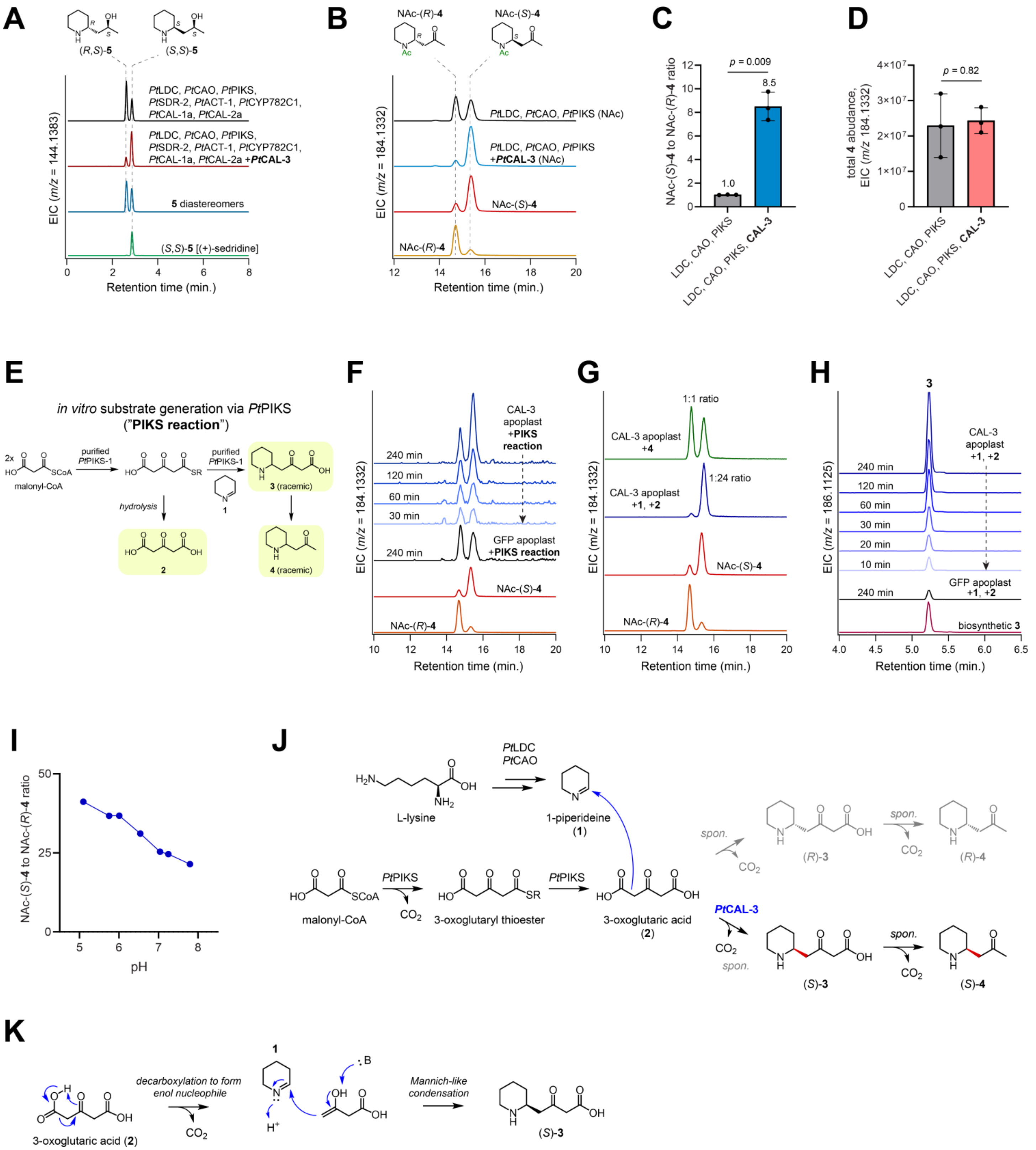

**Figure S7. Functional characterization of *Pt*CAL-3. A)** Transient expression of *Pt*CAL-3 with the pathway to produce **9** (*Pt*LDC, *Pt*CAO, *Pt*PIKS, *Pt*SDR-2, *Pt*ACT-1, *Pt*CYP782C1, *Pt*CAL-1a, and *Pt*CAL-2a. Shown are LC-MS extracted ion chromatograms (EICs) for the **5** diastereomer (*m/z* 144.1383) intermediates that remain within this biosynthetic system. **B)** Chiral LC-MS analysis of *N*-acetylated products from a transient expression system that generates **4** (NAc-**4** = *m/z* 184.1332) with or without co-expression of *Pt*CAL-3. **C)** Effect on the ratio of (*S*)-**4** to (*R*)-**4** when *Pt*CAL-3 is included with *Pt*LDC, *Pt*CAO, and *Pt*PIKS in *N. benthamiana*. **D)** Effect of *Pt*CAL-3 on the total accumulation of **4** in *N. benthamiana*. **E)** To probe the potential substrates of *Pt*CAL-3 *in vitro*, an enzyme reaction with purified *Pt*PIKS-1, malonyl-CoA, and **1** was used to generate **2**, **3**, and **4** within the same reactions as *Pt*CAL-3 apoplast extract. **F)** *In vitro* assay with *Pt*CAL-3-enriched apoplast extract generated from *N. benthamiana* transient expression. Shown here are chiral LC-MS EICs for *N*-acetylated **4** enantiomers (*m/z* 184.1332). Apoplast from plants expressing GFP were used as a negative control. Reactions contained an enzymatic mixture for the production of **2**, **3** and **4** (purified *Pt*PIKS-1 +malonyl CoA, +**1**), as defined in panel E. Note the enrichment of (*S*)-**4** over time in the reactions that contain *Pt*CAL-3. **G)** *In vitro* assay with *Pt*CAL-3 apoplast where either racemic **4** or **1** and **2** (which can spontaneously condense to produce **3**, and subsequently, **4**) are included as substrates. Shown here are chiral LC-MS EICs for *N*-acetylated **4** enantiomers (*m/z* 184.1332). The ratio of enantiomers is listed next to the peaks for each reaction. Note that the ratio is unaffected when **4** is added as the substrate, indicating that *Pt*CAL-3 does not act on this compound. **H)** HILIC LC-MS analysis of *in vitro* time-course *Pt*CAL-3 apoplast reactions where **1** and **2** are used as substrates. Shown are EICs for **3** (*m/z* 186.1125), which accumulates over time in the *Pt*CAL-3 apoplast reaction. Biosynthetic **3** was generated via transient expression of *Pt*LDC, *Pt*CAO, and *Pt*PIKS in *N. benthamiana*, as usual. **I)** Assessment of *Pt*CAL-3 apoplast activity over a range of different pH conditions. This was measured by determining the ratio (*S*)-**4** to (*R*)-**4** (*N*-acetylated derivatives) via chiral LC-MS at the end point of each reaction. **J)** Biosynthetic proposal for the activity of *Pt*CAL-3 in the condensation of **1** and **2** to produce (*S*)-**3**. Additional data to support this is found in Figure 3. **K)** Proposed mechanism for the decarboxylative condensation of **2** with **1**.

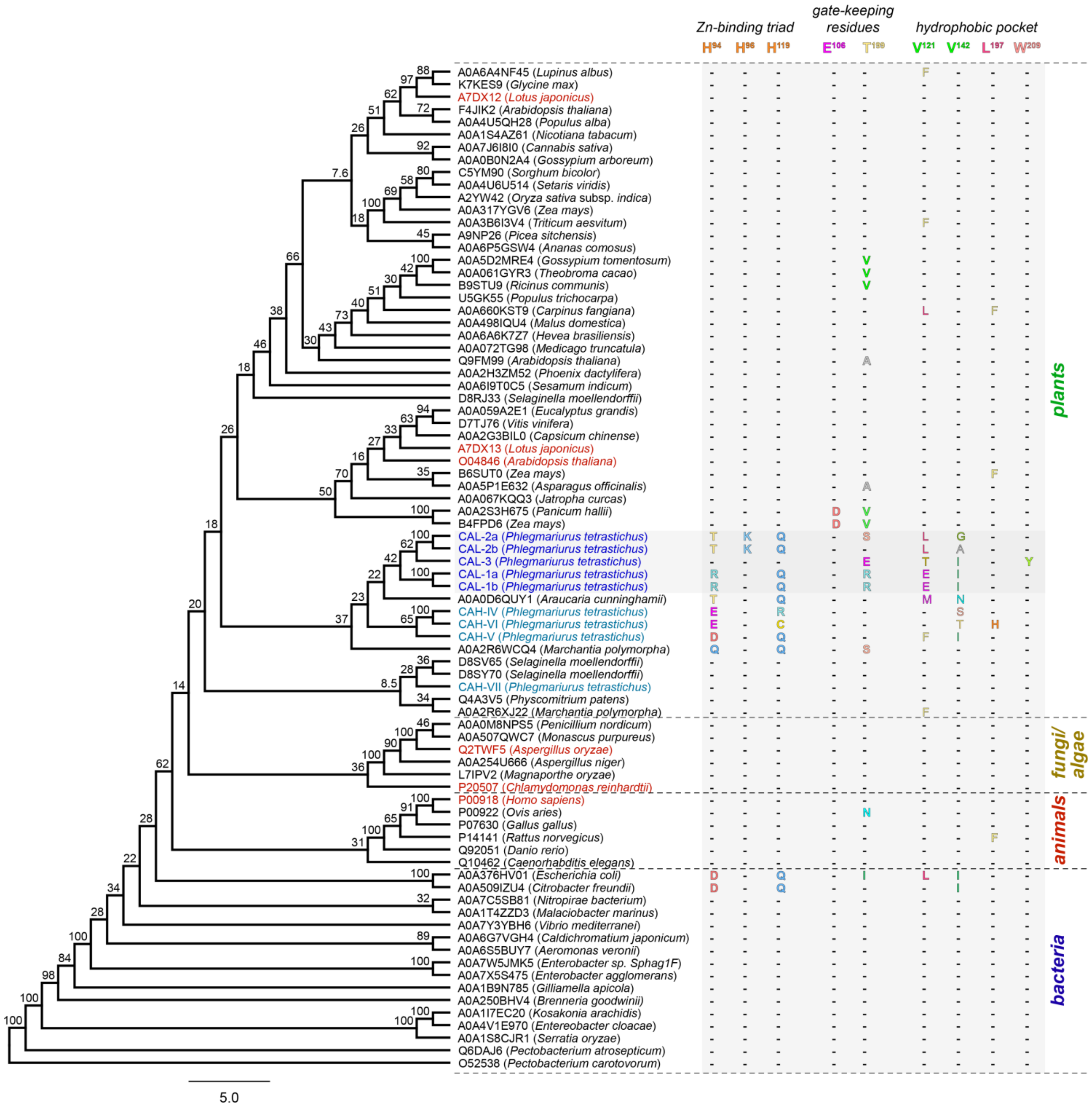

**Figure S8. Phylogenetic analysis of CAH family proteins across multiple kingdoms of life.** Sequences containing an alpha-carbonic anhydrase domain were downloaded from the UniProt database, aligned using MUSCLE, and a phylogenetic tree was generated in Geneious software using a Neighbor-Joining method. Bootstrap values (100 replicates) are shown at nodes. The scale bar indicates substitutions per amino acid Located next to the tree are the amino acid residues that pertain to the histidine-binding triad, gate keeping residues, and substrate binding pocket of canonical carbonic anhydrases. The human CAII protein (UniProt ID: P00918) is used as the reference sequence, and reference amino acid number is based upon this protein. Consensus with this reference is shown as a dash ("-"), while changes are shown by listing the mutated amino acid. Biochemically verified CAH proteins with canonical activity are highlighted in red. The CAL proteins identified and characterized in this study are shown in blue and are highlighted with a gray box. Other CAH-like proteins identified within our *P. tetrastichus* transcriptome, but with unknown function, are shown in light blue. A selected subset of these sequences are shown within the phylogenetic tree in Figure 3.

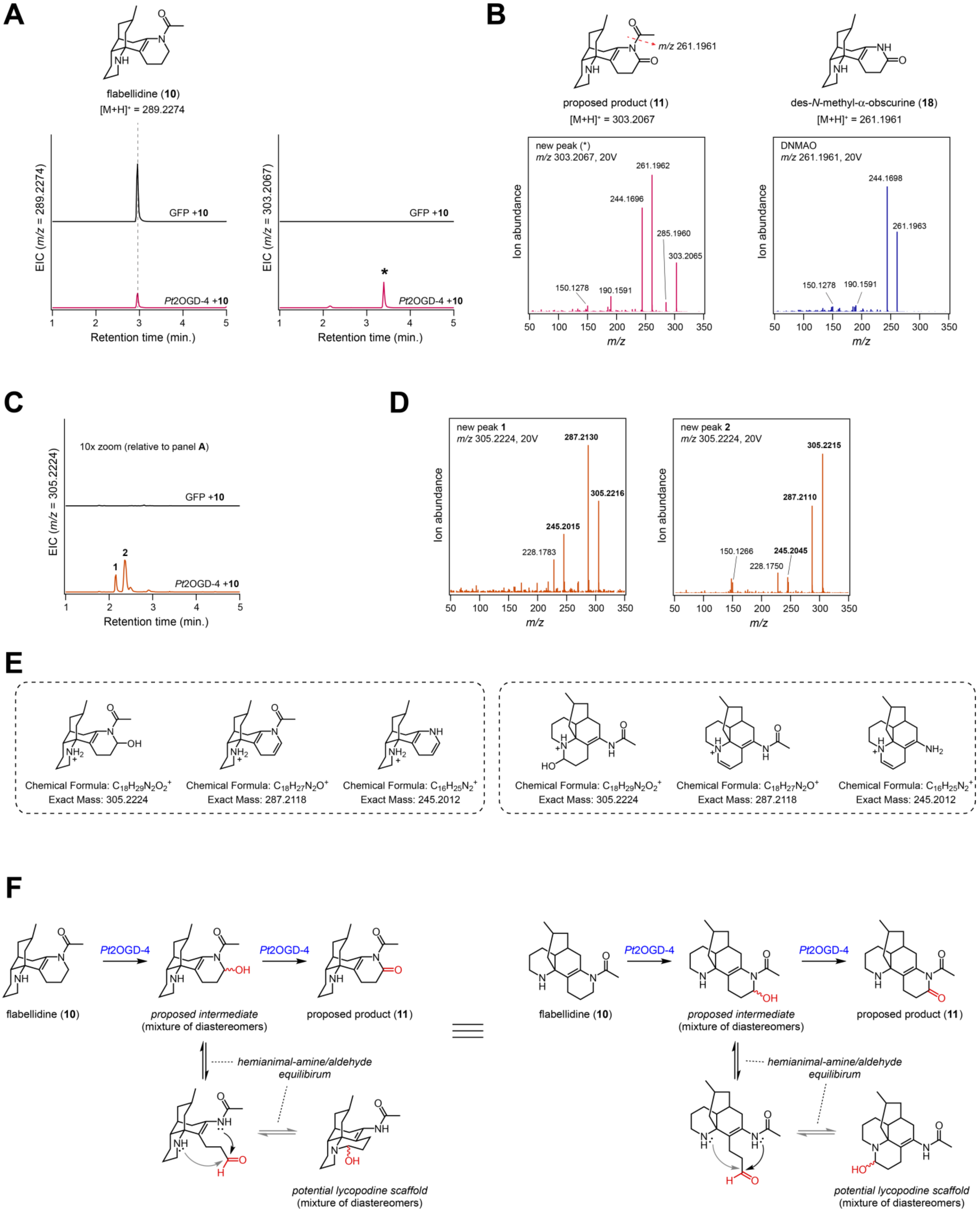
A
flabellidine (10)
[M+H]+ = 289.2274
EIC (m/z = 289.2274)
GFP +10
Pt2OGD-4 +10
Retention time (min.)
EIC (m/z = 303.2067)
*
B
m/z 261.1961
proposed product (11)
[M+H]+ = 303.2067
des-N-methyl-α-obscurine (18)
[M+H]+ = 261.1961
new peak (*)
m/z 303.2067, 20V
261.1962
244.1696
285.1960
303.2065
150.1278
190.1591
Ion abundance
m/z
DNMAO
m/z 261.1961, 20V
244.1698
261.1963
150.1278
190.1591
C
10x zoom (relative to panel A)
EIC (m/z = 305.2224)
GFP +10
Pt2OGD-4 +10
Retention time (min.)
D
new peak 1
m/z 305.2224, 20V
287.2130
305.2216
245.2015
228.1783
new peak 2
m/z 305.2224, 20V
305.2215
287.2110
150.1266
245.2045
228.1750
E
Chemical Formula: C18H29N2O2+
Exact Mass: 305.2224
Chemical Formula: C18H27N2O+
Exact Mass: 287.2118
Chemical Formula: C16H25N2+
Exact Mass: 245.2012
Chemical Formula: C18H29N2O2+
Exact Mass: 305.2224
Chemical Formula: C18H27N2O+
Exact Mass: 287.2118
Chemical Formula: C16H25N2+
Exact Mass: 245.2012
F
Pt2OGD-4
Pt2OGD-4
flabellidine (10)
proposed intermediate
(mixture of diastereomers)
proposed product (11)
hemianimal-amine/aldehyde
equilibirum
potential lycopodine scaffold
(mixture of diastereomers)
Pt2OGD-4
Pt2OGD-4
flabellidine (10)
proposed intermediate
(mixture of diastereomers)
proposed product (11)
hemianimal-amine/aldehyde
equilibirum
potential lycopodine scaffold
(mixture of diastereomers)

**Figure S9. Functional characterization of *Pt*2OGD-4. A)** Transient expression of *Pt*2OGD-4 in *N. benthamiana* with co-infiltration of **10** as substrate. Shown are LC-MS extracted ion chromatograms (EICs) for the **10** substrate ($[M+H]^+$ = *m/z* 289.2274, left panel) and a product (*) of *Pt*2OGD-4 that corresponds to the addition of a carbonyl ($[M+H]^+$ = *m/z* 303.2067, right panel). **B)** MS/MS spectra of the new compound (*m/z* 303.2067, 20V) in comparison to that of **18** (*m/z* 261.1961). Note the similarity in major ion fragments, which suggests that the new compound (proposed as **11**) bears structural similarity to **18**. **C)** Minor products (A & B) pertaining to the addition of a hydroxyl ($[M+H]^+$ = *m/z* 305.2224) are also generated by *Pt*2OGD-4 activity. **D)** MS/MS spectra (*m/z* 305.2224, 20V) for compounds "A" and "B" generated by *Pt*2OGD-4. **E)** Putative structures of the ion fragments shown in bold within panel D. **F)** Biosynthetic proposal for the conversion of **10** into **11** by *Pt*2OGD-4. Note that the right panel shows the same chemistry as the left panel, but in a different 3D orientation.

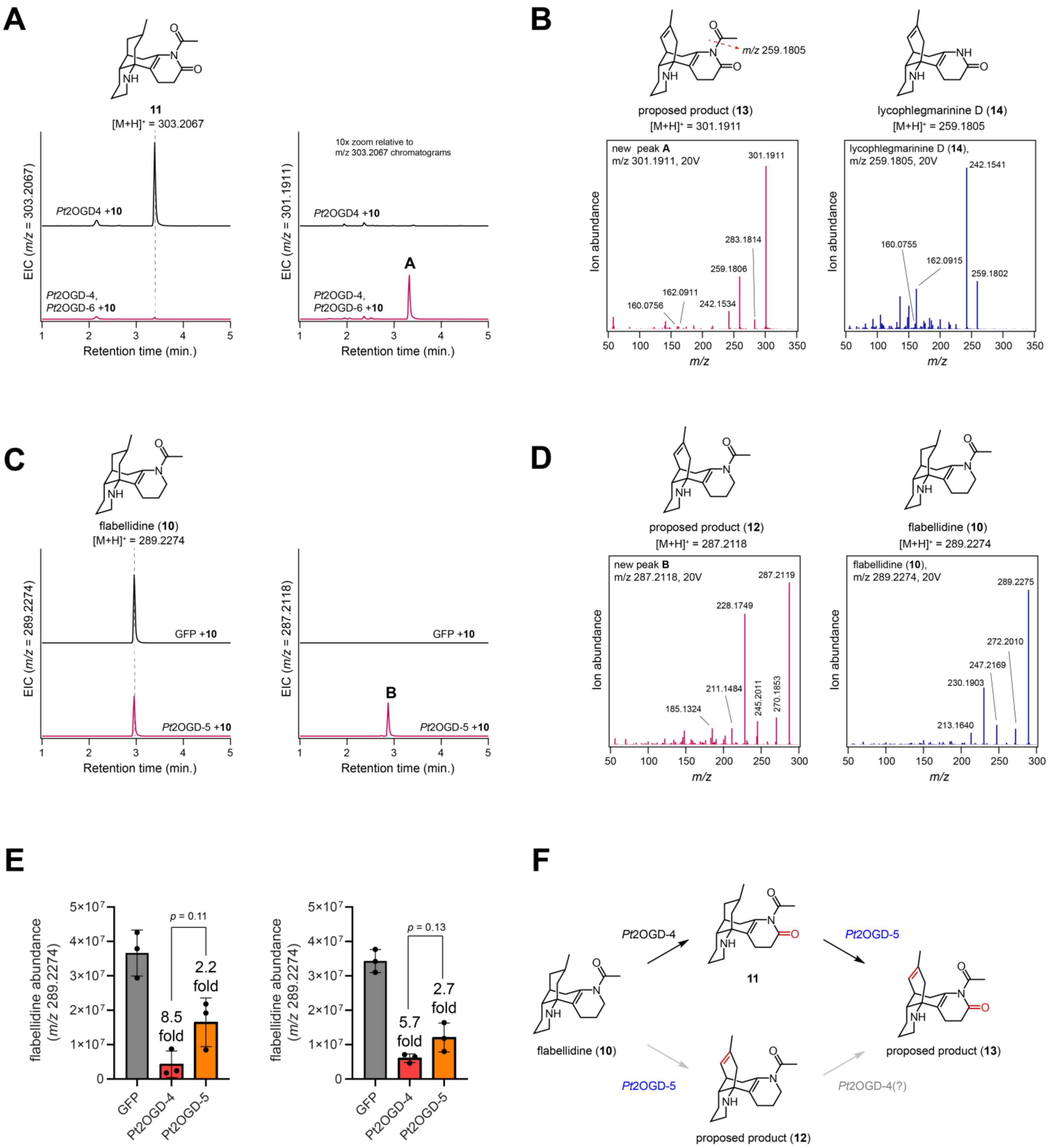

**Figure S10. Functional characterization of *Pt*2OGD-5. A)** Transient expression of *Pt*2OGD-5 with *Pt*2OGD-4 in *N. benthamiana* with co-infiltration of **10** as substrate. Shown are LC-MS extracted ion chromatograms (EICs) for the product of *Pt*2OGD-4 (**11**, *m/z* 303.2067, left panel) and a product (A) of *Pt*2OGD-5 that corresponds to a desaturation ($[M+H]^+$ = *m/z* 301.1911, right panel). **B)** MS/MS spectra of the new compound "A" (*m/z* 301.1911, 20V) in comparison to that of **14** (*m/z* 259.1805). Note the similarity in major ion fragments, which suggests that the new compound (proposed as **12**) bears structural similarity to **14**. **C)** Transient expression of *Pt*2OGD-5 alone in *N. benthamiana* with co-infiltration of **10** as substrate. Shown are LC-MS extracted ion chromatograms (EICs) for **10** as substrate (*m/z* 289.2274, left panel) and a product (B) of *Pt*2OGD-5 that corresponds to a desaturation ($[M+H]^+$ = *m/z* 287.2118, right panel). **D)** MS/MS spectra of the new compound “B” (*m/z* 287.2118, 20V) in comparison to that of **10** (*m/z* 289.2274). Note that the major ion fragments in “B” are typically 2 *m/z* units less than those of **10**, which supports that the new compound (proposed as **13**) bears the same scaffold as **10**, but with a desaturation. **E)** Comparison of **10** consumption by *Pt*2OGD-4 vs. *Pt*2OGD-5. Each of the bar graphs represents an independent experiment. Pairwise comparisons between *Pt*2OGD-4 and *Pt*2OGD-5 reactions were assessed using Welch’s t-test. n = 3 for each experiment. **F)** Biosynthetic proposal for the activity of *Pt*2OGD-5. While *Pt*2OGD-5 can desaturate **10** to produce **13** (putative), *Pt*2OGD-4 appears to have higher activity on **10**, suggesting that *Pt*2OGD-4 activity prior to *Pt*2OGD-5 activity is the major metabolic route for the production of **13** (putative).

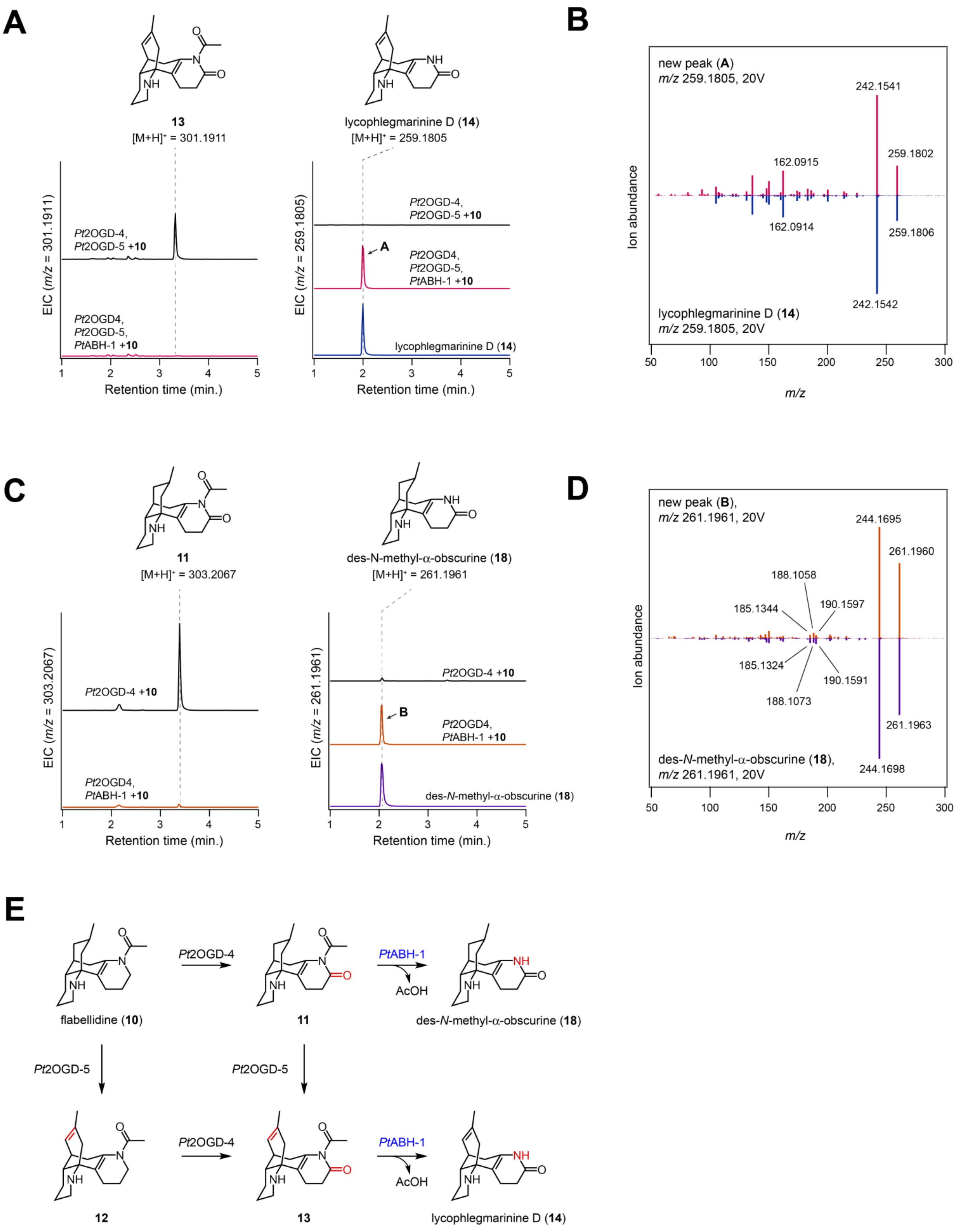
A
13
[M+H]+ = 301.1911
lycophlegmarinine D (14)
[M+H]+ = 259.1805
EIC (m/z = 301.1911)
Pt2OGD-4,
Pt2OGD-5 +10
Pt2OGD4,
Pt2OGD-5,
PtABH-1 +10
Retention time (min.)
EIC (m/z = 259.1805)
Pt2OGD-4,
Pt2OGD-5 +10
A
Pt2OGD4,
Pt2OGD-5,
PtABH-1 +10
lycophlegmarinine D (14)
Retention time (min.)
B
new peak (A)
m/z 259.1805, 20V
242.1541
259.1802
162.0915
162.0914
259.1806
242.1542
lycophlegmarinine D (14)
m/z 259.1805, 20V
Ion abundance
m/z
C
11
[M+H]+ = 303.2067
des-N-methyl-α-obscurine (18)
[M+H]+ = 261.1961
EIC (m/z = 303.2067)
Pt2OGD-4 +10
Pt2OGD4,
PtABH-1 +10
Retention time (min.)
EIC (m/z = 261.1961)
Pt2OGD-4 +10
B
Pt2OGD4,
PtABH-1 +10
des-N-methyl-α-obscurine (18)
Retention time (min.)
D
new peak (B),
m/z 261.1961, 20V
244.1695
261.1960
188.1058
185.1344
190.1597
185.1324
190.1591
188.1073
261.1963
des-N-methyl-α-obscurine (18),
m/z 261.1961, 20V
244.1698
Ion abundance
m/z
E
Pt2OGD-4
PtABH-1
AcOH
flabellidine (10)
11
des-N-methyl-α-obscurine (18)
Pt2OGD-5
Pt2OGD-5
Pt2OGD-4
PtABH-1
AcOH
12
13
lycophlegmarinine D (14)

**Figure S11. Functional characterization of *Pt*ABH-1. A)** Transient expression of *Pt*ABH-1 with *Pt*2OGD-5 and *Pt*2OGD-4 in *N. benthamiana* with co-infiltration of **10** as substrate. Shown are LC-MS extracted ion chromatograms (EICs) for the product of *Pt*2OGD-4 and *Pt*2OGD-5 (**13**, *m/z* 301.1911, left panel) and a product (A) of *Pt*ABH-1 that corresponds to a loss of an acetyl group ($[M+H]^+$ = *m/z* 259.1805, right panel), which is confirmed to be **14** via comparison to an authentic standard. **B)** MS/MS spectra of the new compound "A" (*m/z* 259.1805, 20V) in comparison to that of **14** (*m/z* 259.1805, 20V). **C)** Transient expression of *Pt*ABH-1 with *Pt*2OGD-4 (*Pt*2OGD-5 omitted) in *N. benthamiana* with co-infiltration of **10** as substrate. Shown are LC-MS extracted ion chromatograms (EICs) for the product of *Pt*2OGD-4 (**11**, *m/z* 303.2067, left panel) and a new product (B) of *Pt*ABH-1 that corresponds to the loss of an acetyl-group, ($[M+H]^+$ = *m/z* 261.1961, right panel), which is confirmed to be **18** via comparison to an authentic standard. **D)** MS/MS spectra of the new compound "B" (*m/z* 261.1961, 20V) in comparison to that of **18** (*m/z* 261.1961). **E)** Biosynthetic proposal for the activity of *Pt*ABH-1, which can deacetylate either **11** or **13** to produce **18** or **14**, respectively. Critically, the ability to reach confirmed standards verifies the proposed location of the carbonyl installed by *Pt*2OGD-4 and the double bond installed by *Pt*2OGD-5.

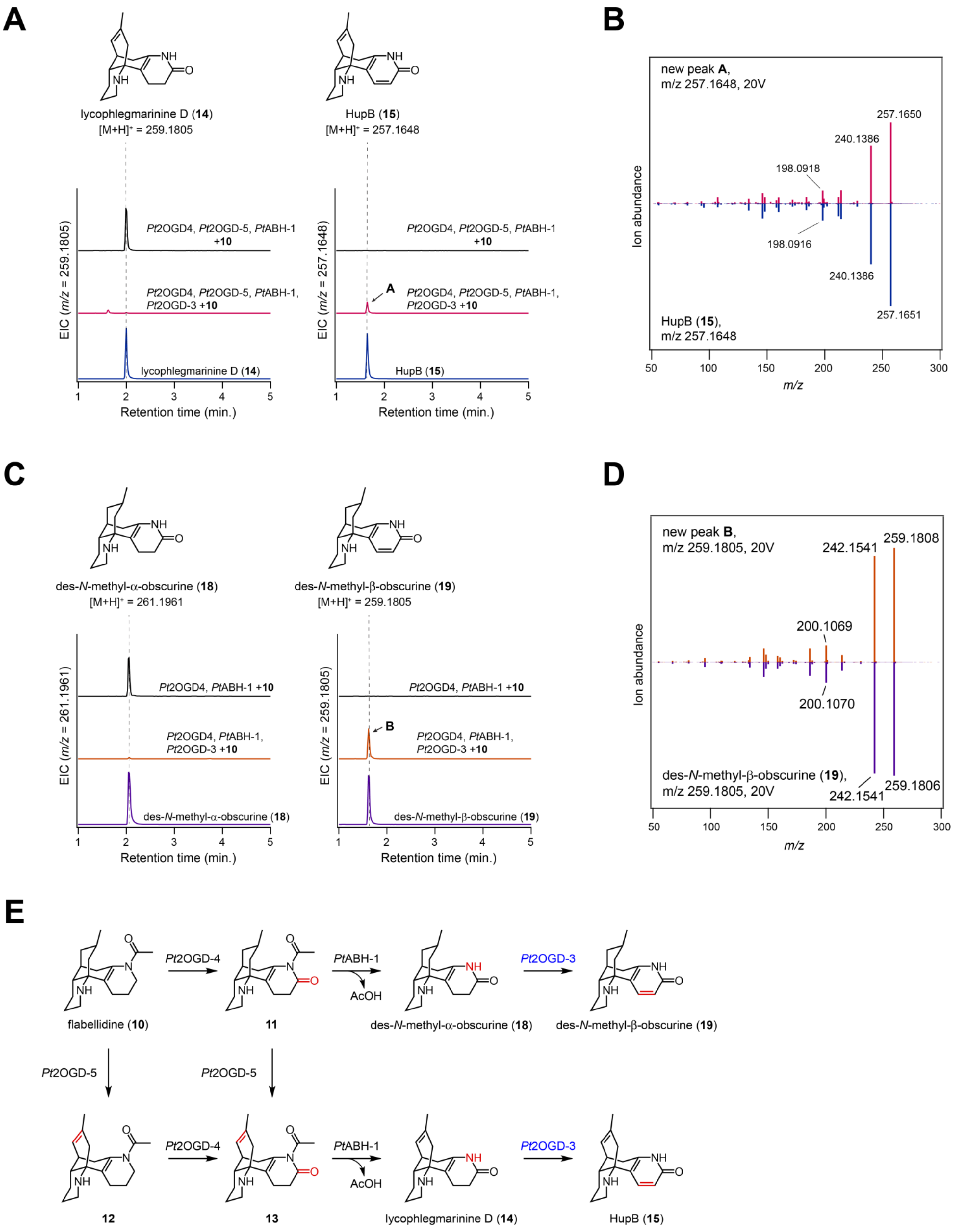

A
lycophlegmarinine D (14)
[M+H]+ = 259.1805
HupB (15)
[M+H]+ = 257.1648
EIC (m/z = 259.1805)
Pt2OGD4, Pt2OGD-5, PtABH-1 +10
Pt2OGD4, Pt2OGD-5, PtABH-1, Pt2OGD-3 +10
lycophlegmarinine D (14)
Retention time (min.)
EIC (m/z = 257.1648)
A
HupB (15)
B
new peak A, m/z 257.1648, 20V
257.1650
240.1386
198.0918
198.0916
240.1386
257.1651
HupB (15), m/z 257.1648
Ion abundance
m/z
C
des-N-methyl-α-obscurine (18)
[M+H]+ = 261.1961
des-N-methyl-β-obscurine (19)
[M+H]+ = 259.1805
EIC (m/z = 261.1961)
Pt2OGD4, PtABH-1 +10
Pt2OGD4, PtABH-1, Pt2OGD-3 +10
des-N-methyl-α-obscurine (18)
EIC (m/z = 259.1805)
B
des-N-methyl-β-obscurine (19)
Retention time (min.)
D
new peak B, m/z 259.1805, 20V
242.1541
259.1808
200.1069
200.1070
des-N-methyl-β-obscurine (19), m/z 259.1805, 20V
242.1541
259.1806
Ion abundance
m/z
E
Pt2OGD-4
PtABH-1
AcOH
Pt2OGD-3
flabellidine (10)
11
des-N-methyl-α-obscurine (18)
des-N-methyl-β-obscurine (19)
Pt2OGD-5
Pt2OGD-5
12
13
lycophlegmarinine D (14)
HupB (15)

**Figure S12. Verification and additional characterization of *Pt*2OGD-3 function. A)** Transient expression of *Pt*2OGD-3 with *Pt*2OGD-4, *Pt*2OGD-5, and *Pt*ABH-1 in *N. benthamiana* with co-infiltration of **10** as substrate. Shown are LC-MS extracted ion chromatograms (EICs) for the product of *Pt*2OGD-4/*Pt*2OGD-5/*Pt*ABH-1 (**14**, *m/z* 259.1805, left panel) and the product (A) of *Pt*2OGD-3 that corresponds to a desaturation ($[M+H]^+$ = *m/z* 257.1648, right panel), which is confirmed to be **15** via comparison to an authentic standard. **B)** MS/MS spectra of the new compound "A" (*m/z* 257.1648, 20V) in comparison to that of **15** (*m/z* 257.1648, 20V). **C)** Transient expression of *Pt*2OGD-3 with *Pt*2OGD-4 and *Pt*ABH-1 (*Pt*2OGD-5 omitted) in *N. benthamiana* with co-infiltration of **10** as substrate. Shown are LC-MS extracted ion chromatograms (EICs) for the product of *Pt*2OGD-4/*Pt*ABH-1 (**18**, *m/z* 261.1961, left panel) and a new product (B) of *Pt*2OGD-3 that corresponds to a desaturation, ($[M+H]^+$ = *m/z* 259.1805, right panel), which is confirmed to be **19** via comparison to an authentic standard. **D)** MS/MS spectra of the new compound "B" (*m/z* 259.1805, 20V) in comparison to that of **19** (*m/z* 259.1805). **E)** Biosynthetic proposal for the activity of *Pt*2OGD-3, which can act on either **18** or **14** to yield the A ring pyridone-containing structures of **19** or **15**, respectively. Note that we had previously demonstrated *Pt*2OGD-3 activity on **18** as a substrate.(*5*)

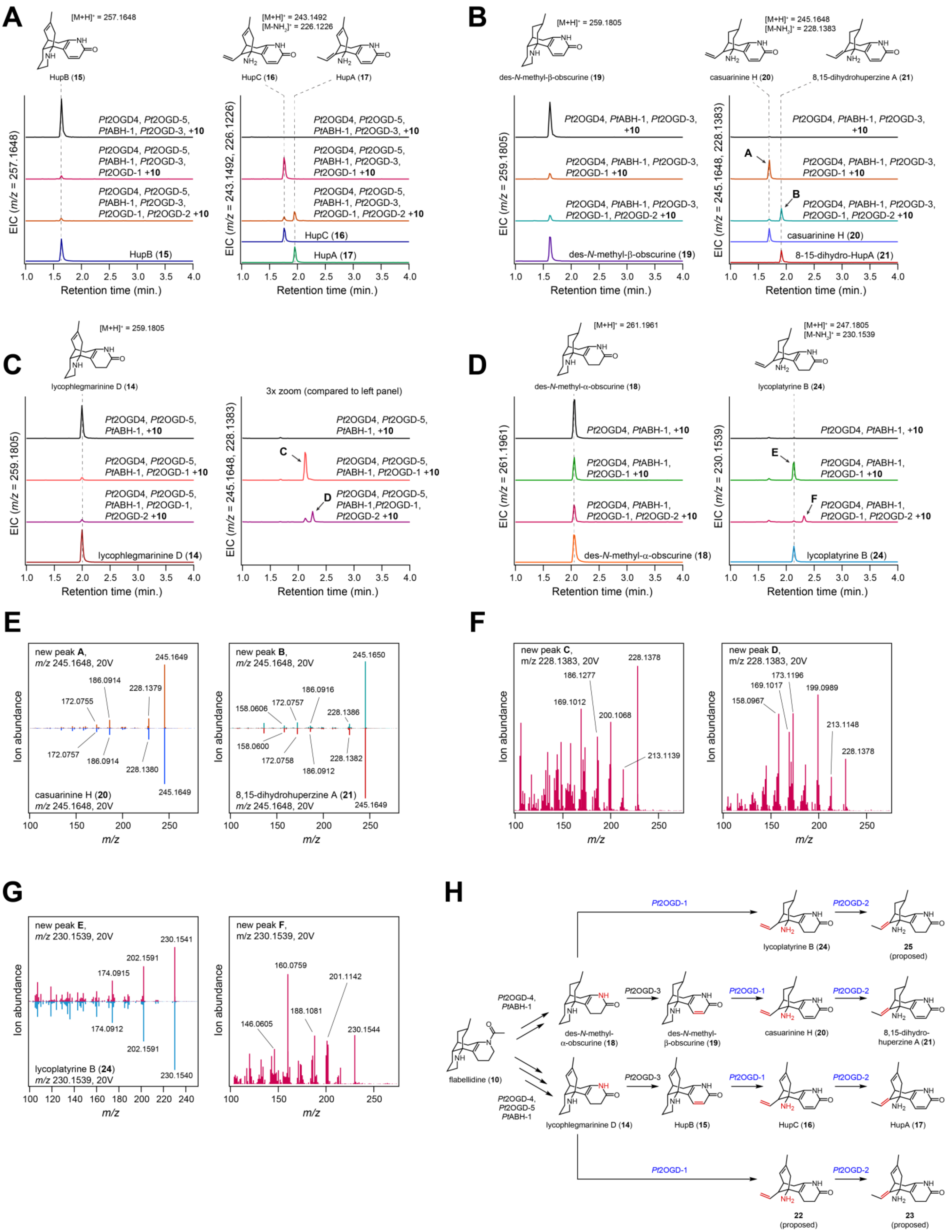

A
HupB (15)
HupC (16)
HupA (17)
B
des-N-methyl-β-obscurine (19)
casuarinine H (20)
8,15-dihydrohuperzine A (21)
C
lycophlegmarinine D (14)
3x zoom (compared to left panel)
D
des-N-methyl-α-obscurine (18)
lycoplatyrine B (24)
E
F
G
H
flabellidine (10)
Retention time (min.)
Ion abundance
m/z

**Figure S13. Verification and additional characterization of function for *Pt*2OGD-1 and *Pt*2OGD-2. A)** Transient expression of *Pt*2OGD-1 and *Pt*2OGD-2 with *Pt*2OGD-4, *Pt*2OGD-5, *Pt*ABH-1, and *Pt*OGD-3 in *N. benthamiana* with co-infiltration of **10** as substrate. Shown are LC-MS extracted ion chromatograms (EICs) for the product of *Pt*2OGD-4/*Pt*2OGD-5/*Pt*ABH-1/*Pt*2OGD-3 (**15**, *m/z* 257.1648, left panel) and the products of *Pt*2OGD-1 and *Pt*2OGD-2 ($[M+H]^+$ = *m/z* 243.1492, $[M-NH_2]^+$ = 226.1226, right panel), which are confirmed to be **16** and **17** via comparisons to authentic standards. **B)** Transient expression of *Pt*2OGD-1 and *Pt*2OGD-2 with *Pt*2OGD-4, *Pt*ABH-1, and *Pt*2OGD-3 (*Pt*2OGD-5 omitted) in *N. benthamiana* with co-infiltration of **10** as substrate. Shown are LC-MS extracted ion chromatograms (EICs) for the product of *Pt*2OGD-4/*Pt*ABH-1/*Pt*2OGD-3 (**19**, *m/z* 259.1805, left panel) and two new products (A & B) from the activities of *Pt*2OGD-1 and *Pt*2OGD-2 ($[M+H]^+$ = *m/z* 245.1648, $[M-NH_2]^+$ = 228.1383 right panel), which are confirmed to be **20** and **21** via comparisons to authentic standards. **C)** Transient expression of *Pt*2OGD-1 and *Pt*2OGD-2 with *Pt*2OGD-4, *Pt*ABH-1, and *Pt*2OGD-5 (*Pt*2OGD-3 omitted) in *N. benthamiana* with co-infiltration of **10** as substrate. Shown are LC-MS extracted ion chromatograms (EICs) for the product of *Pt*2OGD-4/*Pt*ABH-1/*Pt*2OGD-5 (**14**, *m/z* 259.1805, left panel) and two new products (C & D) from the activities of *Pt*2OGD-1 and *Pt*2OGD-2 ($[M+H]^+$ = *m/z* 245.1648, $[M-NH_2]^+$ = 228.1383 right panel), that putative pertain to the 2,3-dihydro congeners of **16** and **17**. **D)** Transient expression of *Pt*2OGD-1 and *Pt*2OGD-2 with *Pt*2OGD-4 and *Pt*ABH-1 (*Pt*2OGD-3 and *Pt*2OGD-5 omitted) in *N. benthamiana* with co-infiltration of **10** as substrate. Shown are LC-MS extracted ion chromatograms (EICs) for the product of *Pt*2OGD-4/*Pt*ABH-1/ (**18**, *m/z* 261.1961, left panel) and two new products (E & F) from the activities of *Pt*2OGD-1 and *Pt*2OGD-2 ($[M+H]^+$ = *m/z* 247.1805, $[M-NH_2]^+$ = 230.1539 right panel). "E" is confirmed to be **24** via comparison to an authentic standard, and "F" is proposed to be the 2,3,8,15-tetrahydro congener of **17**. **E)** MS/MS spectra of the new compound "A" (*m/z* 245.1648, 20V) in comparison to that of **20** (*m/z* 245.1648, 20V) and "B" (*m/z* 245.1648, 20V) in comparison to **21** (*m/z* 245.1648, 20V). **F)** MS/MS spectra of the new compounds "C" and "D" (both *m/z* 228.1383, 20V). **G)** MS/MS spectra of the new compound "E" (*m/z* 230.1539, 20V) in comparison to that of **24** (*m/z* 230.1539, 20V), as well as that of "F" (*m/z* 230.1539. **H)** Biosynthetic proposal for the activities of *Pt*2OGD-1 and *Pt*2OGD-2. These enzymes appear to be able to act independently of the degree of unsaturation of precursors. Note that we had previously demonstrated *Pt*2OGD-1 and *Pt*2OGD-2 activity for conversion of **15** into **16** and **17**, as well as the conversion of **19** into **20** and **21**.(*5*) However, in this prior work, we were not able to confirm the nature of **20** and **21** with authentic standards, as shown here.

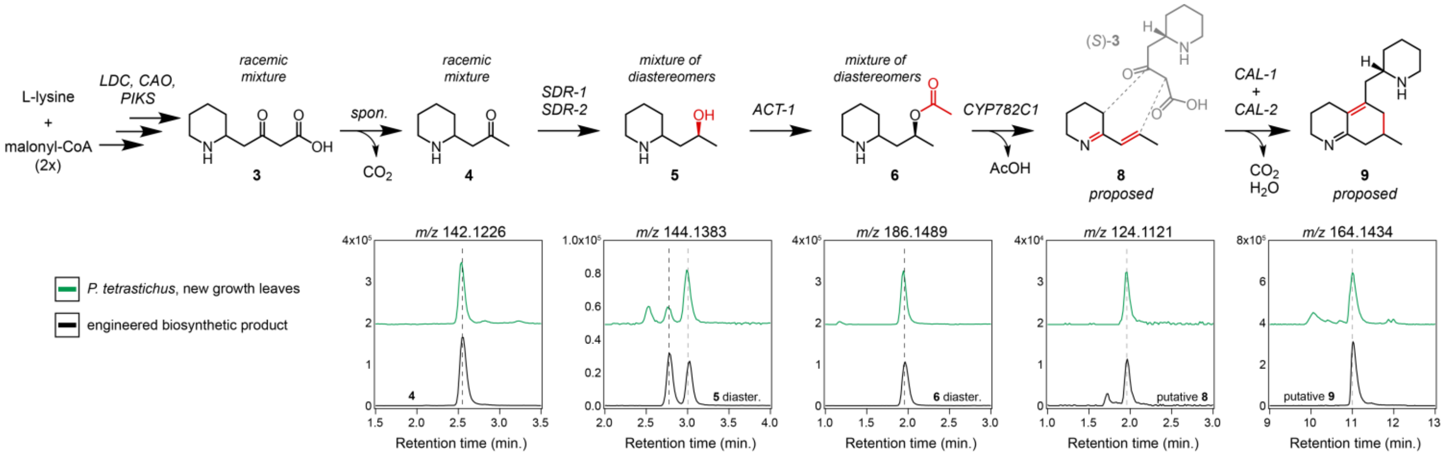

**Figure S14. Detection of early biosynthetic intermediates in extracts of *P. tetrastichus*.** Shown are LC-MS extracted ion chromatograms (EICs) for the various upstream intermediates of Lycopodium alkaloid biosynthesis from the extracts of the native plant (*P. tetrastichus*, green traces) and from the compounds generated through metabolic engineering in *N. benthamiana* (black traces). All samples here were analyzed with HILIC LC-MS. Note that diastereomers of **6** are not separated via HILIC analysis.

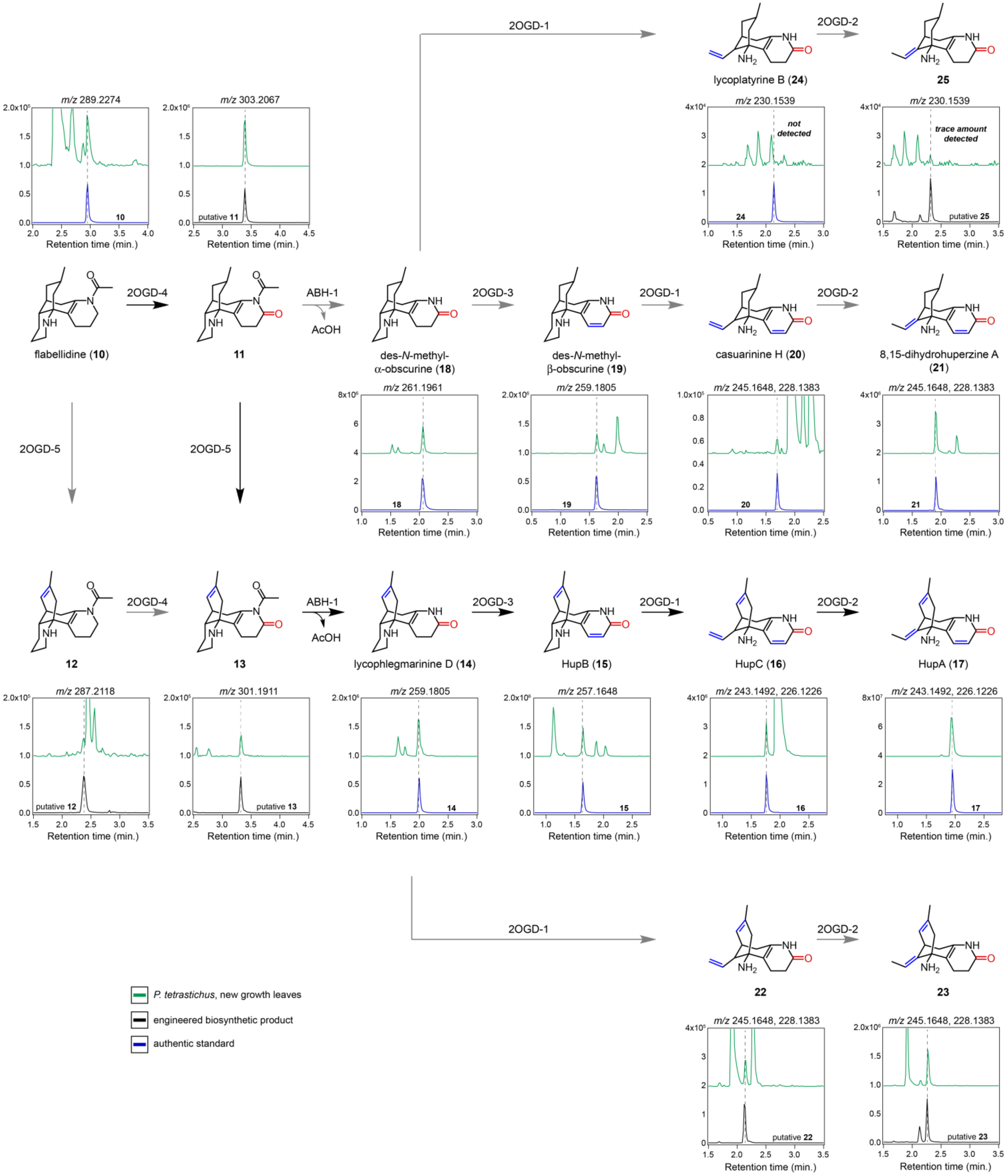

**Figure S15. Detection of downstream biosynthetic intermediates in extracts of *P. tetrastichus*.** Shown are LC-MS extracted ion chromatograms (EICs) for the downstream intermediates of Lycopodium alkaloid biosynthesis from the extracts of the native plant (*P. tetrastichus*, green traces), from the compounds generated through pathway engineering in

*N. benthamiana* (black traces), and for authentic standards (blue), when available. All samples here were analyzed with C18 LC-MS.

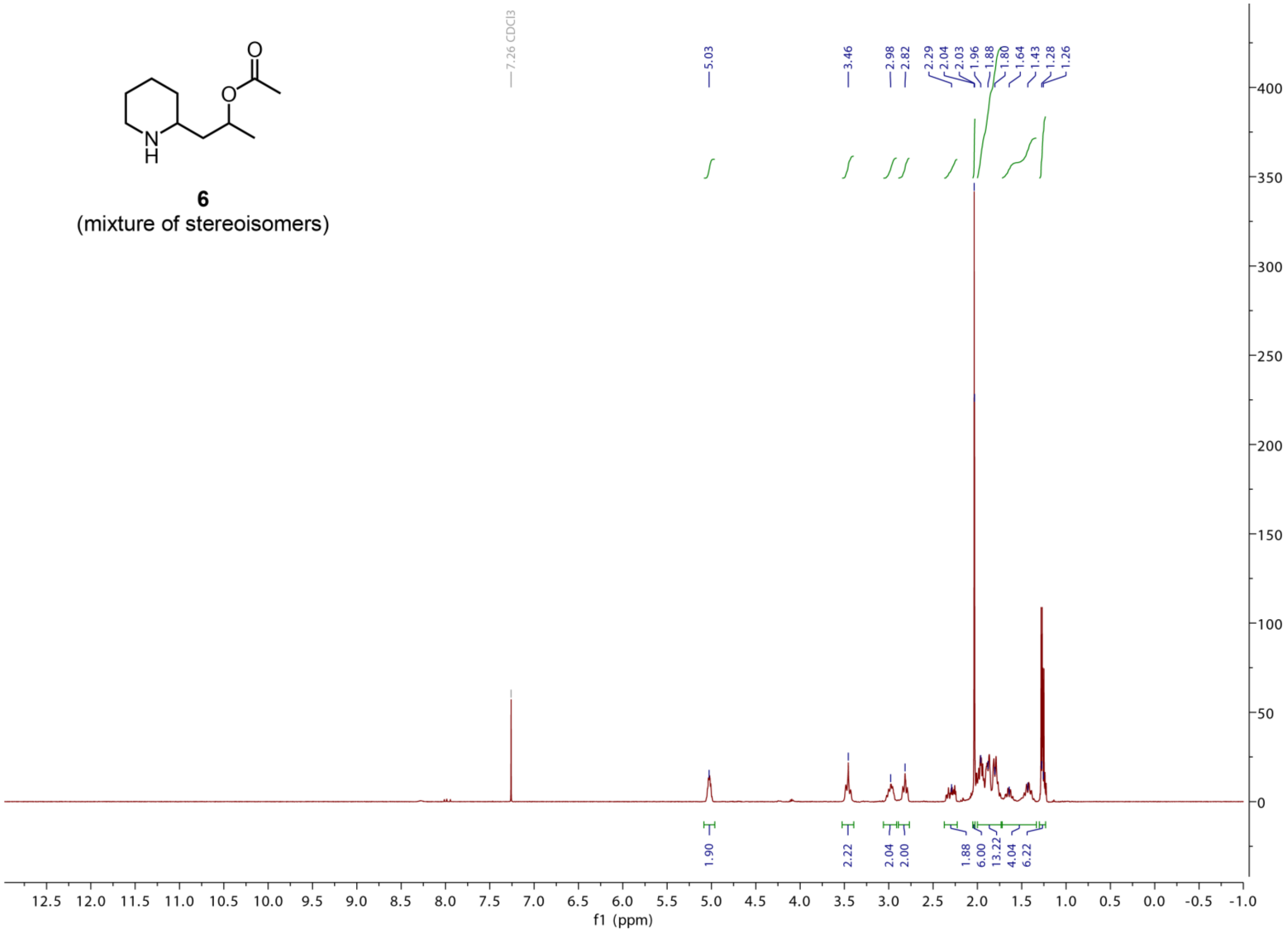

**Figure S16. $^{1}$H NMR spectrum from the synthesis of 6 stereoisomers.** This experiment was recorded at 500 MHz in deuterated chloroform ($CDCl_3$).

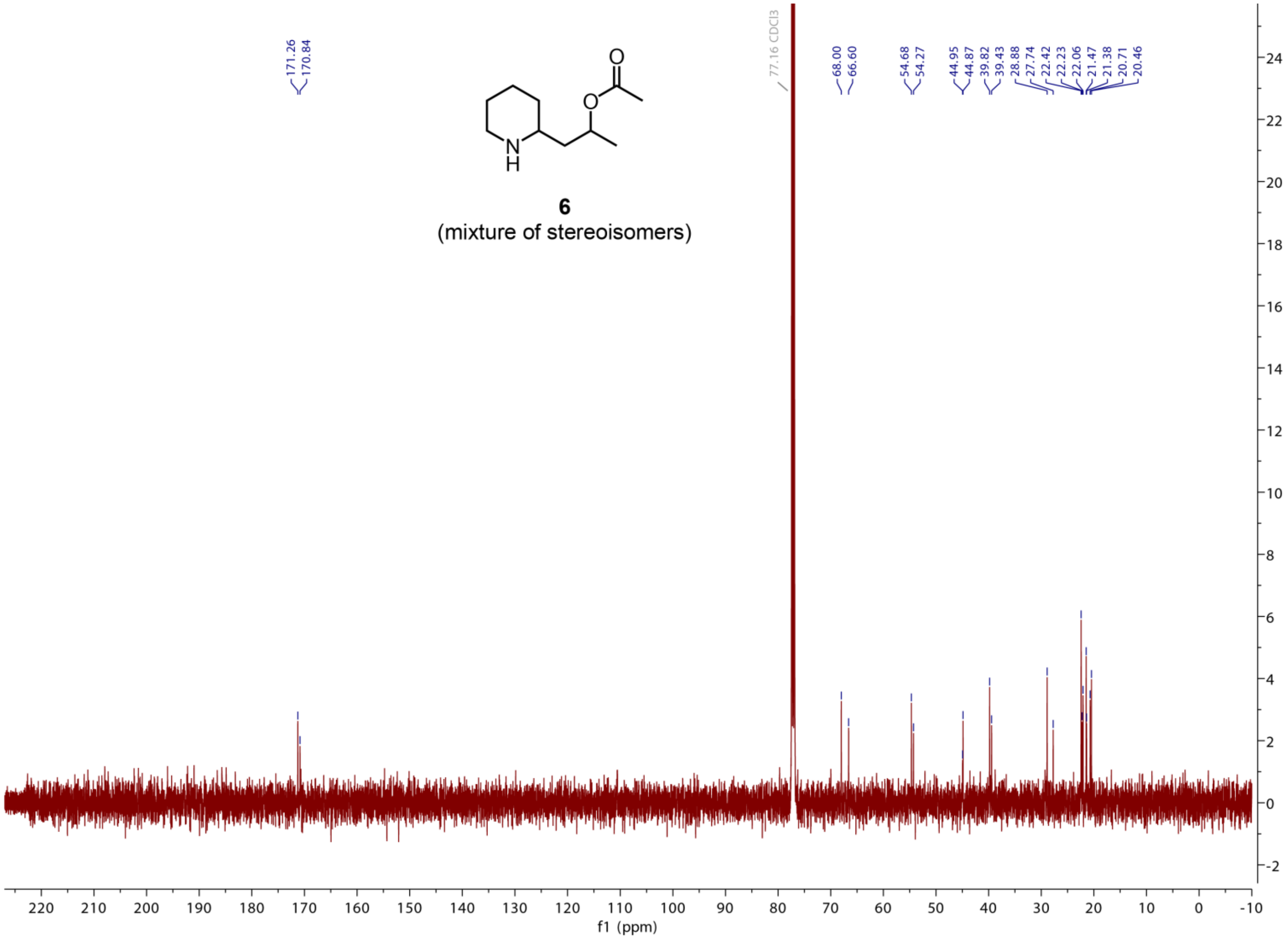

**Figure S17. $^{13}$C NMR spectrum from the synthesis of 6 stereoisomers.** This experiment was recorded at 500 MHz in deuterated chloroform ($CDCl_3$).

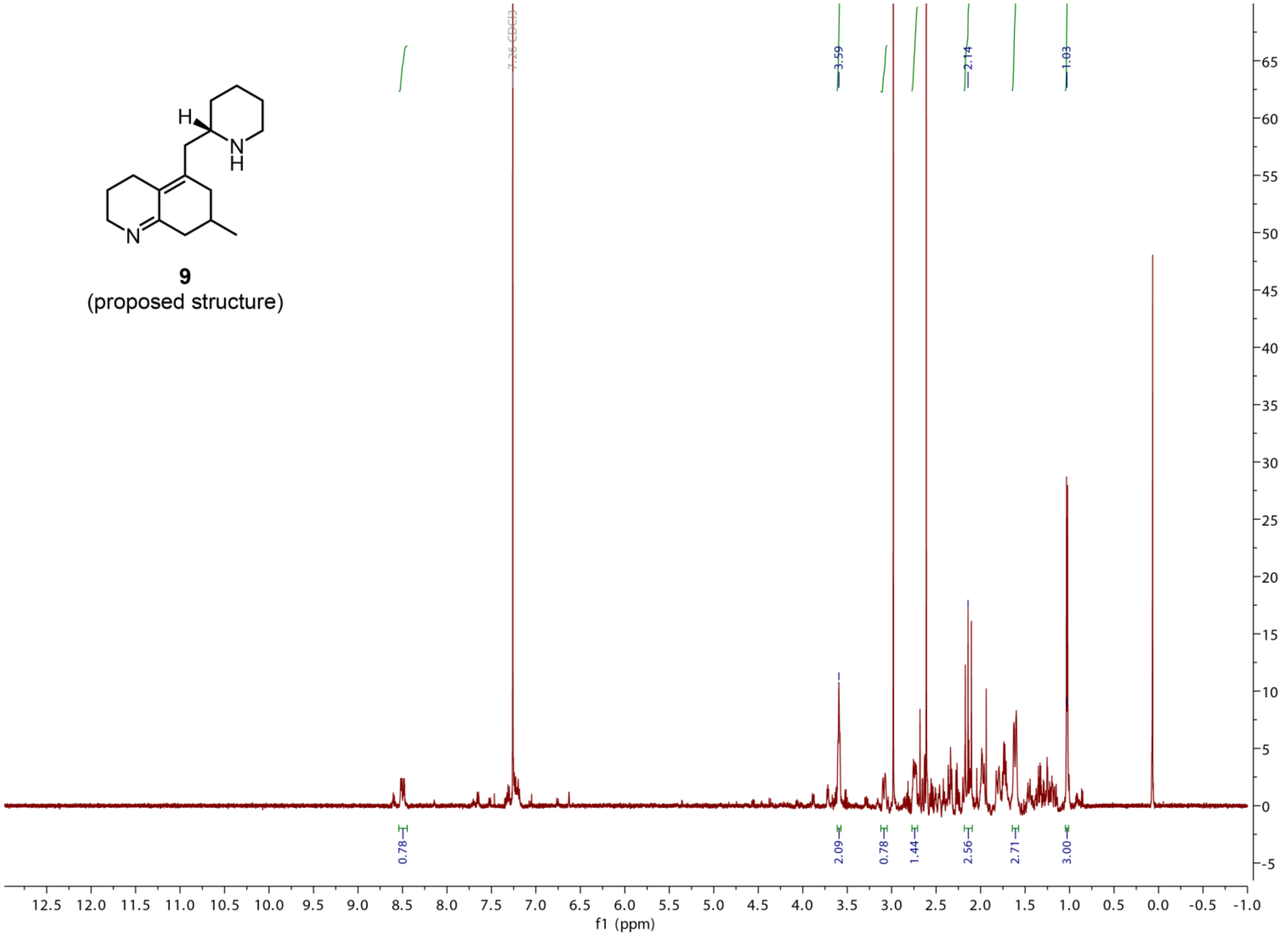

**Figure S18. 1H NMR (crude) of the purified product (putative 9, *m/z* 247.2169) of *Pt*CAL-1/*Pt*CAL-2.** This experiment was recorded at 500 MHz in deuterated chloroform ($CDCl_3$).

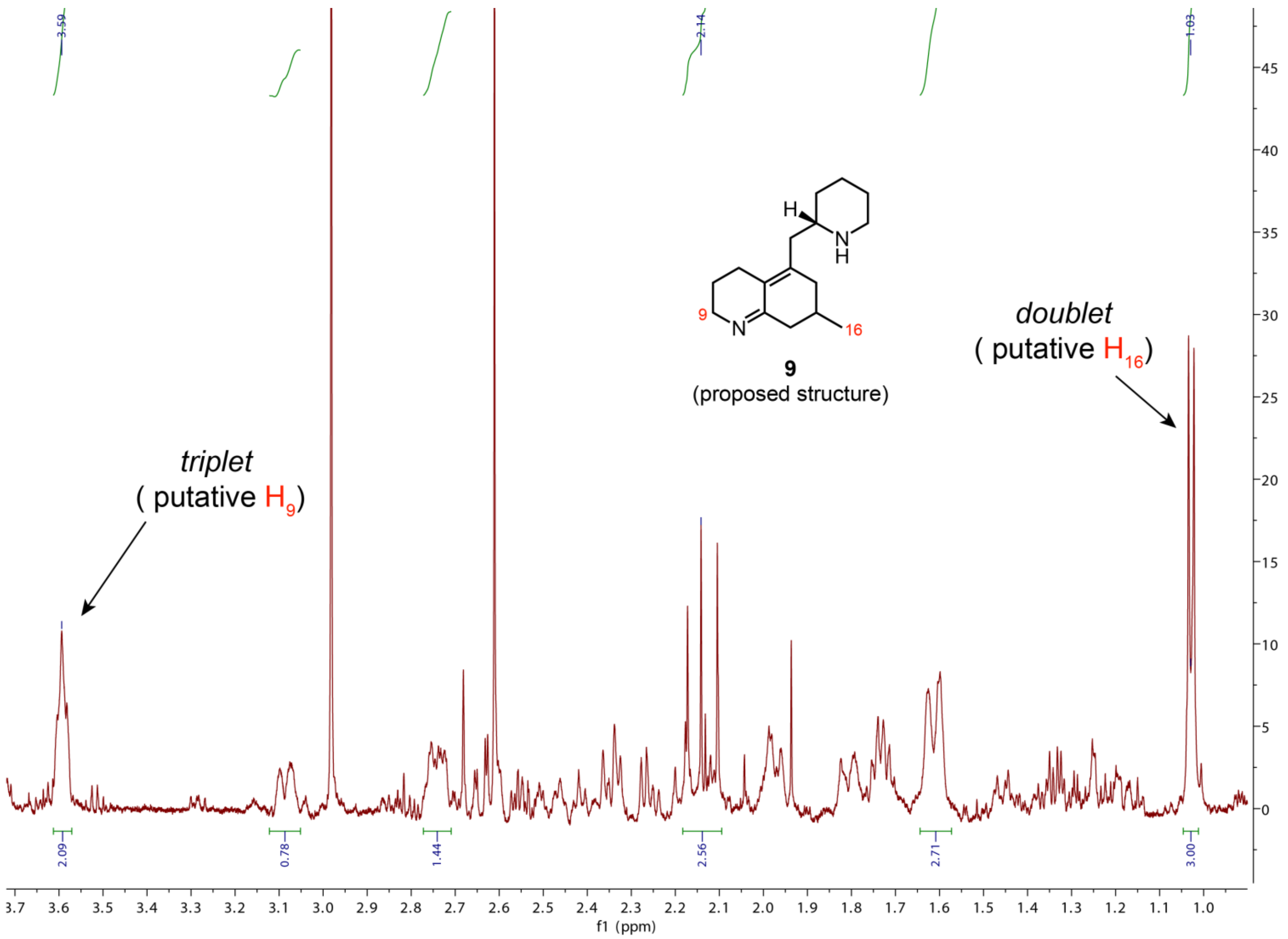

**Figure S19. [1]H NMR (crude) of putative 9 (zoomed in).** Zoomed in view from Figure S18.

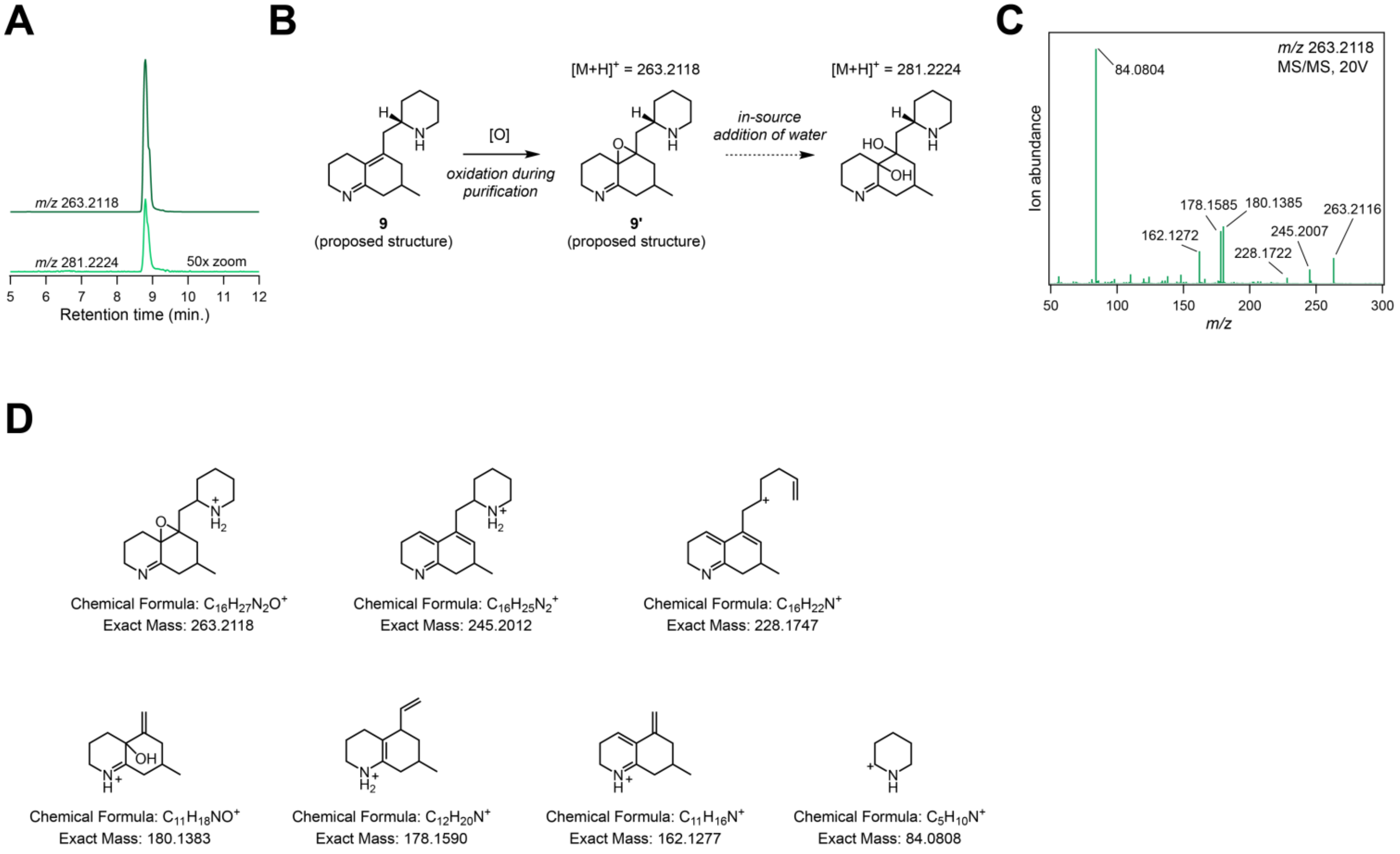

**Figure S20. LC-MS identification of a 9-related oxidized by-product (putative 9').** **A)** HILIC LC-MS analysis of a new major compound ($[M+H]^+$ = *m/z* 263.2118) purified while trying to isolate **9**. This compound corresponds to the addition of an oxygen, suggesting this to be an oxidized product of **9**. We also observed an in-source ion fragment that pertains to the addition of a water ($[M+H]^+$ = *m/z* 281.2224). **B)** Proposed oxidation of **9** to produce **9'**, which can undergo water addition during ionization. **C)** MS/MS spectrum (*m/z* 263.2118, 20V) of putative **9'**. **D)** Predicted structures of major MS/MS ion fragments.

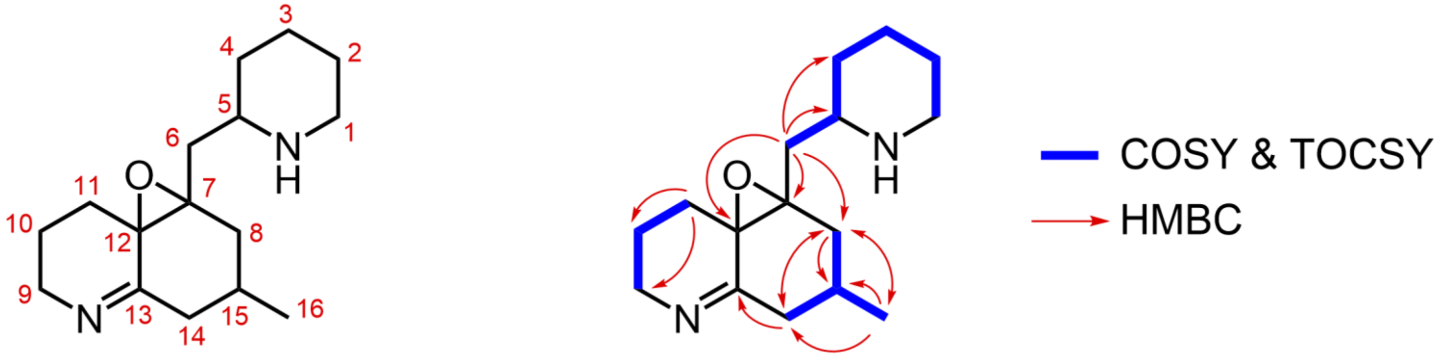

**9'**
(proposed)

**Figure S21. NMR assignment of the oxidized scaffold by-product 9' (*m/z* 263.2118).** Assignments, as listed in the table below, are based on the NMR spectra shown in Figures S22-S27. All experiments were recorded at 600 MHz in deuterated acetonitrile ($CD_3CN$).

| **carbon** | | **$^{13}$C δ (ppm)** | **$^{1}$H δ (ppm, *J* in Hz)** | |
|---|---|---|---|---|
| **1** | CH2 | 47.6 | 2.58 (1H, td, *J* =11.8, 2.8) | 2.97 (1H, dm, *J* = 12.0) |
| **2** | CH2 | 27.0 | 1.32b (1H, m) | 1.52 (1H, m) |
| **3** | CH2 | 25.7 | 1.32a (1H, m) | 1.74 (1H, m) |
| **4** | CH2 | 34.0 | 1.07 (1H, m) | 1.57 (1H, m, *J* = 5.6) |
| **5** | CH | 55.0 | 2.76 (dddd, *J* = 2.58, 5,46, 8.00, 10.59) | |
| **6** | CH2 | 39.9[a] | 1.56 (1H, dd, *J* = 5.6, 14,1) | 1.65 (1H, dd, *J* = 7.8, 14,1) |
| **7** | C | 66.8[a] | -- | |
| **8** | CH2 | 38.2 | 1.51 (1H, dd, *J* = 10.2, 14.5) | 2.14 (1H)[b] |
| **9** | CH2 | 50.1 | 3.39 (1H, m) | 3.78 (1H, dm, *J* = 17.9) |
| **10** | CH2 | 22.1 | 1.73 (1H, m) | 1.77 (1H, m) |
| **11** | CH2 | 26.4 | 1.62 (1H, m) | 1.88 (1H, td, *J* = 13.4, 3.9) |
| **12** | C | 57.6[a] | -- | |
| **13** | C | 167.2[a] | -- | |
| **14** | CH2 | 42.7 | 1.76 (1H, m) | 2.28 (1H, d, *J* = 13.8) |
| **15** | CH | 24.8 | 1.82 (1H, m) | |
| **16** | CH3 | 21.5 | 0.88 (3H, d, *J* = 6.43) | |

[a]Annotated based upon HMBC and/or HSQC experiments. [b]Hidden in the 1D proton spectrum.

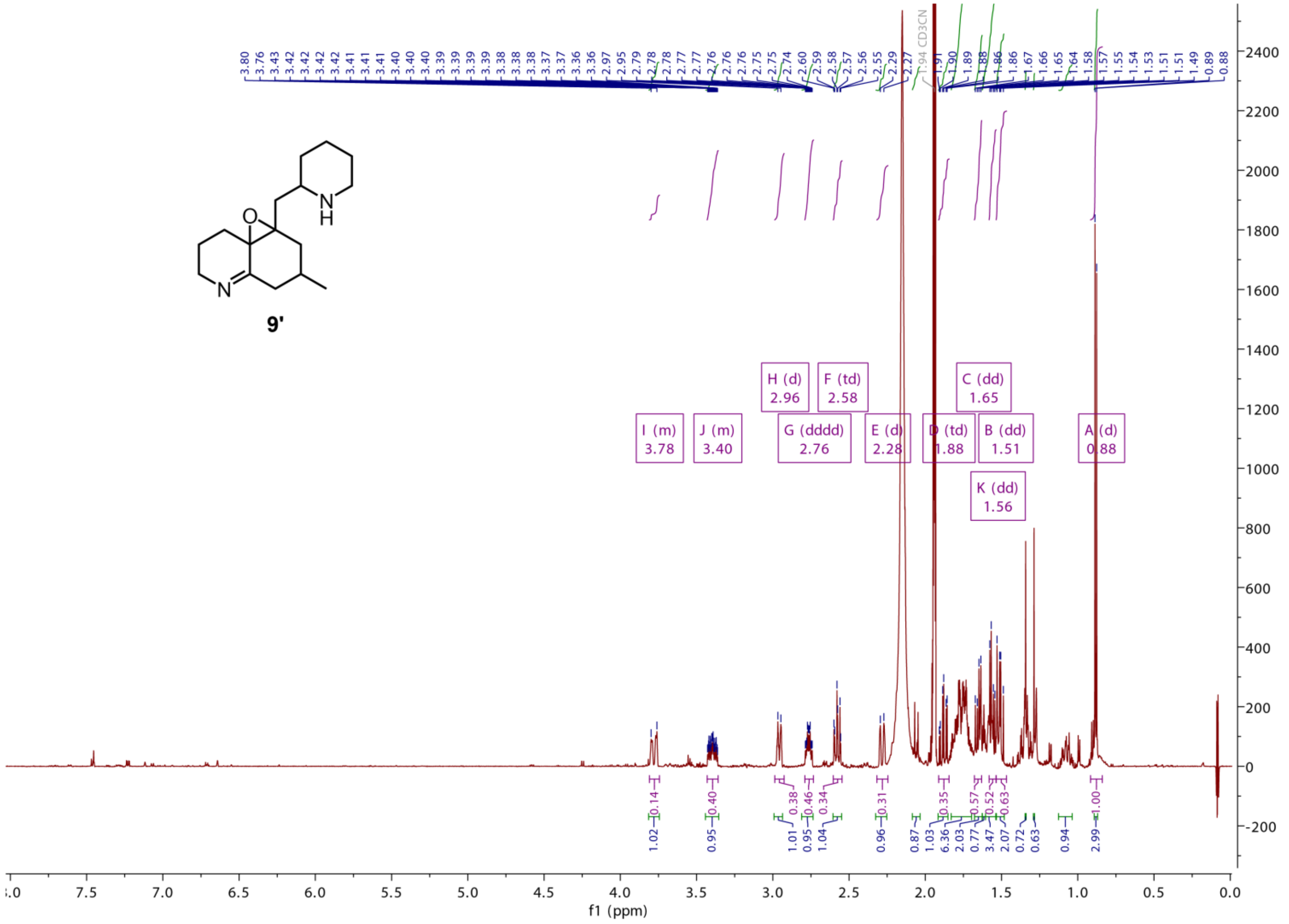

Figure S22. [1]H spectrum of the oxidized scaffold by-product 9’ (*m/z* 263.2118).

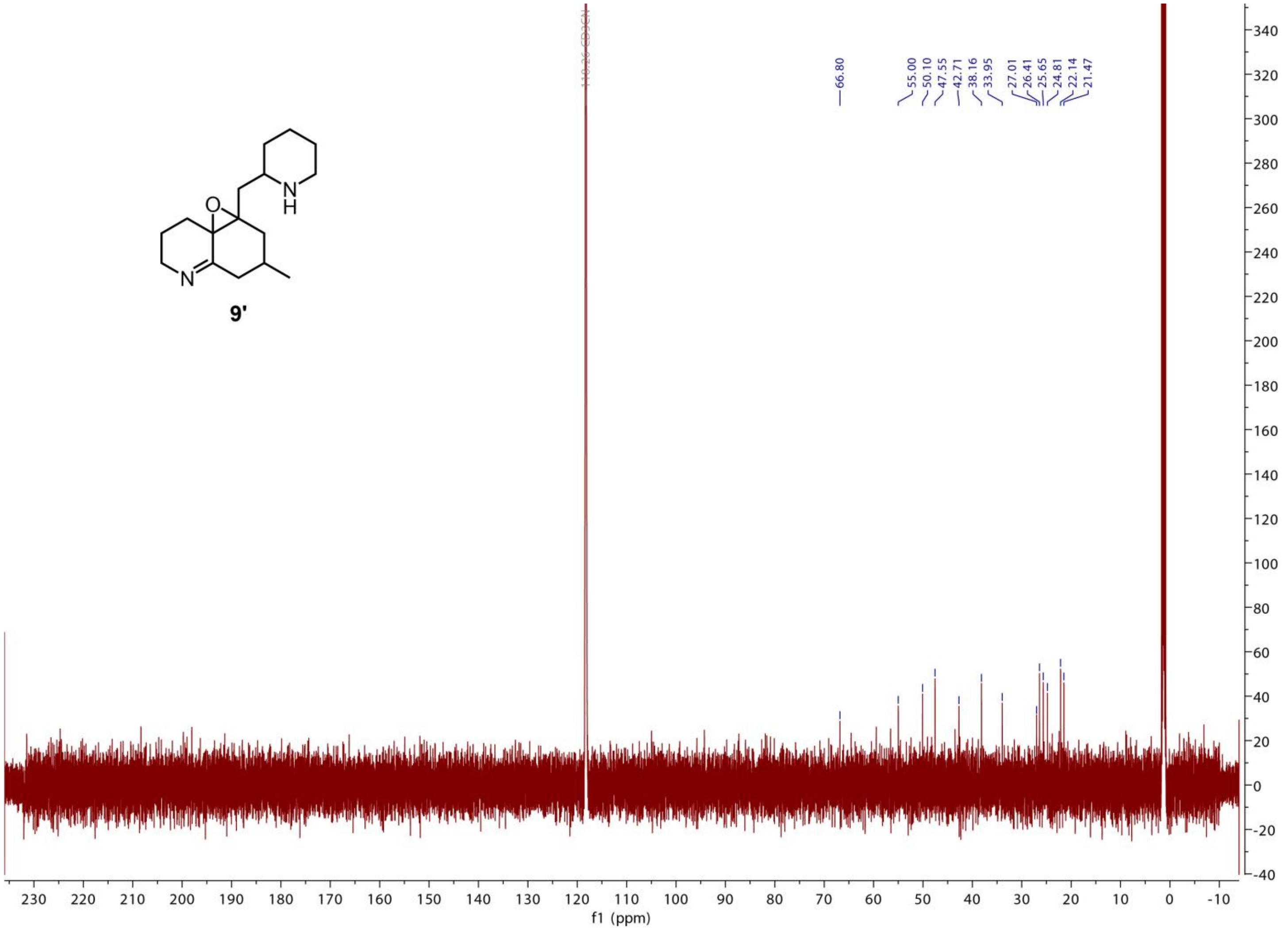

**Figure S23. $^{13}$C spectrum of the oxidized scaffold by-product 9’ (*m/z* 263.2118).**

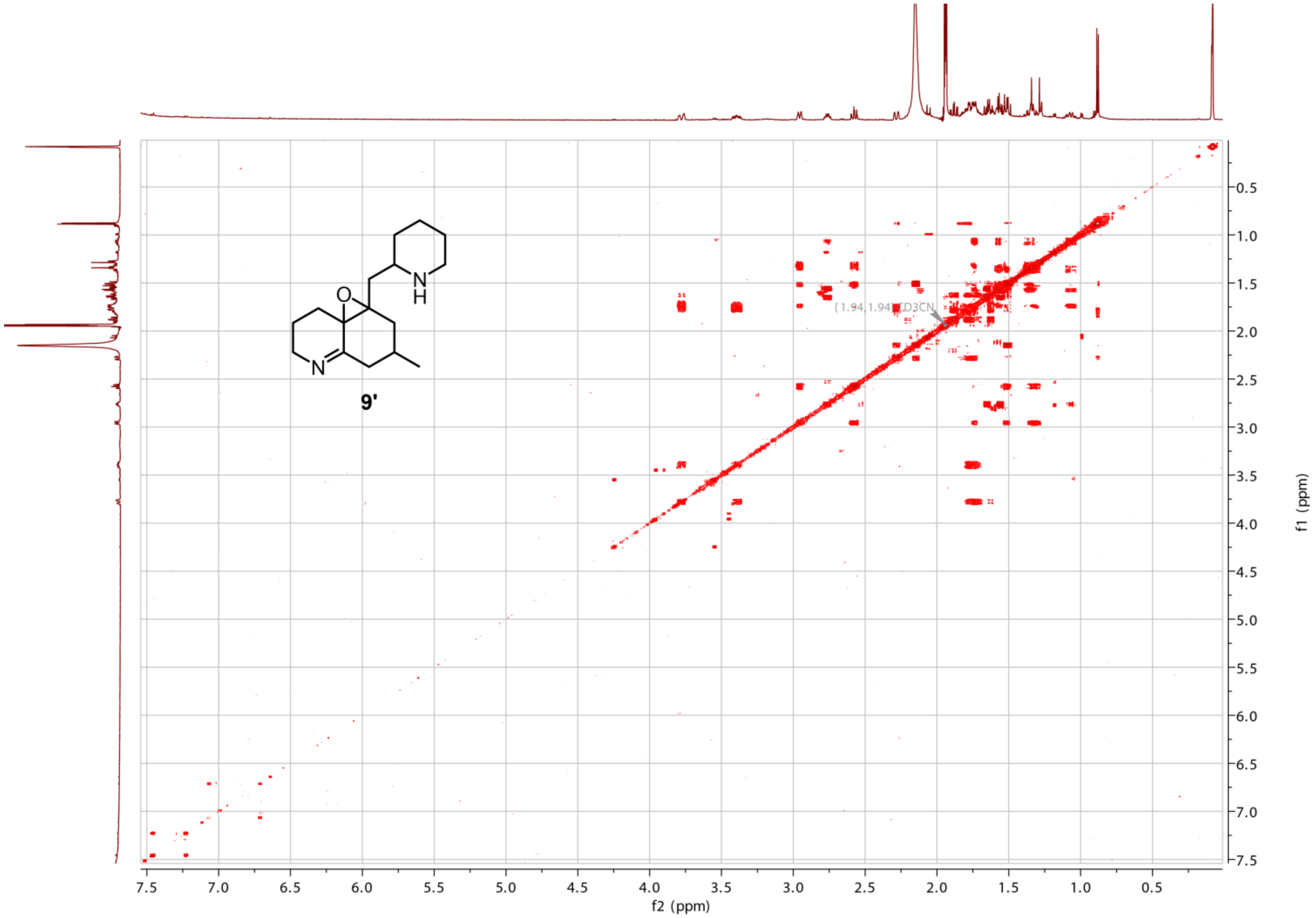

**Figure S24. COSY spectrum of the oxidized scaffold by-product 9’ (*m/z* 263.2118).**

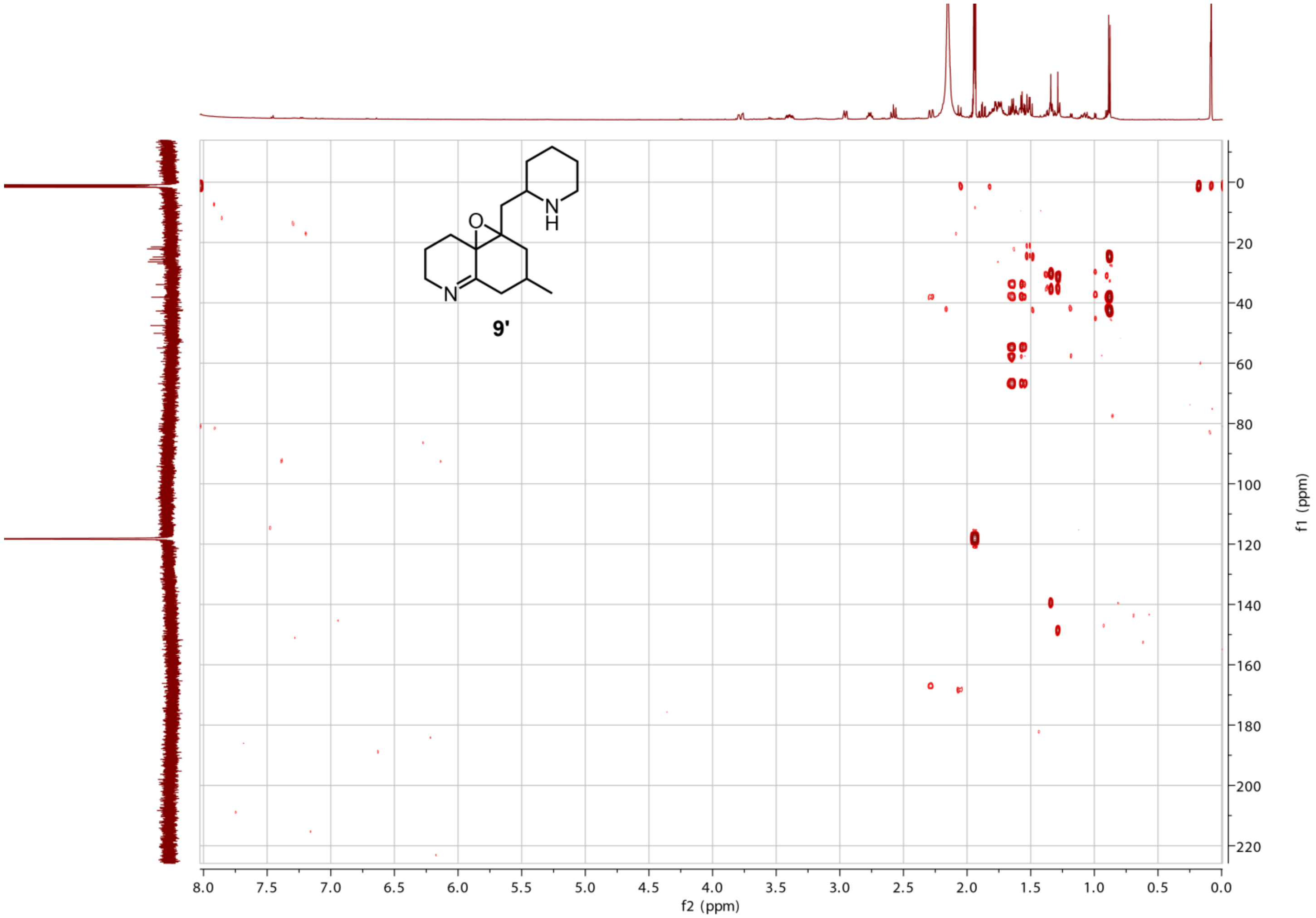

**Figure S25. HMBC spectrum of the oxidized scaffold by-product 9' (*m/z* 263.2118).**

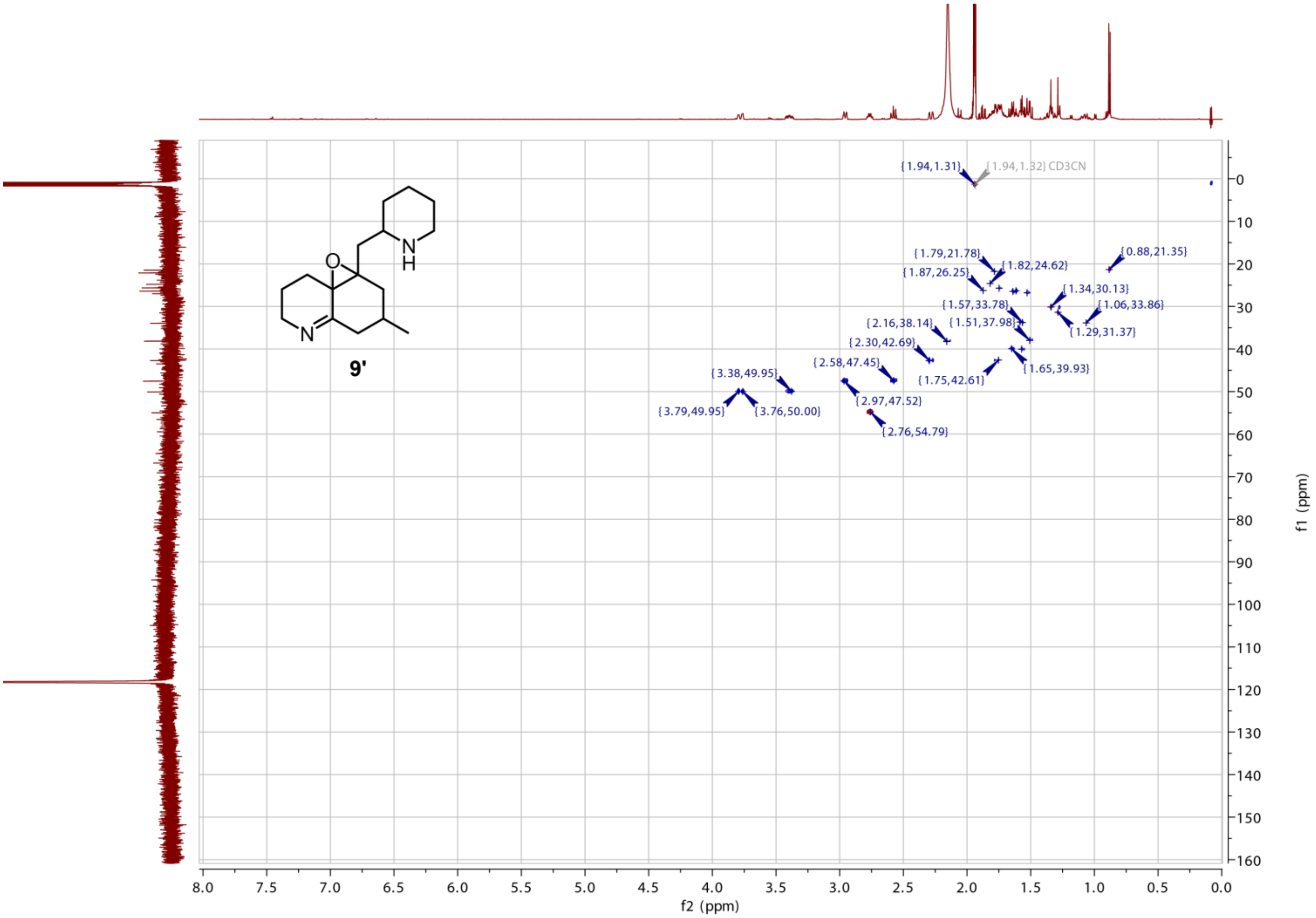

**Figure S26. HSQC spectrum of the oxidized scaffold by-product 9’ (*m/z* 263.2118).**

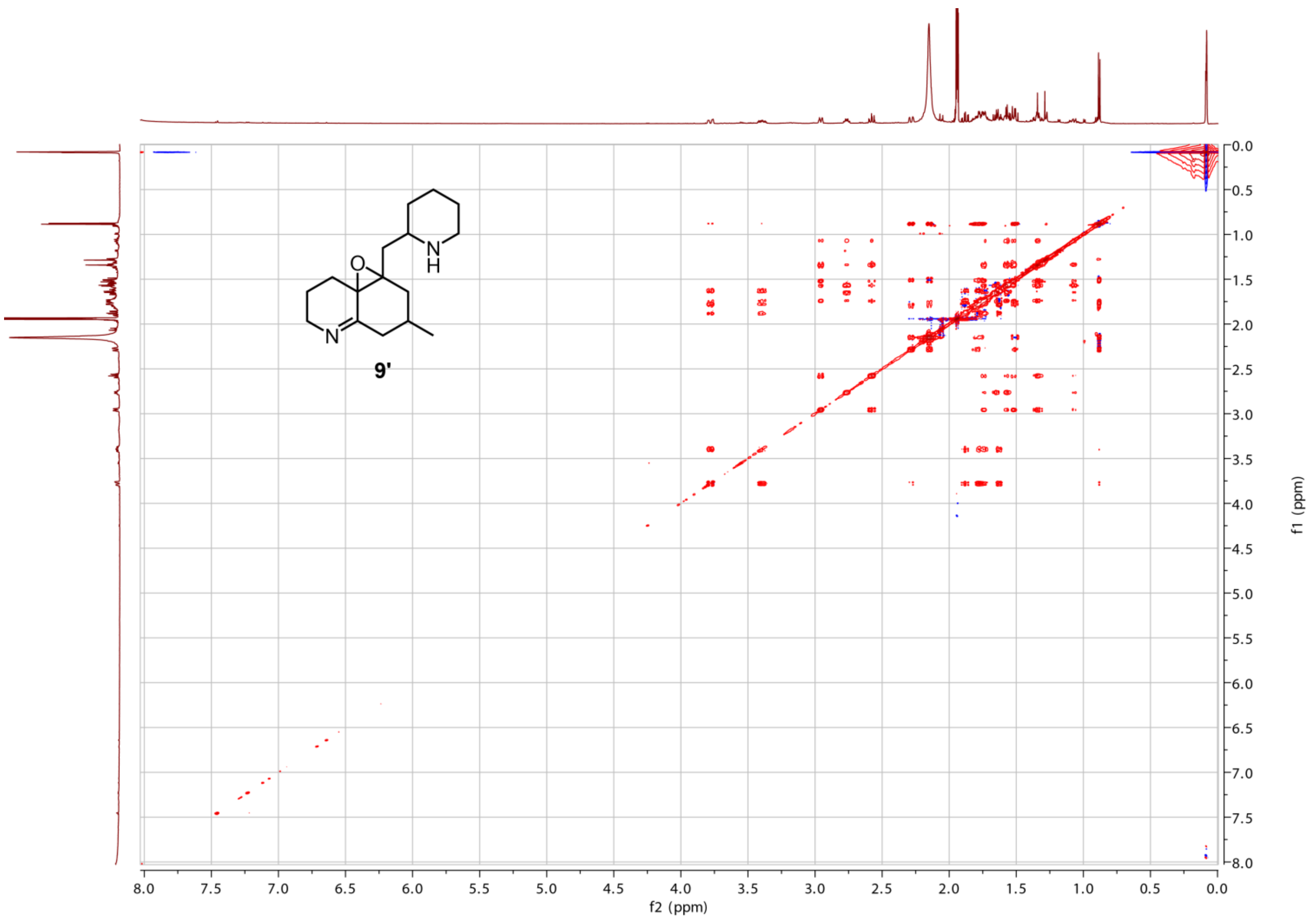

**Figure S27. TOCSY spectrum of the oxidized scaffold by-product 9’ (*m/z* 263.2118).**

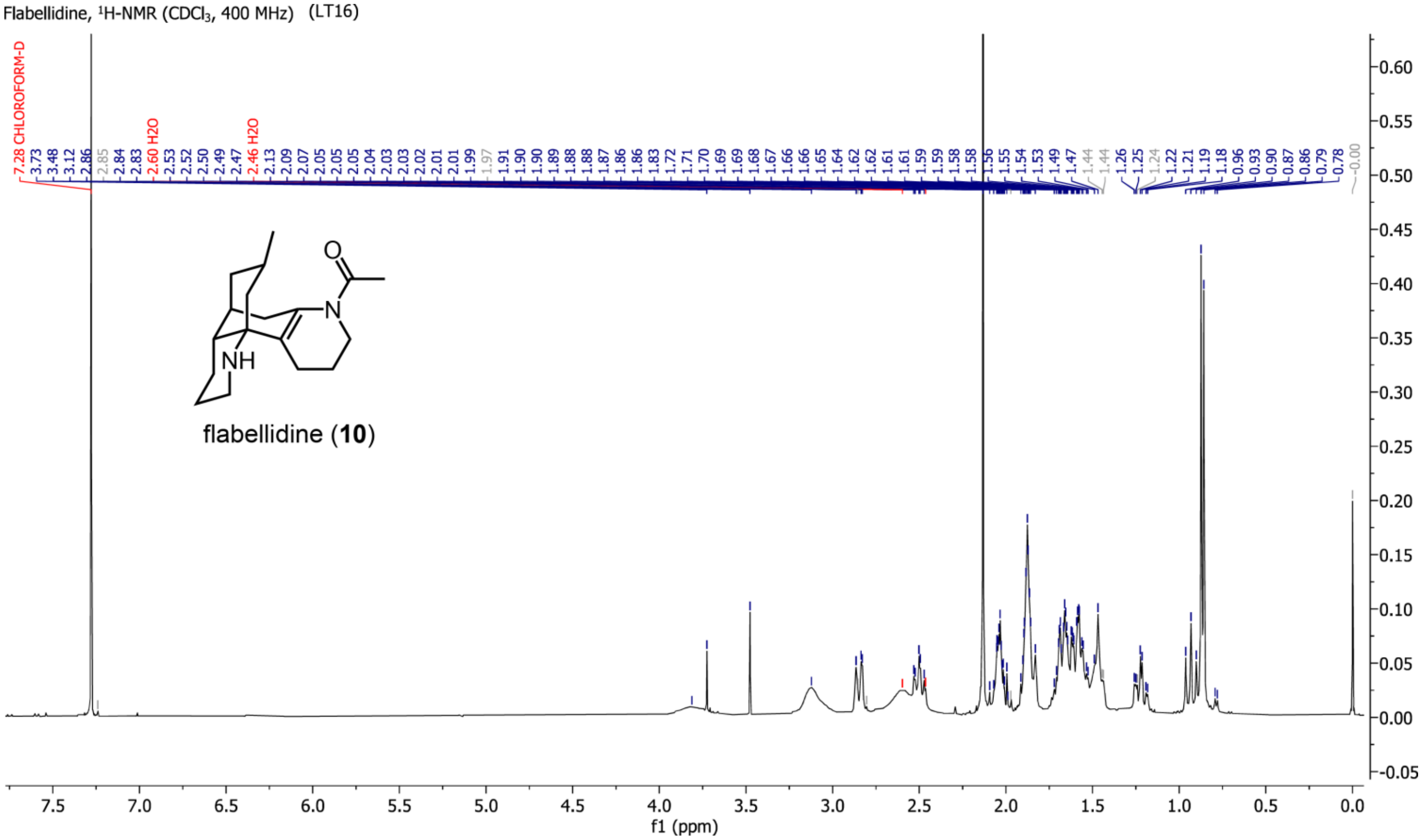

**Figure S28. $^1$H NMR spectrum ($CDCl_3$, 400 MHz) of flabellidine (10) isolated from *Lycopodium platyrhizoma*.(*1*)**

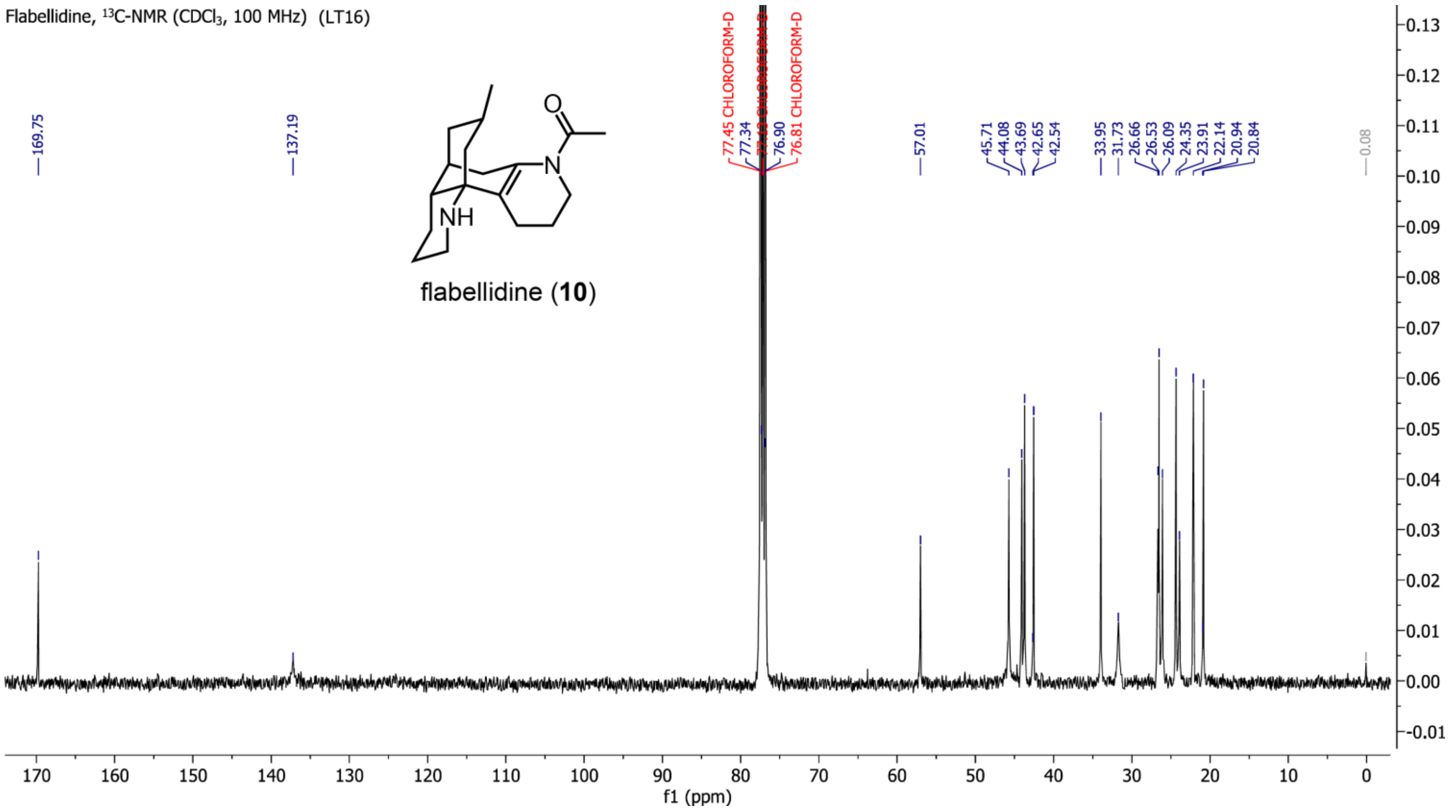

**Figure S29.** $^{13}$**C NMR spectrum (CDCl$_3$, 100 MHz) of flabellidine (10) isolated from *Lycopodium platyrhizoma*.**(*1*)

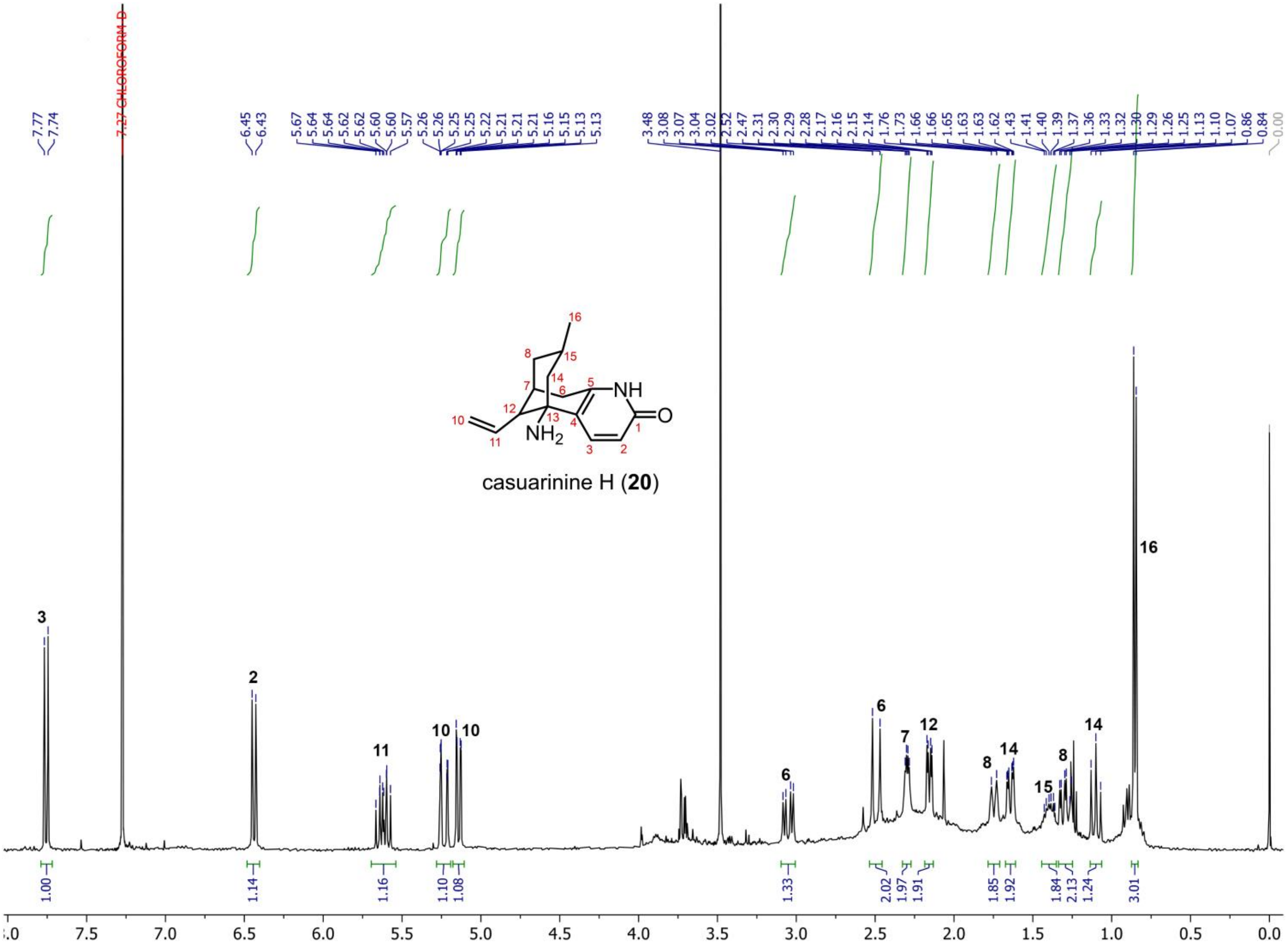

**Figure S30.** $^{1}$**H NMR spectrum (CDCl$_3$, 400 MHz) of casuarinine H (20) isolated from *Lycopodium platyrhizoma*.**(*1*)

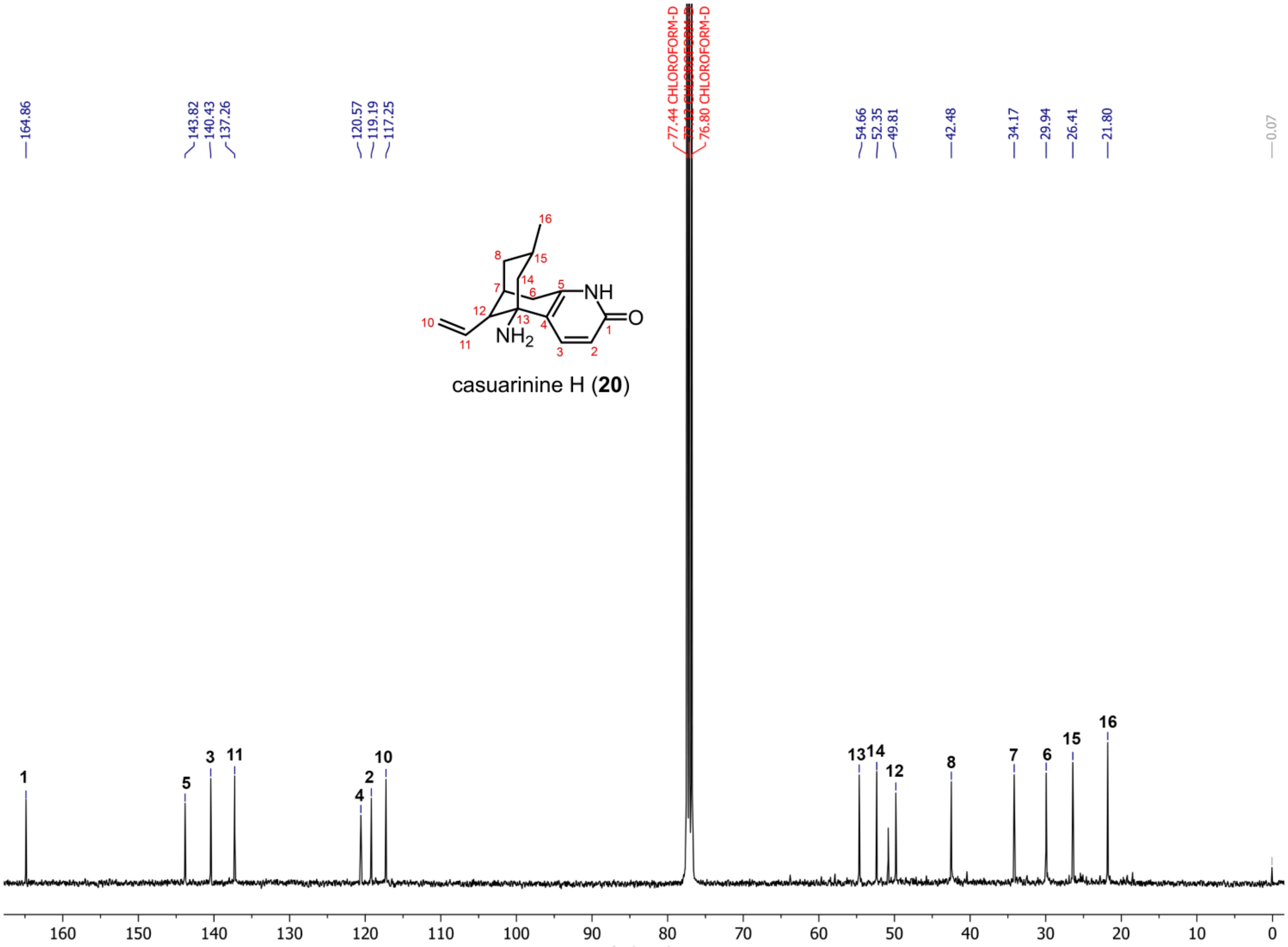

**Figure S31.** $^{13}$**C NMR spectrum (**$CDCl_3$**, 100 MHz) of casuarinine H (20) isolated from** ***Lycopodium platyrhizoma*****.**(*1*)
